# Stress-Associated Alterations in Amygdala-Striatal Activity: A Multi-Level Analysis of Distributional, Dynamical, and Computational Signatures

**DOI:** 10.64898/2025.11.29.691336

**Authors:** Feng Lin

## Abstract

Chronic stress is associated with alterations in neural circuit dynamics, yet the computational principles underlying these changes remain incompletely understood. Here, we reanalyzed in vivo GCaMP8s recordings from BLA-DMS and CeA-DMS projection pathways using three complementary descriptive approaches. First, Kullback-Leibler (KL) divergence was used to quantify stress-related changes in activity distributions, revealing context-dependent alterations in population-level neural responses. Second, a phenomenological second-order regression framework was employed to characterize local recovery-related dynamical features, providing a compact descriptive representation of pathway activity without implying mechanistic identification. Third, a simplified artificial neural network (ANN) was implemented as a hypothesis-generating sufficiency test. Under asymmetric optimization constraints, the model generated response profiles qualitatively resembling selected features observed experimentally. Robustness analyses indicated that similar qualitative behaviors emerged across multiple parameter initializations. Together, these results provide a multi-level descriptive characterization of stress-associated neural activity alterations across statistical, dynamical, and computational representations, while explicitly avoiding assumptions of a unified mechanistic mapping between analytical levels.

## 1 Introduction

Establishing a precise mapping between psychological states and their underlying neural substrates remains a central tenet of cognitive neuroscience [1, chap. 1, p. 1-17]. Psychological processes arise from the dynamics of specific neural circuits, and external behavioral patterns therefore serve as observable read-outs of internal neural computations[2]. Quantitative analyses of neural activity associated with behavior can therefore provide useful insights into how neural circuit dynamics vary across psychological states [3]. Furthermore, this mind-brain-behavior relationship is inherently dynamic: environmental factors and constraints often drive shifts in psychological states, triggering a modulation of neural circuits that results in adaptive behavioral changes.

This dynamic mapping is particularly evident in the context of stress, a potent environmental constraint that precipitates significant shifts in both psychological states and behavioral strategies [4]. Chronic stress induces a transition from flexible goal-directed behavior to rigid habitual responding [5]. While classical models of habit formation emphasize the functional transition from the goal-directed dorsomedial striatum (DMS) to the habit-mediating dorsolateral striatum (DLS) [6], recent evidence suggests that chronic stress is associated with shifts in the relative contribution of amygdala inputs to DMS circuit activity [5]. Specifically, Giovanniello et al. [5] demonstrated that chronic stress differentially modulates two amygdala projections to the DMS. Reward-related activity in the BLA-DMS pathway, which has been implicated in action-outcome learning, is suppressed by stress, whereas activity in the CeA-DMS pathway becomes progressively engaged during learning and is associated with the emergence of inflexible habit-like behavior. These findings motivate a model-based hypothesis in which stress may lead to a context-dependent reinterpretation of reward-related neural signals. Rather than reflecting a fixed mapping between reward signals and behavioral outputs, the functional significance of reward-related activity may depend on the environmental context. Under chronic stress, the behavioral significance of these reward-related signals may differ depending on circuit context.

In the experimental paradigm of Giovanniello et al. [5], mice were first assigned to control or stress groups (8-12 mice per group). The stress group underwent 14 days of chronic mild unpredictable stress, consisting of pseudorandom exposure to several mild stressors, including footshock, restraint, white noise, tilted cages, damp bedding, and continuous illumination. After the stress period, mice were trained in an operant lever-press task to obtain food rewards across multiple reinforcement schedules. Following training, outcome-based behavioral tests were performed to assess the balance between goal-directed and habitual control. Throughout the experiment, GCaMP8s fiber photometry was used to record calcium signals in the BLA-DMS and CeA-DMS projections.

Beyond experimental approaches, computational modeling has become an important tool for characterizing neural dynamics and calcium signaling processes. Mathematical and machine learning models provide flexible frameworks for describing nonlinear patterns in neural activity and generating testable hypotheses regarding neuronal responses under varying conditions[7, 8]. In this context, connectionist frameworks provide a heuristic tool to test whether discrete experimental observations can emerge from continuous, optimization-driven computational representations[7]. Collectively, these approaches have contributed to the quantitative analysis and interpretation of complex neural systems [9].

From a dynamical systems perspective, neural activity can be characterized in terms of local stability properties and trajectory evolution within an effective state space. Because the available recordings are limited to perievent windows, we focus specifically on local stability-related features that can be estimated from short temporal segments. Chronic stress may influence this balance, potentially biasing neural activity toward more rigid dynamical regimes associated with habitual behavior.

Artificial neural networks provide a flexible computational framework for testing whether simple constraints can generate complex activity patterns[10, 11, 12, 13]. In the present study, the ANN framework is used exclusively as a hypothesis-generating and sufficiency-testing tool rather than as a biologically realistic model of neural processing[9].

Because the BLA-DMS and CeA-DMS recordings were obtained from separate animals rather than simultaneous recordings, the regression framework is not intended to infer real-time interactions between pathways. Instead, it provides a phenomenological description of population-level statistical relationships among pathway-specific activity summaries under common experimental conditions.

Despite demonstrating a quantitative association between chronic stress, pathway signal alterations, and behavioral rigidity, the computational principles that may account for these relationships remain unclear. Traditional methods, such as direct signal observation and behavioral analysis, do not fully capture the complexity of the observed neural activity patterns. To address this gap, we analyze the publicly available GCaMP8s calcium fluorescence time-series dataset reported by Giovanniello et al. [5]. Using these data, we develop a multi-level descriptive framework designed to examine stress-related activity changes from complementary analytical perspectives, including distributional structure, local recovery dynamics, and constraint-based generative modeling. This approach is designed to evaluate whether chronic stress is associated with persistent shifts in the distributional structure of neural activity (potentially consistent with the aforementioned hysteresis-like signatures) and to compare how these alterations appear under different descriptive representations.

Because stress-related alterations may manifest simultaneously at multiple descriptive levels, no single analytical representation is sufficient to characterize the observed activity patterns. We therefore examine the data from three complementary perspectives: distributional structure, local recovery dynamics, and constraint-based generative modeling. These perspectives are not intended to form a unified formal theory; rather, they provide distinct descriptive views of the same experimental observations. Nevertheless, if independent descriptive approaches reveal qualitatively consistent stress-dependent trends, such consistency may provide useful guidance for subsequent hypothesis generation.

To gain a multi-level insight into these stress-induced alterations, we do not restrict our analysis to a single statistical metric; instead, we deploy a tripartite descriptive framework spanning complementary analytical resolutions. First, we utilize information-theoretic measures (Kullback–Leibler divergence) to map the macroscopic shifts in population activity distributions to quantify condition-dependent changes in population activity distributions. Second, we construct a phenomenological second-order regression model to characterize local recovery dynamics and stability-related trajectory features that accompany the observed distributional changes. Under this framework, pathway-specific summaries obtained from different animals are treated as condition-level observations that may share common statistical regularities. Finally, we implement an artificial neural network (ANN) as a computational sufficiency-testing framework to examine whether a minimal set of asymmetric local constraints is capable of reproducing selected qualitative signatures observed experimentally. The ANN is not intended to predict neural activity or validate biological mechanisms, but rather to evaluate whether the proposed constraints are sufficient to generate the observed phenomenological patterns.

This non-mechanistic, cross-scale approach allows us to compare whether different descriptive levels reveal qualitatively compatible patterns of stress-related neural adaptation. Together, these analyses provide complementary descriptive perspectives on stress-related neural adaptation across multiple analytical scales. Each analytical level addresses a different question: distributional analysis characterizes what changes under stress, dynamical modeling characterizes how these changes evolve locally in time, and ANN-based sufficiency testing examines which minimal constraints can reproduce the resulting phenomenological signatures.

## 2 Method

### 2.1 Data Source and Event Classification

We performed a secondary analysis on GCaMP8s fiber photometry recordings (sampling rate: 20 Hz) originally reported by Giovanniello et al. [5]. Signals were recorded from BLA–DMS and CeA–DMS projections in mice under two experimental conditions: Control and Chronic Stress (14 days). For each behavioral event, we extracted *±*10 s time windows to characterize peri-event neural activity.

For analytical purposes, behavioral events were grouped into two categories according to whether they were contingent on learned instrumental behavior:

1. Non-Learned Modality (Non-Contingent Stimuli): These events occur independently of the mouse’s actions. The primary event in this category is the aversive footshock stimulus delivered outside the instrumental task context. Because it is not contingent on ongoing behavior, this stimulus provides an externally imposed perturbation for examining neural responses under stress conditions.
2. Learned Modality (Contingent Events): This modality includes task-related events during the operant lever-press paradigm in which mice learned to associate lever pressing with food rewards. Task difficulty was manipulated using Random Ratio (RR) schedules, in which reward probability gradually decreased from 1.0 (FR1) to 0.1 (RR10). These schedules introduce increasing outcome uncertainty during learning. In addition, unpredicted rewards were occasionally delivered during the task. Although these rewards were not contingent on the mouse’s immediate action, they occurred within a task context in which mice had already learned an association between lever pressing and reward delivery.

In the present study, footshock events from the non-learned modality were used as the primary dataset for dynamical analysis. Because these stimuli occur independently of instrumental behavior, they provide a relatively controlled perturbation for examining stress-related changes in neural activity. Data from the learned modality were subsequently analyzed as a complementary comparison dataset to examine whether similar statistical and dynamical patterns were observable in task-related neural activity.

### 2.2 Data analysis

#### 2.2.1 Statistical Perspective: Measuring Condition-dependent Changes in Population Activity Distributions via KL Divergence

To quantify condition-dependent changes in population activity distributions following perturbations, we measured the extent to which post-stimulus activity distributions remained distinguishable from their pre-stimulus baseline distributions over time. Because recovery of the mean signal does not necessarily imply recovery of the full distribution, we focused on distributional persistence at the population level.

The baseline distribution was estimated from pooled pre-stimulus samples, and time-resolved post-stimulus distributions were reconstructed using a sliding-window procedure. The Kullback–Leibler (KL) divergence (*D_KL_*) was then used to quantify the difference between each post-stimulus distribution and the corresponding baseline distribution.

In this study, KL divergence is used purely as a descriptive statistical measure of distributional deviation. It is not interpreted as a direct indicator of latent neural states, attractor transitions, or specific dynamical mechanisms. Rather, sustained KL divergence indicates that the statistical structure of neural activity remains distinguishable from baseline activity over the observation period.

We reconstructed the PDFs (*p_baseline_* and *p_t_*) of GCaMP8s fluorescence signals using Gaussian Kernel Density Estimation (KDE). A sliding window approach (window size = 1 s, step = 0.05 s) was used to estimate time-resolved distributions, pooling samples across trials within each window. The Kullback–Leibler (KL) divergence is defined as:

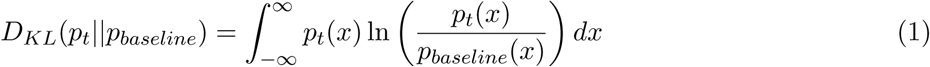

where *x* represents normalized signal intensity. In addition to distributional differences captured by KL divergence, changes in lower-order statistics such as mean and variance are reported in the Supplementary Materials (see Supplementary Materials Table D8 for empirical values). We define an integrated metric *I* as the temporal sum of KL divergence over the recovery period:

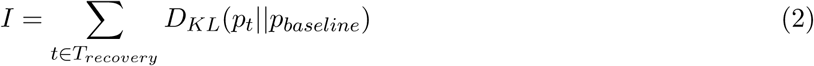

Larger values of *I* indicate greater cumulative divergence of the signal distribution from baseline over time. *I* is used as a descriptive statistical summary of these differences.

To ensure robustness against sampling variability and parameter selection, two validation procedures were implemented:

- Automated Bandwidth Selection: We employed Scott’s Rule to determine the KDE bandwidth, reducing subjective bias in PDF reconstruction. Scott’s Rule adaptively determines the optimal bandwidth based on the data’s variance and sample size, ensuring that the PDF faithfully represents the underlying stochastic structure without over-smoothing or introducing noise-driven artifacts. This selection is consistent with the smoothed inference framework proposed by McNealis et al. 2026 [14], which suggests that data-driven bandwidths optimize the mean-squared error and ensure the robustness of uncertainty quantification in finite-population contexts.
- Subject-level Bootstrap: We reported 95% confidence intervals (CIs) derived from 1000 bootstrap iterations. By resampling at the subject level, we ensure that the observed distributional differences represent a robust group-level phenomenon rather than an artifact of individual outliers.

This statistical analysis was conducted independently of the subsequent dynamical modeling, which was performed directly on the time-series data. The two analytical approaches therefore provide complementary descriptions of the recorded neural activity at different levels of analysis.

While the KL-divergence analysis characterizes distributional properties of the recorded signals, the dynamical modeling characterizes local dynamical features inferred from the same time-series data. No formal equivalence between these analytical layers is assumed. Although this analysis does not provide information about local stability properties, it provides a complementary statistical description that can be compared with the subsequent stability-related characterization of the same recordings.

#### 2.2.2 Dynamical Perspective

We adopted a second-order phenomenological dynamical model to provide a compact description of recovery trajectories observed in calcium activity. The model is intended to characterize effective dynamical structure in the observed signals rather than to recover underlying biophysical mechanisms. Although calcium signals are often approximated by first-order relaxation processes, projections of high-dimensional neural population dynamics onto low-dimensional observables can exhibit effective inertial dynamics such as delayed recovery or overshoot. A second-order phenomenological expansion therefore provides a minimal representation capable of capturing both state-dependent recovery tendencies and velocity-dependent relaxation effects. Cross terms are retained to allow interactions between state magnitude and recovery velocity. Such effective second-order dynamics can emerge when high-dimensional neural activity is projected onto low-dimensional observables. In addition, calcium indicators introduce intrinsic temporal filtering, which can effectively generate higher-order temporal dynamics in the observed signals.

##### Single-Variable Dynamical Modeling and Quasi-potential Analysis

To characterize local recovery dynamics in neural activity, we modeled the BLA–DMS and CeA–DMS pathways using phenomenological autonomous dynamical systems reconstructed from the observed calcium trajectories. For each pathway, a phase space was constructed using the signal amplitude (*x_BLA_* or *x_CeA_*) and its first derivative (*̇x_BLA_* or *̇x_CeA_*), representing the instantaneous state and rate of change of neural activity.

To ensure robustness against individual variability, we implemented a subject-level bootstrap procedure. For each of *n* = 1000 bootstrap iterations, subjects were resampled with replacement and the governing differential equation was estimated using Ridge Regression. From these fitted models, we derived the distribution of Jacobian eigenvalues (*E_i_, i* = 1, 2), where the real part of the Jacobian eigenvalues provides a measure of local dynamical stability, with more negative values indicating faster recovery toward the reconstructed steady state. Stability estimates were reported as 95% confidence intervals (CIs) derived from the bootstrap distribution. Further details on the regression formulations and parameter estimation can be found in Supplementary Section A.

To complement local stability analysis, we constructed an effective quasi-potential representation *U* (*x*) and phase portraits to visualize the local topology of the reconstructed phase space. This quasi-potential representation was obtained from the dominant state-dependent restoring component of the fitted dynamics and should be interpreted as a geometric visualization rather than a physical energy function. Because the system exhibits a single stable equilibrium, the curvature of the effective quasi-potential near the fixed point provides a geometric interpretation of the local stability quantified by the Jacobian eigenvalues. This formulation is conceptually related to classical attractor-based descriptions of neural dynamics, where neural states evolve within an energy landscape toward stable fixed points[15]. By integrating the dominant state-dependent restoring component of the fitted dynamics, we obtained an effective quasi-potential representation for each pathway, where the local curvature near equilibrium provides a geometric description of the reconstructed recovery dynamics.

Steady states were identified as stable fixed points of the reconstructed dynamical system satisfying *̇x* = 0 and *̈x*= 0. This representation illustrates how chronic stress is associated with changes in the reconstructed phase space local topology, producing a broader reconstructed quasi-potential profile and altered geometric curvature in the local phase-space representation.

Given the consistency of these dynamical features across perturbation types, we focused the quasi-potential and phase-portrait analyses on representative non-learned (footshock) and learned (unpredicted reward) perturbations. These perturbations provided illustrative examples of how differences in local dynamical stability were associated with changes in the reconstructed phase space local topology.

Finally, to compare statistical and dynamical characterizations of stress-related activity patterns, we examined the relationship between steady-state KL divergence and the principal Jacobian eigenvalue. Steady state was defined as the KL divergence between the pre-perturbation baseline distribution and the neural population distribution during the final 5 seconds of the recovery period. Because KL divergence and Jacobian eigenvalues provide complementary statistical and dynamical descriptions of recovery, respectively, we examined whether they exhibit consistent condition-dependent trends. Any observed correlation should be interpreted as a coarse-grained statistical reflection of the reconstructed dynamical structure rather than evidence of a rigorous theoretical relationship.

The resulting correlation was interpreted as an empirical association between statistical and dynamical characterizations of recovery rather than a formal theoretical mapping between the two quantities. Establishing such a mapping would require additional assumptions regarding stochastic dynamics, stationarity, and ergodicity that cannot be directly evaluated using the present dataset. Even in the absence of a formal mapping, KL divergence may still provide a coarse statistical reflection of differences in local dynamical stability, analogous to a low-resolution projection of the underlying attractor structure.

Accordingly, the correlation analysis was used to assess cross-scale consistency between statistical and dynamical descriptions of recovery, rather than to demonstrate that KL divergence directly reflects dynamical stability.

##### Population-Level Dynamical Association Modeling

To examine whether chronic stress alters population-level relationships between pathways, we constructed a phenomenological cross-pathway association model linking BLA-DMS and CeA-DMS activity. Although recordings from the two pathways were obtained in separate experimental sessions, we assumed that pathway-level activity patterns within a given experimental condition share statistically consistent dynamical regularities across animals. This assumption allows population-level pooling to recover shared statistical structure and condition-dependent dynamical regularities across animals [16]. This approach estimates cohort-level statistical associations rather than moment-to-moment causal interactions between simultaneously recorded neurons. Consequently, this population-level dynamical association model should be understood as a shorthand for statistical association within the reconstructed phenomenological model and not as evidence of real-time neural coupling, information transfer, or causal interaction.

To reconstruct these dynamical regularities across animals, we extended the phase space to four dimensions: *{x_BLA_*, *̇x_BLA_*, *x_CeA_*, *̇x_CeA_}*. We employed a multivariate Ridge Regression to model the acceleration of a target pathway that chosen for its ability to penalize non-robust terms and handle multi-collinearity:

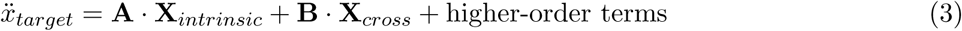

where **X***_intrinsic_* and **X***_cross_*represent the state vectors |*x, {x*} of the target and partner pathways, respectively. Because the model is phenomenological and estimated using regularized regression, individual coefficients are not interpreted as direct physical parameters. Instead, the fitted terms are used to characterize how pathway states are associated with changes in the reconstructed dynamics. The explicit polynomial form of the regression model is provided in the Supplementary Materials Section B.

Parameter reliability was evaluated using a subject-level bootstrap procedure (*n* = 1000). Cross-pathway association terms were considered significant when the 95% confidence interval of the corresponding cross-pathway coefficient excluded zero. Because the two pathways were not recorded simultaneously, the estimated association terms should not be interpreted as real-time coupling. Instead, they describe reproducible population-level coordination patterns observed across animals exposed to the same experimental condition.

##### Comparative Analysis Across Perturbation Contexts

To determine whether chronic stress is associated with systematic changes in the reconstructed dynamical signatures of neural activity, we categorized experimental perturbations into two modalities: (1) Non-Learned Modality (Unpredicted Footshock) probes the circuit’s intrinsic response and relaxation characteristics from high-arousal, un-modeled perturbations; (2) Learned Modality (Unpredicted Reward and Operant Actions) examines the circuit’s adaptive stability within a learned operant context. Using Jacobian stability analysis and the population-level association model, we compared dynamical signatures across the two perturbation modalities to assess whether chronic stress was associated with systematic changes in recovery dynamics and cross-pathway statistical associations. Because recordings from the two pathways were not obtained simultaneously, the estimated cross-pathway terms should be interpreted as cohort-level statistical associations within the reconstructed model rather than evidence of direct population-level dynamical association or causal interactions between pathways.

### 2.3 Neural Network Architecture: a Minimal Toy Model

The ANN framework was developed after the descriptive analyses. The purpose was not to infer mechanisms directly from the experimental recordings, but to examine whether a minimal set of asymmetric constraints could reproduce the qualitative signatures independently identified by the distributional and stability-related analyses.

Inspired by the experimentally reported asymmetry between the BLA-DMS and CeA-DMS pathways (Giovanniello et al. [5]), we constructed a simplified ANN framework to explore whether distinct optimization constraints could generate qualitatively different dynamical behaviors.

We generated time-series data where a high-frequency periodic baseline with Gaussian noise served as the homeostatic equilibrium. Environmental stressors were operationalized as stochastic impulse perturbations (spikes). The Control group was trained exclusively on the noise-buffered baseline, whereas the Stress group was exposed to a regime characterized by frequent stochastic perturbations. To simulate experimental probes (e.g., reward delivery or foot-shocks), we utilized a test signal comprising the standard homeostatic background interrupted by a single, discrete spike (as shown in Figure 13).

The experimental observation that BLA-DMS activity dominates in the Control group while CeA-DMS activity prevails in the Stress group suggests a context-dependent asymmetry in observed activity patterns under environmental pressure. We hypothesized that the experimentally observed asymmetry could be represented by subnetworks operating under different optimization pressures, one favoring precision-oriented tracking and the other favoring persistence-oriented buffering.

To computationally explore these dynamical patterns (see Section 3.1), we formulated the Unified Control System (UCS) within a simplified artificial neural network (ANN) framework. It is important to note that the UCS model is not intended to simulate the biological processes of BLA and CeA directly, but rather serves as a proof-of-principle model designed to explore the hypothesized trade-offs between precision-driven tracking and robust buffering. In this framework, the ”BLA-DMS” and ”CeA-DMS” pathways represent abstract sub-networks that capture the functional roles of these brain regions within the model, rather than actual biological circuits.

Specifically, asymmetric loss functions were introduced as hypothesis-driven modeling assumptions motivated by the experimental observations. These loss functions simulate distinct functional constraints on the sub-networks representing the BLA-DMS and CeA-DMS pathways, with the relative weighting and transition dynamics between these sub-networks learned during training. The design implements a division of labor between the sub-networks, with a Meta-Controller dynamically gating their contributions based on the environmental context.

The UCS model should be viewed as a minimal computational framework to explore functional asymmetry between the pathways, rather than a biologically realistic representation of neural circuits. It is designed to test whether asymmetric objectives can reproduce the observed dynamical behaviors, with the understanding that it is a simplified, abstract representation rather than a literal simulation of biological processes.

The UCS is formulated as a dual-pathway adaptive controller. In this framework, we define the fundamental state of each pathway by the relation:

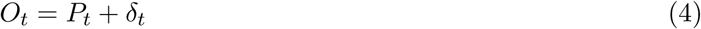

where *O_t_* is the observable output sent to the DMS, *P_t_* is the predicted calibration (baseline), and *δ_t_* is the latent deviation (accumulated internal strain). The system’s global objective is to minimize the total loss *L_T otal_* = *L_BLA_* + *L_CeA_* relative to the homeostatic reference (*T*), which corresponds to the aforementioned periodic baseline.

The internal dynamics of the BLA-DMS sub-network (*x_BLA_*) and the CeA-DMS sub-network (*x_CeA_*) are governed by an adaptive LSTM cell state, where the gating signals are modulated by the meta-learned parameter set Φ (see Figure 1). The two sub-networks are therefore trained under distinct objective functions, creating different optimization pressures during learning (for detailed values please refer to Supplementary Materials C). The asymmetric loss functions should therefore be viewed as hypothesis-driven design choices rather than emergent properties learned from the data. Consequently, successful reproduction of the empirical patterns should be interpreted only as evidence that the proposed asymmetries are sufficient to generate such behaviors within the model, not as evidence that biological circuits implement the same objectives.

**Figure 1:**
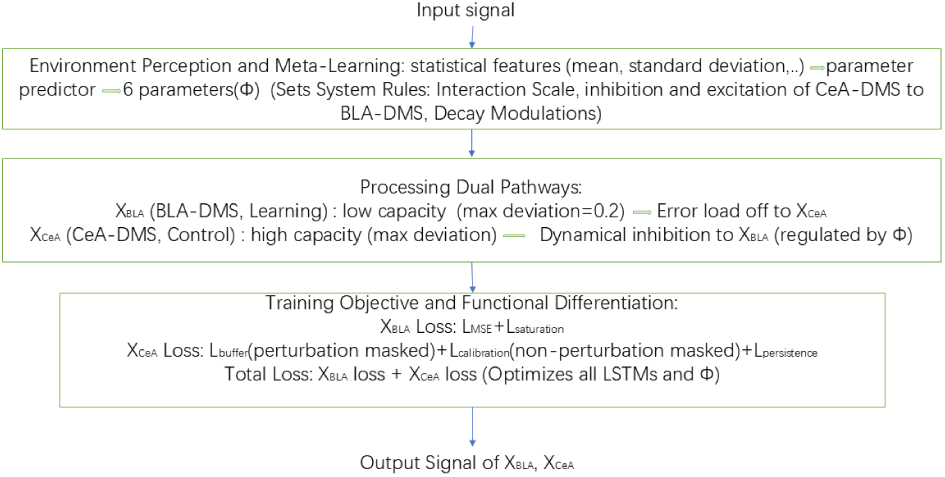
Schematic of the adaptive controller Unified Control System (UCS) architecture. The model illustrates a possible transition in the relative contribution of the two abstract sub-networks: (1) Macro-Perception Layer: A parameter predictor (Φ) that senses environmental volatility to dynamically adjust system dual-pathway architecture rules. (2) Dual-Pathway Processing Layer: Parallel Adaptive LSTMs representing the Precision-oriented (BLA-DMS) sub-network (*x_BLA_*, capacity-limited) and Robustness-oriented (CeA-DMS) sub-network (*x_CeA_*, persistence-favoring) circuits. (3) Optimization Layer: Functional differentiation driven by Asymmetric Loss functions, where *L_BLA_*prioritizes tracking precision and *L_CeA_* optimizes for robustness and informational persistence. The core mechanism is adaptive learning, enabling adaptive responses and pathway specialization through dynamic parameter modulation.

We defined two sub-networks: the Precision-Oriented sub-network as a Precision Tracker (Subnetwork A) and the Robustness-Oriented subnetwork as a Robustness Buffer (Subnetwork B). For interpretive convenience, Subnetwork A and Subnetwork B are later compared with experimentally observed BLA-DMS and CeA-DMS activity patterns.

The Precision-Oriented sub-network as a Precision Tracker (Subnetwork A): The abstract BLA-like sub-network is constrained by a fidelity-centric objective (*L_BLA_*), aiming to align *O_BLA_*with *T* under a hard physical capacity barrier (*θ_BLA_*):

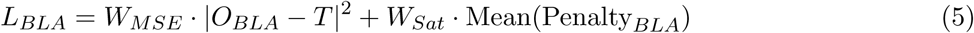

where *W_MSE_* and *W_Sat_* are weights for tracking precision and saturation penalty. The term Penalty*_BLA_* = ReLU(*|δ_BLA_| −* 0.2) simulates a ”brittle” system: while highly accurate in stable regimes, the BLA-DMS sub-network becomes functionally paralyzed when its latent strain *δ_BLA_* exceeds its metabolic ceiling (capacity constraint).

The Robustness-Oriented sub-network as a Robustness Buffer (Subnetwork B): CeA-like sub-network (*x_CeA_*) is assigned an objective emphasizing robustness and latent-state persistence under volatility. Its objective function (*L_CeA_*) is engineered to reward the buffering of perturbations and the maintenance of latent strain *δ_CeA_*, even when external stressors subside:

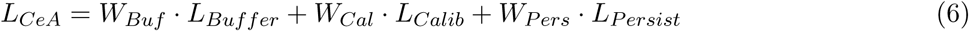

The interaction of these terms defines the ”viscous” dynamics of the habitual state: Dynamic Buffering ^(*L*^*Buffer* ^):^

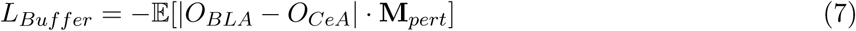

This term encourages the CeA-DMS sub-network to diverge from the BLA-DMS sub-network during perturbations (**M***_pert_*). In this context, CeA-DMS sub-network contributes to moderating the response of the Precision-Oriented sub-network during perturbations. Homeostatic Anchoring (*L_Calib_*)is defined as MSE(*O_CeA_, T*) during non-perturbation periods (**M***_non_*_−*pert*_). This term ensures the system remains tethered to the baseline in stress-free environments, preventing unrestricted drift. The differentiation depending on **M** that denotes the binary masks identifying perturbation segments ensures that while the BLA-DMS sub-network tracks homeostatic precision, the CeA-DMS sub-network retains a persistent latent representation of prior perturbations. Informational Persistence (*L_P ersist_*):

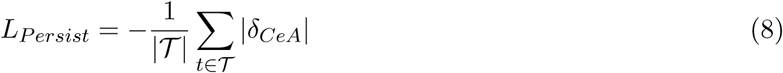

where *T* is the discrete temporal domain. By assigning a negative weight (*W_P ers_* = *−*2.0), the model rewards the maintenance of a non-zero *δ_CeA_*. This design choice encourages the persistence of non-zero latent states following perturbations, generating hysteresis-like behavior within the model that qualitatively resembles the distributional persistence observed in the experimental data. This allows the UCS framework to provide a computational illustration of how persistence-oriented objectives can generate recovery dynamics qualitatively similar to those observed experimentally.

This reward structure substantially alters the optimization landscape of the model. Unlike ”elastic” systems that gravitate toward zero deviation, the CeA-DMS sub-network develops a persistent latent representation of prior perturbations. This produces persistent post-perturbation activity within the model, qualitatively resembling the prolonged recovery patterns observed in the empirical data. Consistent with the observed dominance of model-defined CeA-like sub-network in stressed subjects, *L_CeA_* is specifically engineered to reward informational persistence, allowing the pathway to maintain an active latent state even as external perturbations subside.

The coordination between abstract subnetworks is governed by a hierarchical control structure: a Parameter Predictor sets macroscopic interaction scales based on long-term environmental statistics, while a Meta-Controller provides microscopic adaptation to instantaneous uncertainty.

The dual-pathway architecture of these pathways is governed by a hierarchical control structure: a Parameter Predictor sets macroscopic interaction scales based on environmental statistics, while a Meta-Controller provides microscopic adaptation to instantaneous uncertainty. Schematic construction of the neural network is shown in Figure 1.

in Figure 1, there is target asymmetry (Functional Division). This is a design where the BLA-DMS sub-network and CeA-DMS sub-network are optimized for different functional objectives, thereby formalizing the functional division between the precision-focused BLA-DMS sub-network and the stability-focused CeA-DMS sub-network.

As shown in Figure 1, this leads to a shift in the relative contribution of the two abstract sub-networks within the model: as the BLA-DMS sub-network reaches its saturation limit under high-entropy stress, the learned controller increasingly allocates computational load to the high-capacity CeA-DMS sub-network. This transition is quantified by the Max Deviation, defined as the peak internal strain max *|δ_t_|* during a perturbation. The systematic reallocation of deviation from the BLA-DMS sub-network to the CeA-DMS sub-network provides a computational metric describing the redistribution of latent deviation between the two abstract sub-networks. We refer to the two units as ”BLA-DMS sub-network” and ”CeA-DMS sub-network” purely for interpretability, based on their correspondence to the empirical signals. This naming does not imply anatomical or mechanistic equivalence.

To explore the consequences of training under different environmental statistics, we trained independent UCS instances on distinct environmental conditions (Control vs. Stress). Initially, both subnetworks were trained using identical MSE-based objectives. Under this configuration, the model did not reproduce the qualitative dynamical differences observed in the experimental data. This observation motivated the introduction of asymmetric objectives as a modeling hypothesis. The resulting behavior should therefore be interpreted as a proof-of-concept demonstration that such asymmetries are sufficient to generate the observed qualitative patterns, rather than evidence that biological circuits implement the same optimization strategy.

## 3 Results

### 3.1 Data Analysis

#### 3.1.1 Non-Learned Modality: Footshock Statistical Perspective

The temporal evolution of KL divergence following acute footshock to quantify stimulus-evoked deviation from baseline activity distributions and subsequent return dynamics (Figure 2). In the Control group, both pathways exhibited a transient increase in KL divergence followed by rapid recovery toward baseline within approximately 4 seconds. In the Stress group, the BLA-DMS pathway displayed a prolonged elevation in KL divergence that sustained deviation from baseline distribution during the observation period.

**Figure 2:**
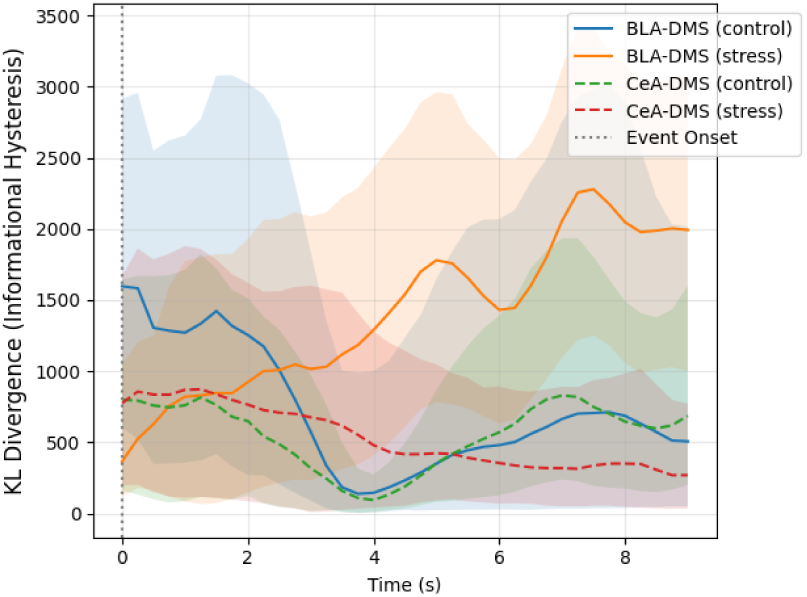
Time-resolved Kullback-Leibler (KL) divergence relative to the pre-stimulus baseline (*t <* 0) following a single footshock (*t* = 0, gray dotted line). Solid and dashed lines represent the mean KL divergence for BLA-DMS and CeA-DMS pathways, respectively; shaded areas denote the 95% confidence intervals (CIs) derived from subject-level bootstrapping (*n* = 1000 iterations). In control mice, KL divergence for both pathways returned toward baseline within approximately 4 s. In stressed mice, KL divergence of the BLA-DMS pathway remained elevated beyond 6 s, whereas the CeA-DMS pathway showed a smaller amplitude change.

In the stressed group, from *t* = 6 s onward, the mean KL divergence trajectories of the BLA-DMS and CeA-DMS pathways remained clearly separated. In contrast, in the Control group, the KL divergence trajectories of the two pathways converged following the perturbation.

To assess variability across subjects, we performed a subject-level bootstrap procedure (*n* = 1000 iterations) to estimate the cumulative KL divergence (Table 1). The Mean Cumulative KL divergence (*D_KL_*) for the stressed BLA-DMS pathway was higher than that of the control group (51663 vs. 26979). The 95% CI lower bound was also higher in the stressed group (25083 vs. 6137).

**Table 1:**
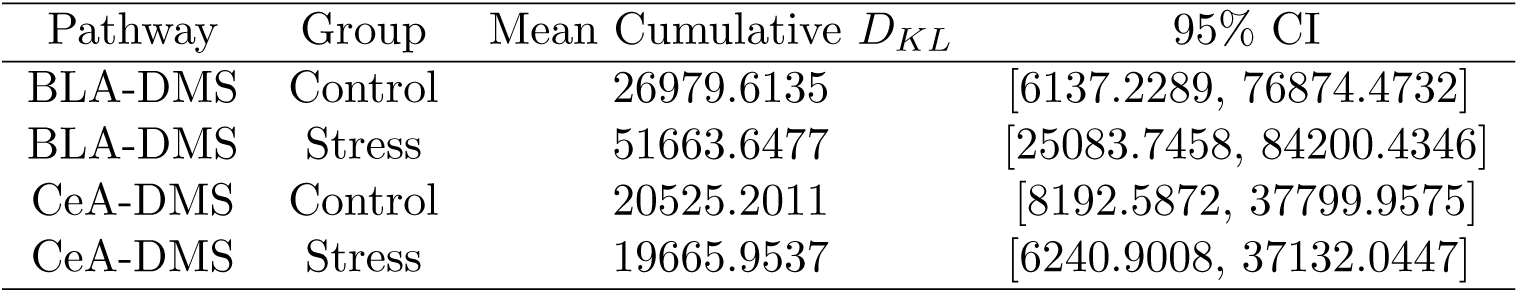
Statistical Robustness of Integrated KL Divergence Following Acute Footshock.

We compared time-resolved KL divergence with pre and post-perturbation distribution moments (Supplementary Materials Table D8). Mean and variance showed modest changes across conditions, whereas KL divergence showed larger magnitude differences between groups.

While KL divergence characterizes how strongly activity distributions deviate from baseline following perturbation, it does not directly describe the local recovery properties of the resulting trajectories. We therefore complemented the distributional analysis with a stability-related characterization based on phenomenological dynamical models. The purpose of this analysis is not to establish a formal mapping between KL divergence and Jacobian eigenvalues, but rather to examine whether both descriptive approaches reveal qualitatively consistent stress-dependent trends.

##### Dynamical Perspective

We next analyzed the stability regimes of the two pathways using Jacobian eigenvalues. Table 2 reports the mean coefficients and their corresponding 95% confidence intervals (CIs) derived from 1000 bootstrap iterations.

**Table 2:** Single Variable Model Statistics of Governing Dynamical Equations for Footshock Response.

| Pathway | Term | Control |  | Stress |  |
| --- | --- | --- | --- | --- | --- |
|  |  | Mean Coefficients | 95% CI | Mean Coefficients | 95% CI |
| BLA-DMS | Intercept | -1.9133 | [-11.0969, 6.5834] | -11.7957 | [-25.3569, -3.9355] * |
| | $x_{BLA}$ | -12.7695 | [-52.9504, 3.9237] | -8.0333 | [-16.1302, -1.5321] * |
| | $\dot{x}_{BLA}$ | -27.5925 | [-30.3323, -23.1249] * | -26.5726 | [-29.2188, -22.9836] * |
| | $x_{BLA}^2$ | -9.9306 | [-50.8948, 19.6649] | 18.2556 | [2.2446, 49.5706] * |
| | $x_{BLA}\dot{x}_{BLA}$ | -0.381 | [-6.3485, 5.3484] | 2.9982 | [-1.5469, 8.5104] |
| | $\dot{x}_{BLA}^2$ | 0.2087 | [-0.5831, 1.3130] | -0.069 | [-1.0170, 0.9458] |
| CeA-DMS | Intercept | -6.2783 | [-12.0011, -1.0509] * | -4.167 | [-10.3045, 6.9254] |
| | $x_{CeA}$ | -8.3031 | [-19.1907, -3.5590] * | -7.9044 | [-18.7739, 2.7631] |
| | $\dot{x}_{CeA}$ | -28.7741 | [-31.5479, -25.3304] * | -27.4448 | [-30.5188, -23.8047] * |
| | $x_{CeA}^2$ | 3.558 | [-20.5504, 14.2200] | 1.6879 | [-16.1770, 14.3044] |
| | $x_{CeA}\dot{x}_{CeA}$ | 3.2412 | [0.0413, 6.9261] * | 1.2205 | [-2.9535, 4.8927] |
| | $\dot{x}_{CeA}^2$ | 0.5948 | [-0.5281, 1.6150] | 1.1538 | [-0.5994, 2.8350] |

\* Indicates statistical significance (95% CI does not overlap with zero).

In the Stress group, the quadratic coefficient (*x*^2^) for the BLA-DMS pathway was consistently positive and statistically significant across all behavioral modalities, including footshock (Table 2), operant pressing, and reward delivery (see Supplementary Tables D5, D13, and D14).

To examine population-level recovery-rate characteristics, we derived the probability density of the principal eigenvalues (*Re*(*E*_1_)) via a subject-level bootstrap-regression pipeline (Figure 3). The principal eigenvalues were obtained from local linearization of the reconstructed phenomenological models around their estimated steady states. We interpret *Re*(*E*_1_) as a local stability indicator, while distributional shifts reflect changes in recovery speed rather than qualitative stability changes.

**Figure 3:**
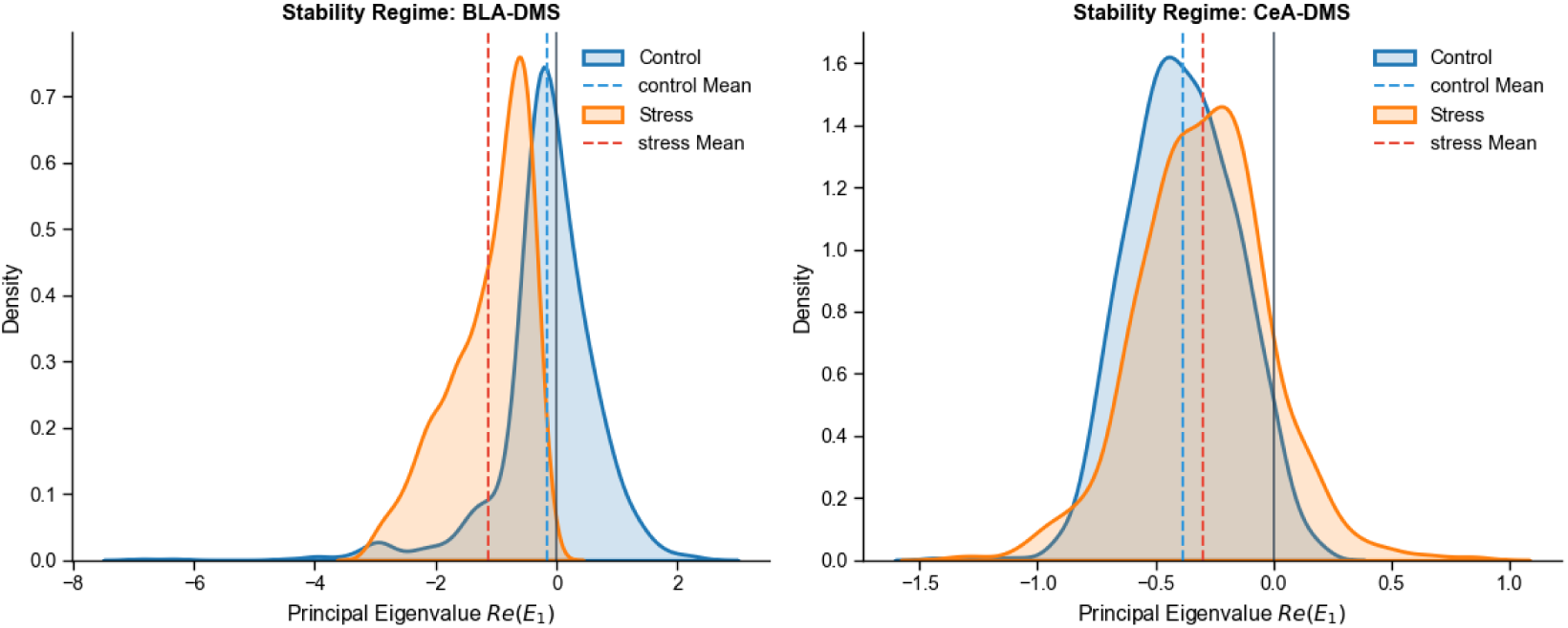
Population-level recovery-rate characteristics analysis of neural trajectories. The probability density functions (PDFs) of the principal Jacobian eigenvalues (*Re*(*E*_1_)) were estimated via subject-level bootstrapping (*n* = 1000 iterations) following acute footshock. (Left) In the BLA-DMS pathway, the distribution of *Re*(*E*_1_) for the stress group (orange) is shifted to more negative values compared to the control group (blue). (Right) In the CeA-DMS pathway, the distribution of *Re*(*E*_1_) for the stress group is shifted to more positive values relative to controls. Dashed lines represent the mean of the bootstrapped distributions.

In the BLA-DMS pathway, the Control group exhibited a distribution centered near zero. In the Stress group, the distribution of *Re*(*E*_1_) shifted leftward, with the 95% confidence interval strictly negative (Table 3).

**Table 3:**
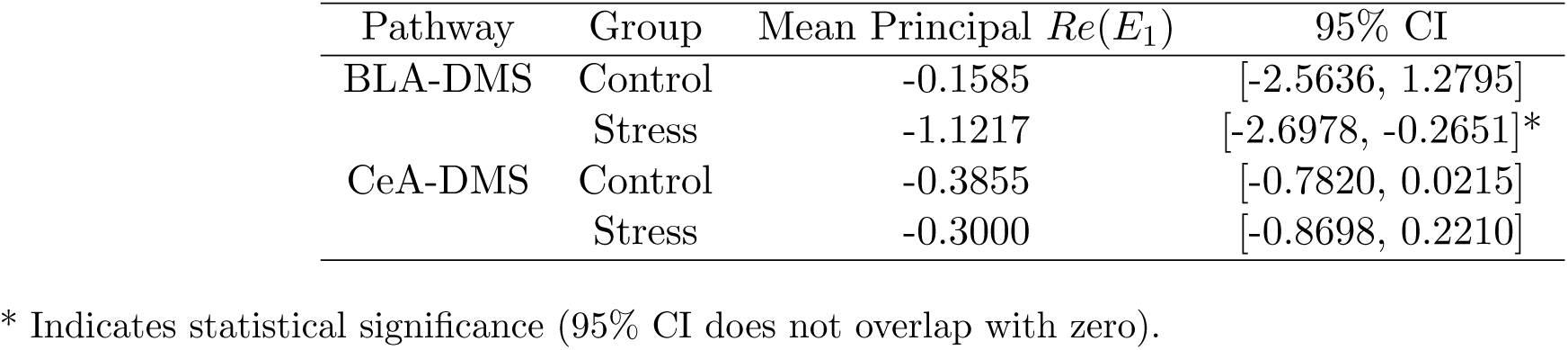
Statistics of Principal Jacobian Eigenvalues for Footshock.

| Pathway | Group | Mean Principal $Re(E_1)$ | 95% CI |
| --- | --- | --- | --- |
| BLA-DMS | Control | -0.1585 | [-2.5636, 1.2795] |
|  | Stress | -1.1217 | [-2.6978, -0.2651] * |
| CeA-DMS | Control | -0.3855 | [-0.7820, 0.0215] |
|  | Stress | -0.3000 | [-0.8698, 0.2210] |

\* Indicates statistical significance (95% CI does not overlap with zero).

The deterministic parameters of the ”average” model (Table 4) were consistent with the bootstrapped mean distributions. The norm of eigenvalues, quantified by the *L*^2^-norm of the eigenvalues (*||E||*_2_), was higher than 26 for most pathways, except for the Control BLA-DMS pathway.

**Table 4:** Deterministic Stability Parameters and Recovery Kinetics Post-footshock.

| Group | Pathway | Steady State |  | Eigenvalues |  | Norm of Eigenvalues |
| --- | --- | --- | --- | --- | --- | --- |
| | | $x_s$ | $\dot{x}_s$ | $E_1$ | $E_2$ | |
| Control | BLA-DMS | -0.397 | 0 | 1.60E-04 | -0.27275 | 0.273 |
|  | CeA-DMS | -0.3924 | 0 | -0.4044 | -29.469 | 29.472 |
| Stress | BLA-DMS | -0.6465 | 0 | -0.837 | -27.2812 | 27.294 |
|  | CeA-DMS | -0.1597 | 0 | -0.2615 | -26.8716 | 26.873 |

In the Stress group, the BLA-DMS pathway exhibited relatively high *||E||*_2_ values compared to the Control group. The CeA-DMS pathway in the Stress group showed eigenvalues of similar magnitude to those observed in controls.

The population-level regression models quantified condition-dependent differences in cross-pathway associations (Table 5 and 6). Across both regression models, the coefficients associated with the velocity terms (either *x_̇BLA_* or *̇x_CeA_*) were consistently statistically significant across groups, with confidence intervals excluding zero in all cases.

**Table 5:** Coefficients of Population-level Regression Model (BLA-DMS as target pathway)

| Variables | Control |  | Stress |  |
| --- | --- | --- | --- | --- |
|  | Mean Coefficients | 95% CI | Mean Coefficients | 95% CI |
| $x_{BLA}$ | -128.9280 | [-239.7098, -33.8852]* | -8.7873 | [-71.4567, 35.6582] |
| $\dot{x}_{BLA}$ | -32.5210 | [-37.6582, -26.5185]* | -29.2282 | [-35.7247, -23.9503]* |
| $x_{BLA}^2$ | -22.6382 | [-164.0459, 105.3996] | 56.2701 | [-7.3992, 116.3720] |
| $x_{BLA}\dot{x}_{BLA}$ | -9.5272 | [-29.8239, 7.8918] | 1.0535 | [-9.3516, 9.0369] |
| $\dot{x}_{BLA}^2$ | -0.2753 | [-1.4229, 0.7945] | -0.2047 | [-1.4371, 0.9042] |
| $x_{CeA}$ | 80.8810 | [23.8786, 144.2312]* | 8.2858 | [-23.6643, 44.6718] |
| $\dot{x}_{CeA}$ | 1.8124 | [-1.6390, 5.1101] | 1.4626 | [-1.1804, 4.5300] |
| $x_{BLA}x_{CeA}$ | 46.7142 | [-53.7033, 187.4726] | -25.8531 | [-85.4224, 40.9728] |
| $x_{BLA}\dot{x}_{CeA}$ | 4.7782 | [-1.3883, 15.1915] | 3.4656 | [-2.2191, 11.0586] |
| $\dot{x}_{BLA}x_{CeA}$ | 8.7838 | [-5.3136, 28.8629] | 1.3723 | [-4.9063, 8.9948] |
| $\dot{x}_{BLA}\dot{x}_{CeA}$ | 0.4897 | [-0.7424, 1.9411] | 0.8116 | [-0.5674, 2.3390] |

\* Indicates statistical significance (95% CI does not overlap with zero).

In the Control group Table 5, BLA-DMS as target pathway for regression, the linear coefficient corresponding to *x_CeA_* was 80.88 with a 95% confidence interval excluding zero (95% CI: [23.88, 144.23]). In the Stress group, the same coefficient did not reach statistical reliability (Figure 4). Other cross-pathway coefficients, including *x_BLA_* and interaction terms, showed confidence intervals overlapping zero in most cases.

**Figure 4:**
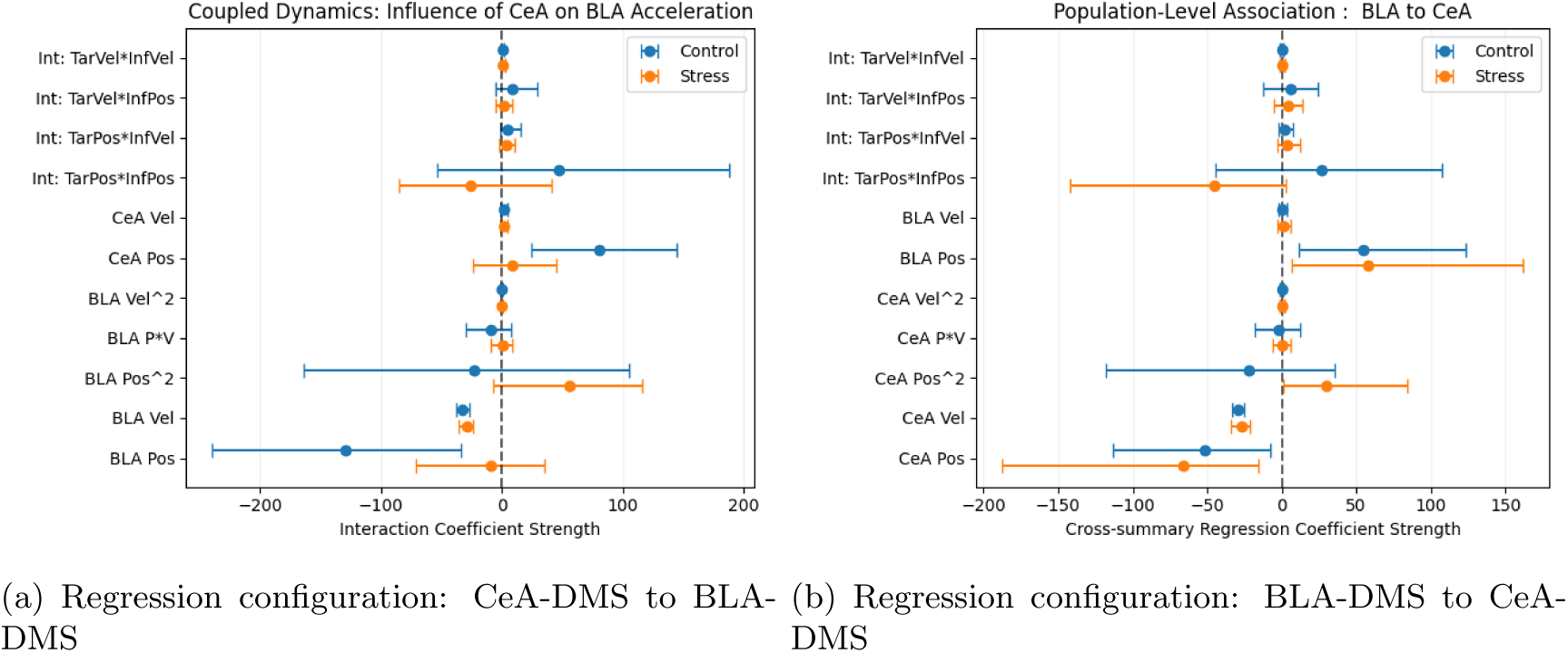
Forest plots show cross-summary regression coefficients from the population-level dynamical model. Variables are defined as Target (Tar) and Influence (Inf), where Pos corresponds to signal amplitude (*x*) and Vel corresponds to its first derivative (*̇x*). Each point represents the mean coefficient estimated from subject-level bootstrapping (*n* = 1000), with error bars indicating 95% confidence intervals (CIs). Coefficients whose confidence intervals do not include zero are marked as statistically reliable.

In the model that CeA-DMS as target for regression (Table 6), the quadratic coefficient *x*^2^ was 29.96 remained positive and statistically reliable in the Stress group (95% CI: [1.42, 84.21]). Other cross-pathway coefficients displayed heterogeneous changes in effect size and confidence interval overlap between groups. Other coefficients were generally not statistically significant, as indicated by confidence intervals overlapping zero.

**Table 6:** Coefficients of Population-level Regression Model (CeA-DMS as target pathway)

| Variables | Control |  | Stress |  |
| --- | --- | --- | --- | --- |
|  | Mean Coefficients | 95% CI | Mean Coefficients | 95% CI |
| $x_{CeA}$ | -51.4172 | [-113.5551, -8.0192]* | -65.9611 | [-187.4206, -15.7276]* |
| $\dot{x}_{CeA}$ | -29.3639 | [-32.8705, -25.6075]* | -27.1137 | [-33.9473, -20.8905]* |
| $x_{CeA}^2$ | -21.9424 | [-117.9379, 35.3150] | 29.9629 | [1.4243, 84.2127]* |
| $x_{CeA}\dot{x}_{CeA}$ | -2.2302 | [-17.9178, 12.2457] | 0.2663 | [-5.9641, 5.6891] |
| $\dot{x}_{CeA}^2$ | 0.1302 | [-1.0966, 1.3994] | 0.3835 | [-0.8955, 1.7972] |
| $x_{BLA}$ | 54.8296 | [11.2364, 123.4697]* | 58.1055 | [6.7276, 162.0060]* |
| $\dot{x}_{BLA}$ | 0.5796 | [-2.1818, 3.6602] | 0.8940 | [-2.5930, 5.7940] |
| $x_{CeA}x_{BLA}$ | 26.4998 | [-44.4399, 107.2743] | -44.9540 | [-141.7844, 2.6415] |
| $x_{CeA}\dot{x}_{BLA}$ | 1.6595 | [-2.3736, 7.8194] | 3.4341 | [-3.0802, 12.2634] |
| $\dot{x}_{CeA}x_{BLA}$ | 5.7149 | [-12.6906, 24.2480] | 4.6674 | [-5.0549, 14.1652] |
| $\dot{x}_{CeA}\dot{x}_{BLA}$ | 0.2680 | [-0.8885, 1.4286] | 0.4331 | [-1.0463, 1.9704] |

\* Indicates statistical significance (95% CI does not overlap with zero).

In stressed mice, several coefficients that reached statistical reliability in the Control group no longer exhibited confidence intervals excluding zero. Following Footshock (Non-Learned Modality), stressed BLA-DMS trajectories exhibited larger deviations from baseline distributions than controls. During Reward (Learned Modality), group differences in trajectory distributions were comparatively smaller (Supplementary Materials Table D6).

#### 3.1.2 Learned Modality: Lever Pressing-Reward Association Statistical Perspective

During the Contingency Acquisition phase (Giovanniello et al., 2025), KL divergence was measured as mice transitioned from deterministic (FR1) to stochastic (RR10) reinforcement (Figure 5). In control mice, the BLA-DMS pathway exhibited KL divergence values near baseline throughout RR10. In the stressed group, the BLA-DMS pathway showed a sustained increase in KL divergence during RR10, following an initial surge during FR1. The rise in KL divergence was larger and more prolonged in stressed mice compared to controls.

**Figure 5:**
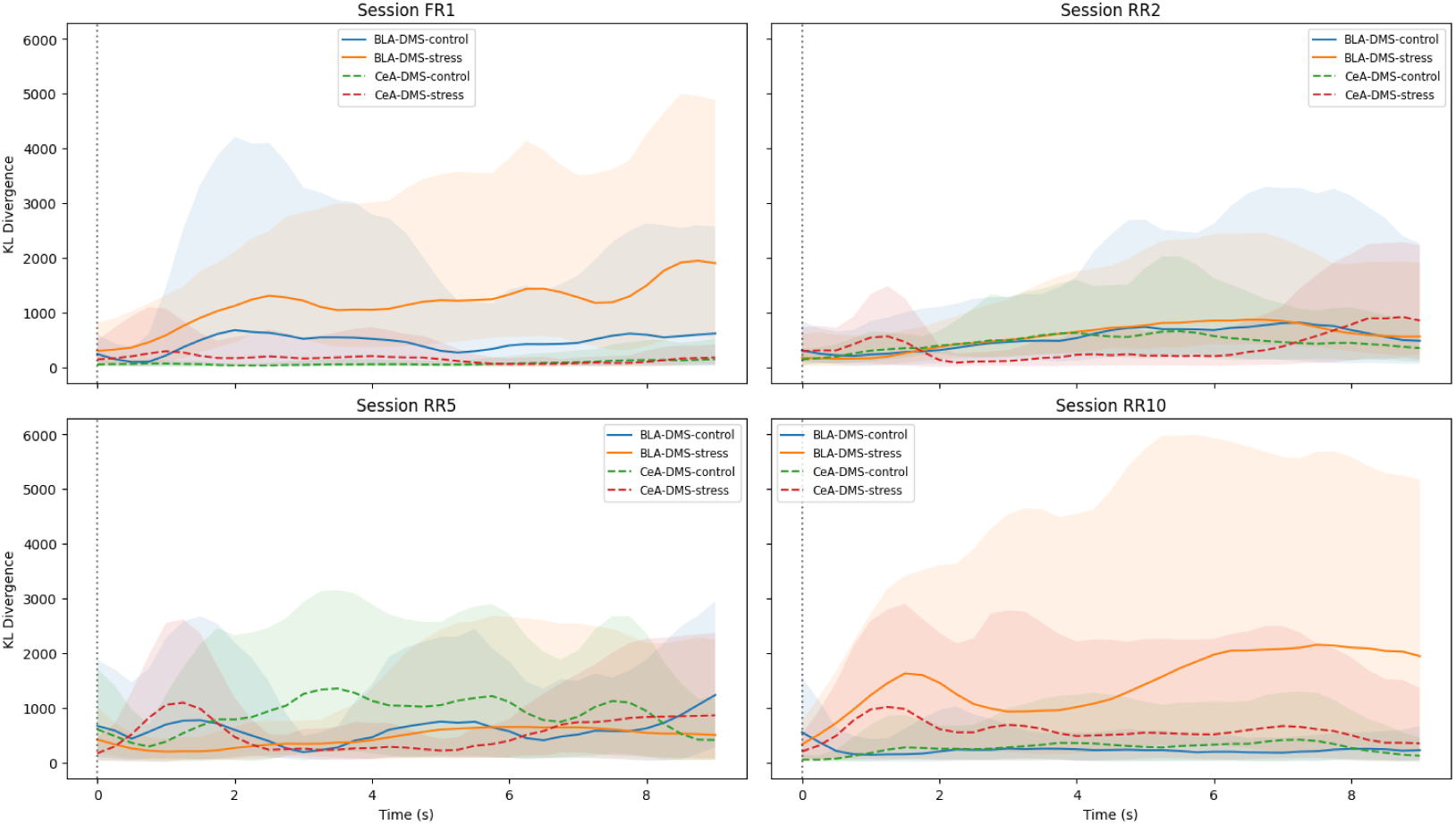
Evolution of KL divergence during operant lever pressing. Time-resolved KL divergence of BLA-DMS (solid lines) and CeA-DMS (dashed lines) aligned to lever press onset across four training stages (FR1, RR2, RR5, and RR10). (Top-Left) During early acquisition (FR1), the stressed group (orange/red) shows higher KL divergence in BLA-DMS compared to controls (blue). (Bottom-Right) During late-stage stochastic reinforcement (RR10), control mice show KL divergence values near baseline, whereas stressed mice exhibit a sustained increase in BLA-DMS KL divergence. Shaded areas represent 95% confidence intervals derived from subject-level bootstrapping (*n* = 1000). KL divergence values are shown on a consistent scale across panels.

The temporal evolution of the 95% confidence intervals (shaded areas) was quantified across training stages (FR1, RR2, RR5, RR10) aligned to lever press onset (Figure 5). In the FR1 stage, control mice exhibited relatively broad confidence intervals around 4 s post-press. Across RR2 and RR5, confidence intervals in the control group remained broad. At RR10, the control group showed a reduction in confidence interval width.

In contrast, the stressed BLA-DMS group exhibited higher KL divergence values across RR stages compared to controls, with no comparable reduction in variability at RR10.

Following operant lever presses, KL divergence was measured aligned to reward delivery (Reward-onset) in BLA-DMS and CeA-DMS pathways (Figure 6).

**Figure 6:**
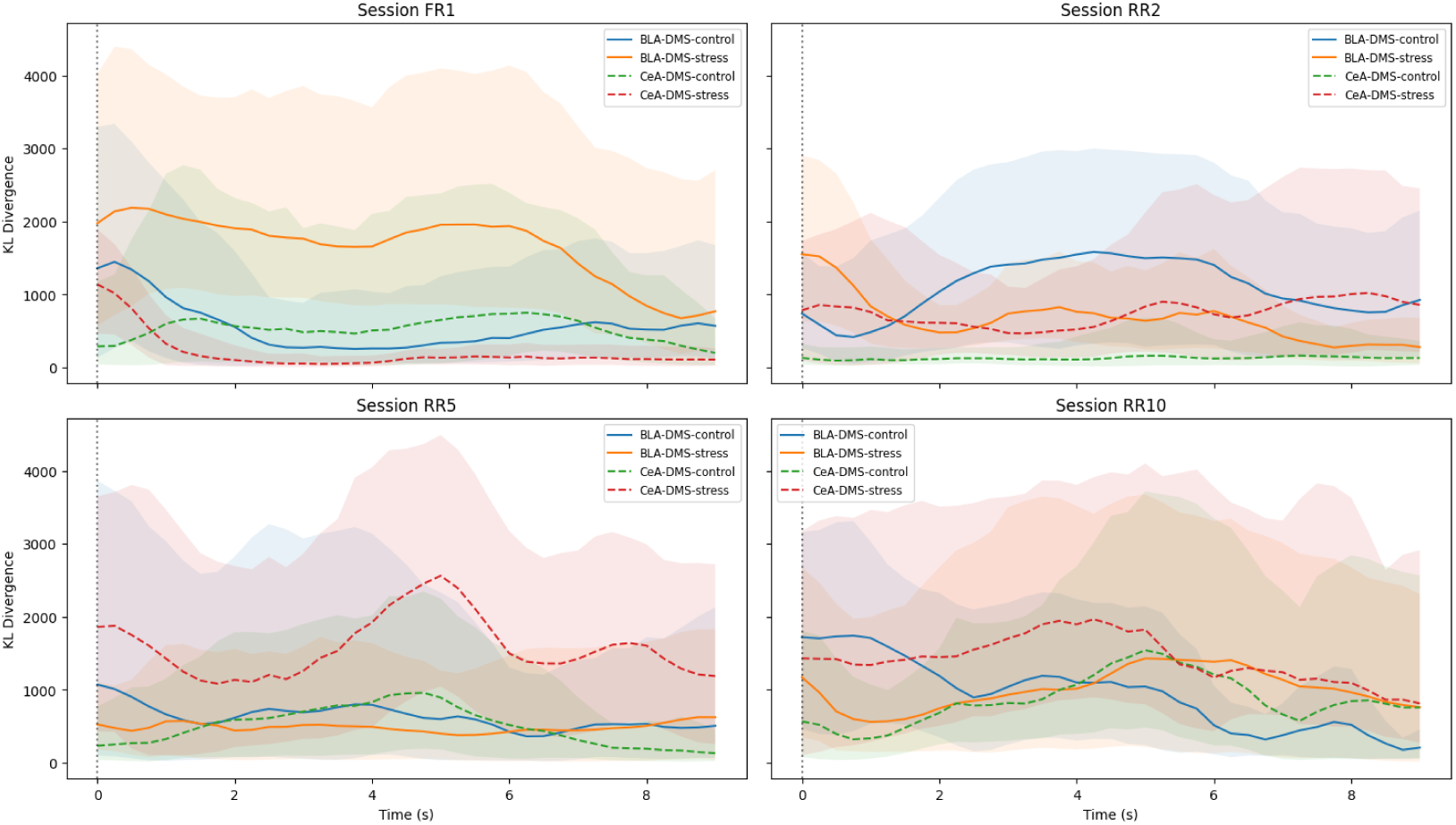
Evolution of KL divergence following lever-press reinforcement. Time-resolved KL divergence trajectories are aligned to the delivery of food reinforcement following lever presses across training sessions (FR1, RR5, RR10). In FR1, the CeA-DMS pathway in the stressed group (red dashed line) shows lower KL divergence compared to controls. In later sessions (RR5–RR10), the stressed CeA-DMS pathway exhibits higher and more sustained KL divergence compared to controls (green dashed line). Shaded areas represent 95% confidence intervals.

In the initial acquisition phase (FR1), stressed mice exhibited lower KL divergence trajectories in CeA-DMS (red dashed line) compared to controls. Across later stages with higher reinforcement uncertainty (RR5, RR10), KL divergence in stressed CeA-DMS increased and remained elevated relative to the control group. In BLA-DMS, stressed mice also showed sustained deviations from control KL divergence trajectories during these stages.

Terminology Note: In the figures, variables are labeled as Target (Tar) and Influence (Inf) for consistency in depicting regression inputs and outputs. The mapping between these graphical terms and the anatomical variables (*x_BLA_*, *x_CeA_*) used in the regression tables is provided in the Main Text.

##### Dynamical Perspective

Beyond informational metrics, we employed single variable dynamical models and population-level regression models to characterize condition-dependent changes in pathway activity patterns. Full coefficient tables for all training sessions and reinforcement regimes are provided in the Supplementary Materials (Tables D13–D14 and Tables D18-D19).

In the single variable dynamical models, the principal eigenvalue’s real part (*Re*(*E*_1_)) remained negative across all training sessions (FR1 to RR10) for both BLA-DMS and CeA-DMS pathways (Figures 7 and 8), with 95% confidence intervals reported in Supplementary Materials D.

**Figure 7:**
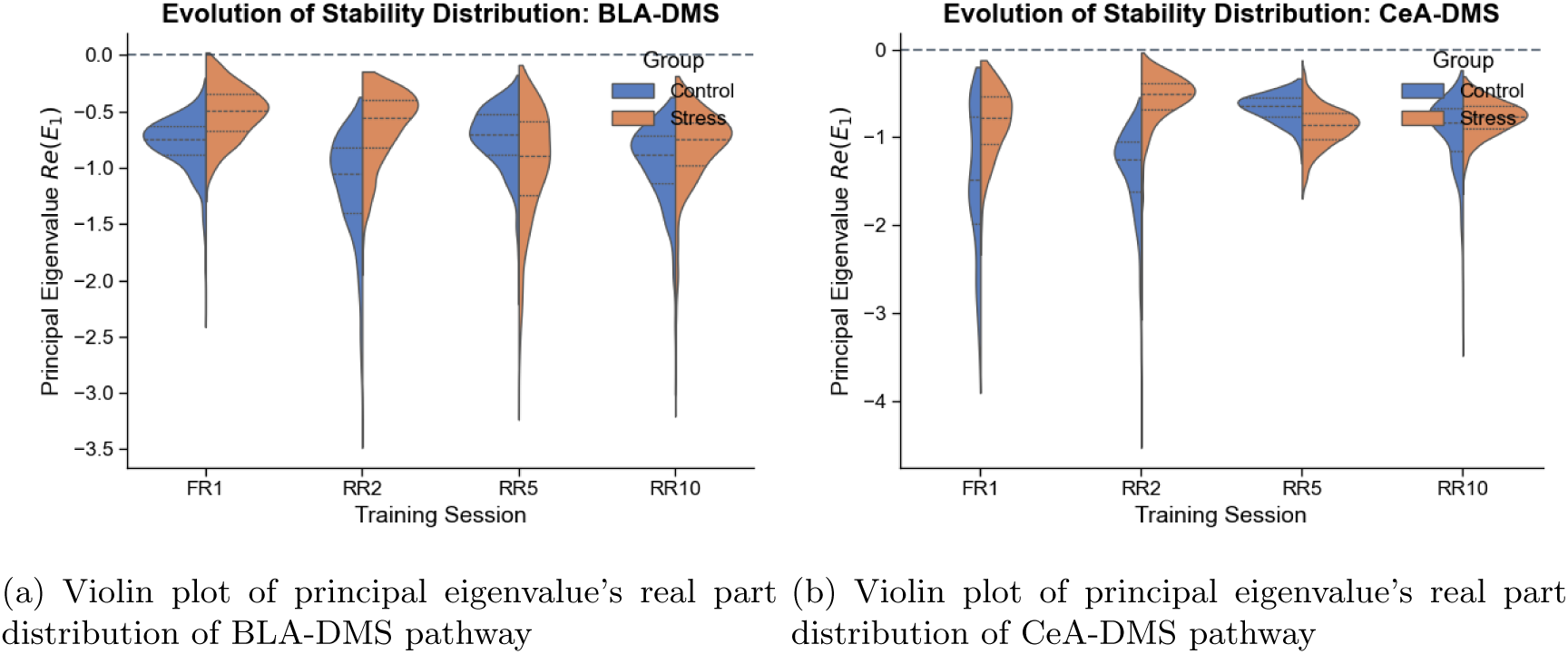
Violin plots and range trajectories represent the distribution of the principal eigenvalue’s real part (*Re*(*E*_1_)) derived from bootstrapped ridge regression (*n* = 1000) for bout-initiating pressing. Across all reinforcement regimes (FR1 to RR10), the *Re*(*E*_1_) distributions remain negative (*Re*(*E*_1_) *<* 0) for both BLA-DMS and CeA-DMS pathways.

**Figure 8:**
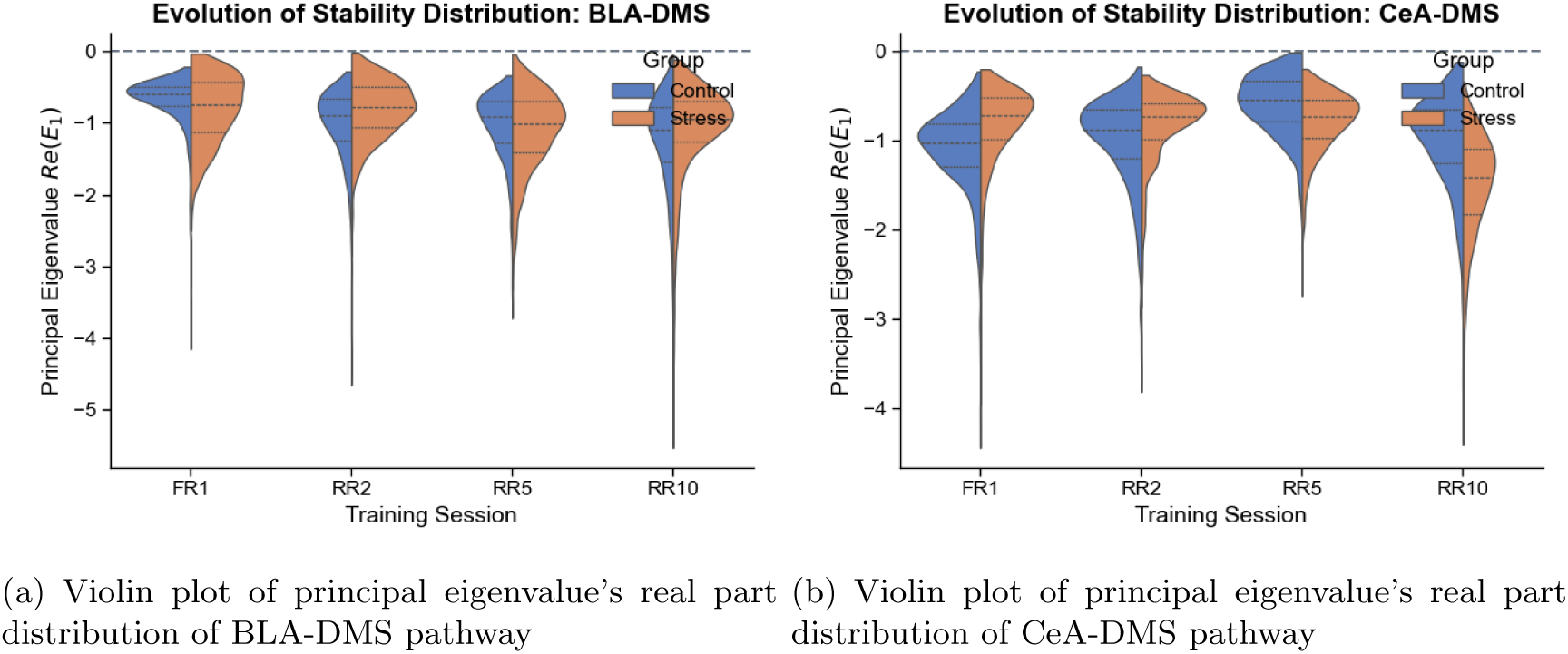
Violin plots and range trajectories represent the distribution of the principal eigenvalue’s real part (*Re*(*E*_1_)) derived from bootstrapped ridge regression (*n* = 1000) for reward. Across all reinforcement regimes (FR1 to RR10), the *Re*(*E*_1_) distributions remain negative (*Re*(*E*_1_) *<* 0) for both BLA-DMS and CeA-DMS pathways.

This stability contrasts with the transient eigenvalue excursions occasionally observed in control mice during unpredicted footshocks.

While the distributions of the principal eigenvalues remained within the negative domain across sessions, the population-level regression models exhibited condition-dependent differences in coefficient structure. Examination of the regression coefficients across all experimental phases (Tables 5–6 and Supplementary Tables D9-D10 and D18-D19) revealed several recurring patterns.

Across reinforcement schedules (FR1–RR10) and behavioral epochs, coefficients associated with the first-derivative terms (*̇x_BLA_* and *̇x_CeA_*) were consistently negative, with bootstrap 95% confidence intervals excluding zero in most models. The estimated coefficients generally ranged from approximately *−*24 to *−*32 (Supplementary Tables D18-D19). In contrast, the significance and magnitude of linear, quadratic, and cross-pathway terms varied across behavioral conditions and experimental groups.

In control mice, both the BLA-DMS and CeA-DMS pathways exhibit consistently negative coefficients for positional terms (*x_BLA_, x_CeA_*) across the majority of models examined (Figures 9(a)-10(c)). These coefficients remain significantly different from zero, indicating reliable within-pathway trends in the regression models.

**Figure 9:**
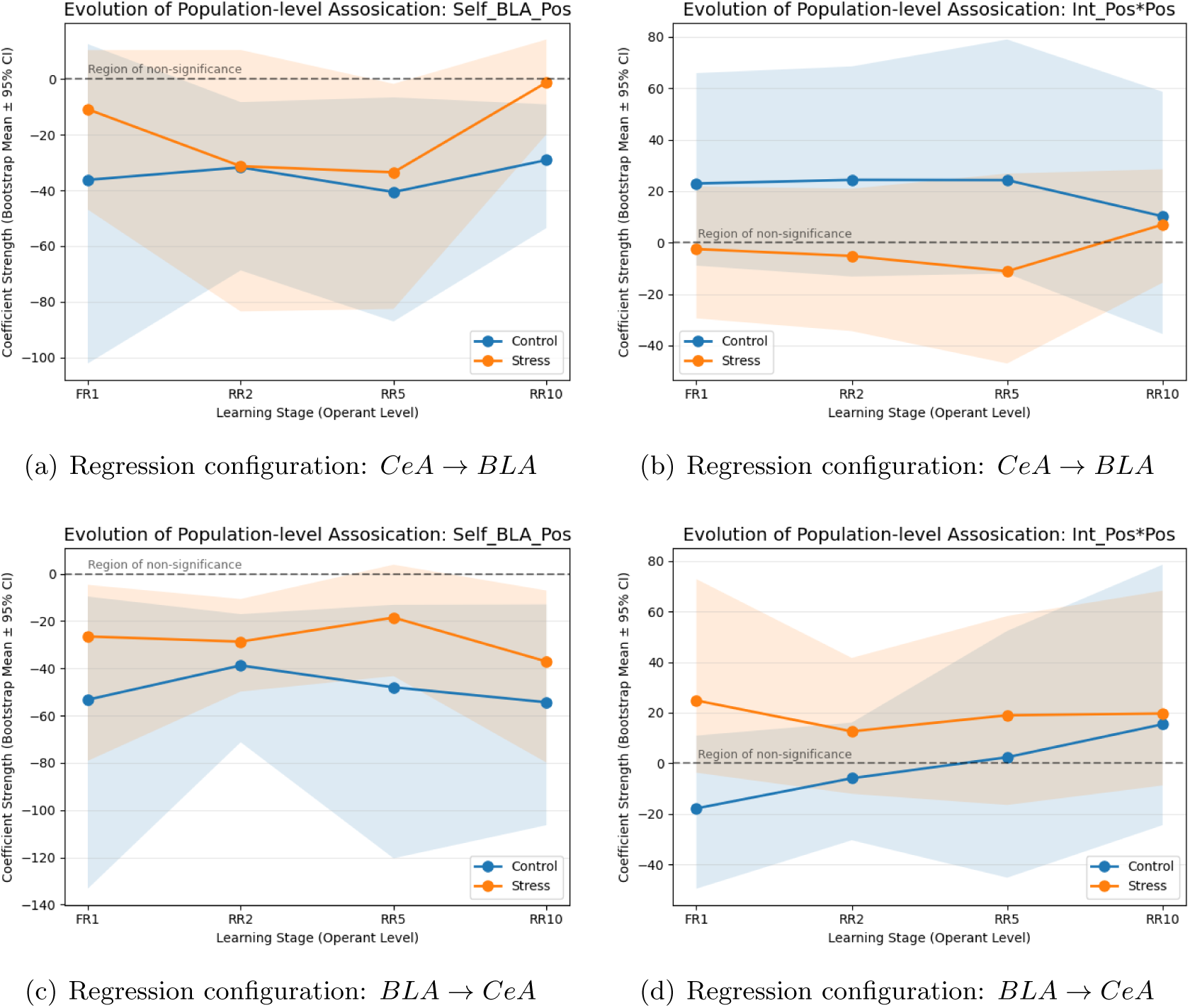
Evolution of key association coefficients during bout-initiating pressing. Parameter trajectories from *n* = 1000 bootstraps show regression coefficient changes from FR1 to RR10. (9(a), 9(c)) Self-feedback positional terms (*x_CeA_* and *x_BLA_*). Control mice (blue) coefficients remain consistently negative with 95% CIs excluding zero, while stressed mice (orange) show coefficients drifting toward non-significance (CIs overlapping zero). (9(b), 9(d)) Nonlinear modulation term (*x_CeA_x_BLA_*). Most coefficients remain non-significant across groups, with overlapping CIs. Shaded areas represent 95% CIs.

In the stressed group, the corresponding positional coefficients frequently reach non-significance (confidence intervals crossing zero, Figures 9-10). This pattern suggests a stress-related alteration in the distribution of explanatory contributions between pathways in the multivariate regression.

**Figure 10:**
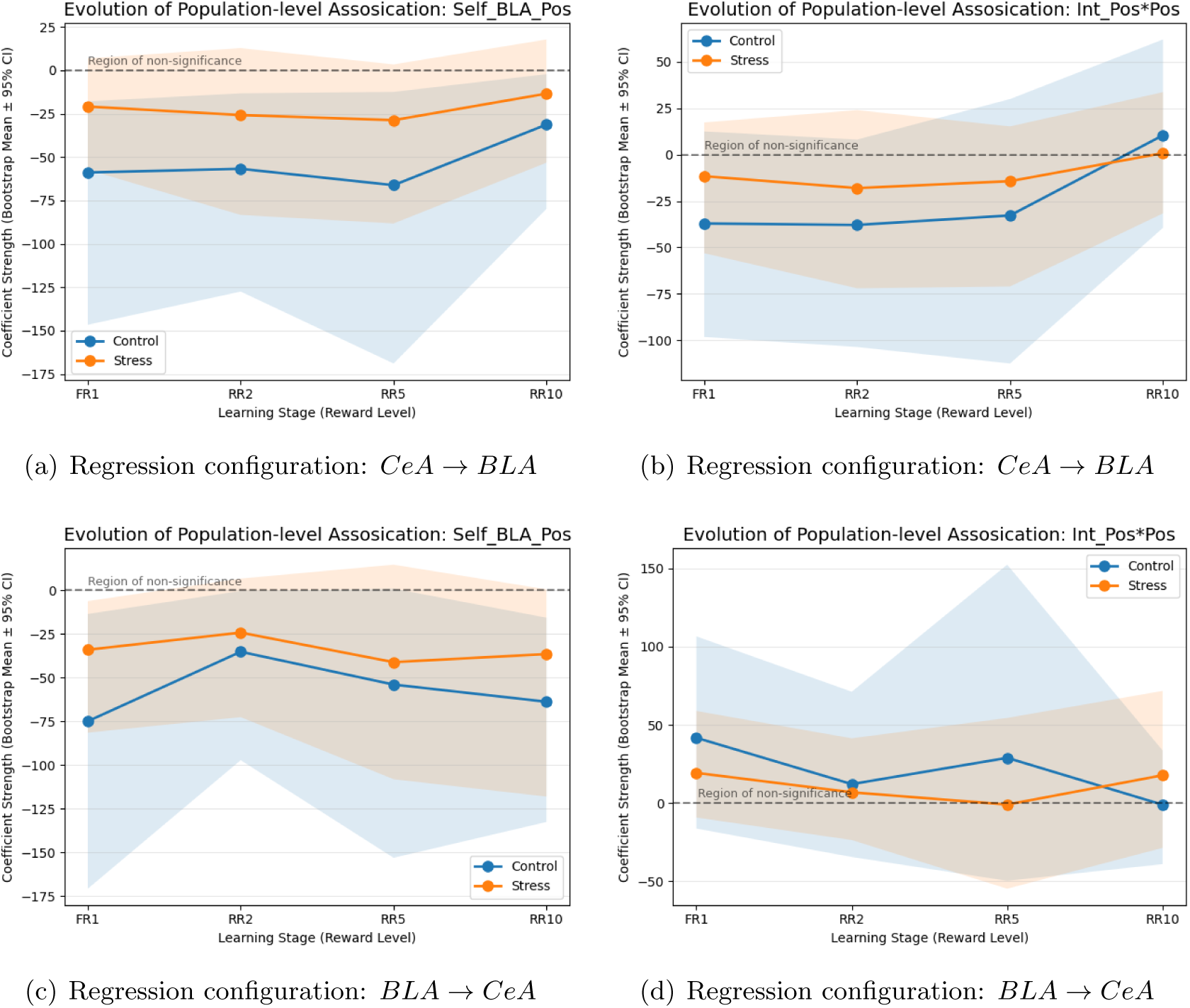
Dynamical association coefficients during reward processing. (10(a), 10(c)) Self-feedback positional terms (*x_CeA_* and *x_BLA_*). Control mice maintain negative coefficients with 95% CIs excluding zero; stressed mice show coefficients frequently reaching non-significance with wider CIs. (10(b), 10(d)) Cross-pathway nonlinear modulation term. Coefficients remain mostly non-significant across groups. Shaded areas represent 95% CIs.

Additionally, positive quadratic terms (*x*^2^) that were significant in univariate models for the stressed group no longer reach significance in the joint regression framework. This change indicates that some variance previously captured by self-feedback nonlinear terms may be redistributed among cross-summary coefficients. Individual cross-pathway terms do not always achieve statistical significance, but their inclusion modifies the attribution of variance in the coupled regression model.

Across both univariate and multivariate population-level regression models, coefficients for velocity terms (*̇x_BLA_, ̇x_CeA_*) remain consistently negative and statistically robust. This consistency suggests that these terms contribute stably to the model across conditions, independent of the significance patterns observed in positional or cross-pathway coefficients.

When considered together across footshock, lever-pressing, and reward-associated models, a common coefficient structure emerged. Coefficients associated with first-derivative terms consistently retained statistical reliability across both single-variable and population-level models, whereas the significance of positional, nonlinear, and cross-pathway terms varied across behavioral contexts and experimental conditions. This pattern suggests that velocity-dependent contributions represent a recurring feature of the reconstructed phenomenological dynamics, while the explanatory contribution attributed to other terms varies across contexts.

Overall, chronic stress appears to modify the pattern of coefficient significance and interrelationships between terms, rather than eliminating individual contributions entirely. The observed differences reflect alterations in the statistical structure of the multivariate population-level regression models rather than a direct mechanistic interpretation.

#### 3.1.3 Changes in Reconstructed Phase Space Local Topology Across Conditions

To further compare the statistical and model-derived representations of pathway activity, we examined the reconstructed effective quasi-potential representation *U* (*x*) derived from the fitted univariate models (Figure 11).

**Figure 11:**
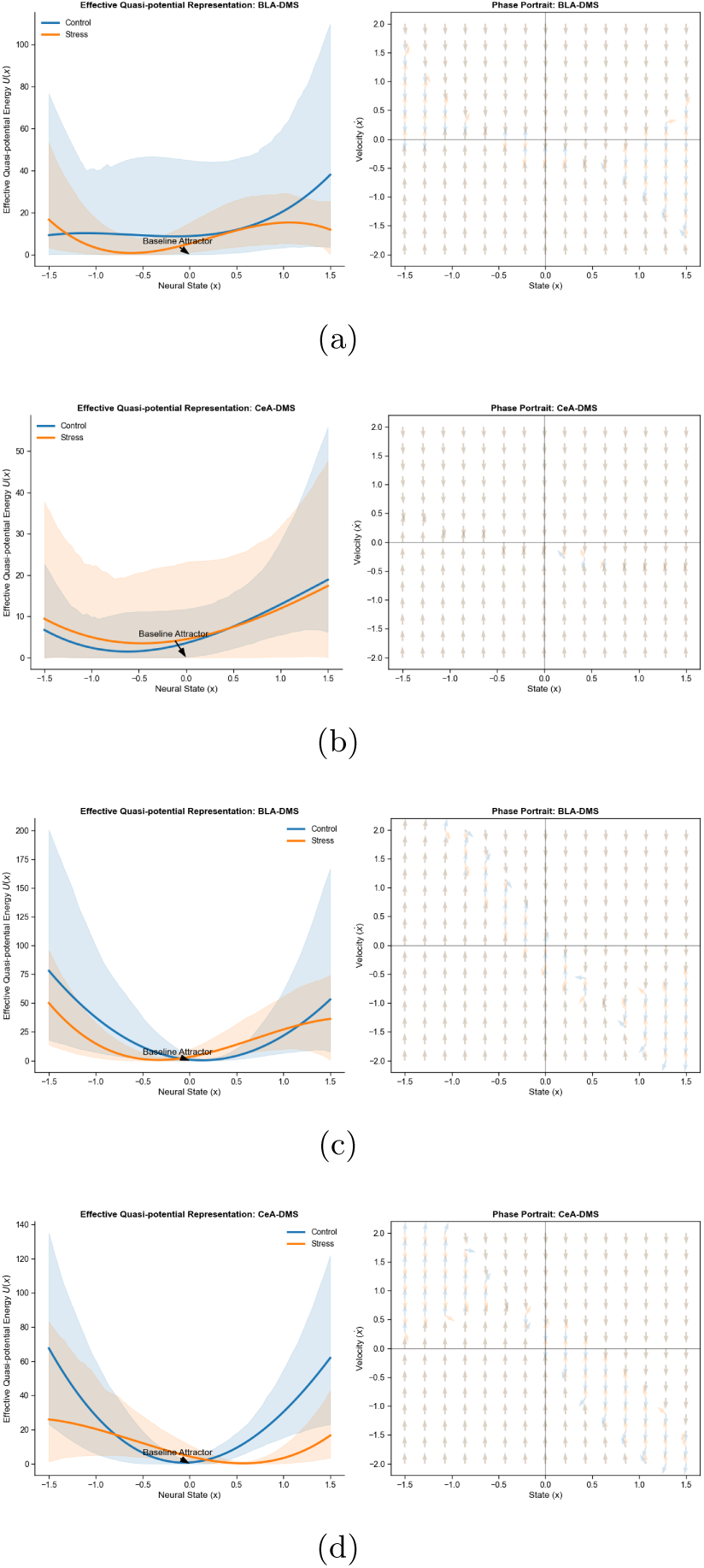
Reconstructed Effective Quasi-potential Representation across Modalities. Quasi-potential and phase portraits of neural circuit dynamics. Blue: Control; Orange: Stress. Shaded areas: 95% bootstrap CIs. For each subgraph, left: Effective quasi-potential *U* (*x*), right: phase portraits (*̇x, x*). 11(a) and 11(b) Non-learned Aversive Modality (Footshock): Stressed BLA–DMS (11(a)) exhibits a significant steady state displacement toward a biased state, while CeA–DMS (11(b)) maintains stability. 11(c)-11(d) Learned Reward Modality (Unpredicted Reward): Both BLA–DMS 11(c) and CeA–DMS 11(d) show qualitatively similar phase space local topology across groups, with noticeable broadening of the reconstructed quasi-potential profile in the stressed CeA–DMS.

Following footshock, the reconstructed effective quasi-potential representation of the BLA-DMS pathway in the Control group exhibited a minimum located near *x ≈* 0 (Figure 11(a)). In the Stress group, the location of the minimum was shifted toward negative values. Similar differences were observed in the corresponding phase portraits, where trajectories converged toward different equilibrium locations across groups.

Interestingly, this stress-induced displacement in the reconstructed quasi-potential representation in the BLA–DMS pathway was modality-specific. While the circuit underwent displacement following footshock, it maintained comparable similar local stability-related characteristics during the processing of unpredicted reward (learned modality) (Figure 11(c); detailed metrics in Supplementary Materials D).

To compare the statistical and model-derived quantities, we calculated correlations between steady-state KL divergence and the principal Jacobian eigenvalue *Re*(*E*_1_) (Table 7; Figure 12). In the footshock condition, the Stress BLA-DMS pathway exhibited a negative correlation between steady-state KL divergence and *Re*(*E*_1_)(*r* = *−*0.215*, p <* 10^−11^). In the unpredicted reward condition, the CeA-DMS pathway exhibited positive correlations in both groups, with larger coefficients observed in the Stress group (*r* = 0.327*, p <* 10^−25^; Table 7). These correlations should not be interpreted as evidence of a direct theoretical relationship between KL divergence and local stability, but rather as an empirical consistency check across two descriptive levels of analysis.

**Figure 12:**
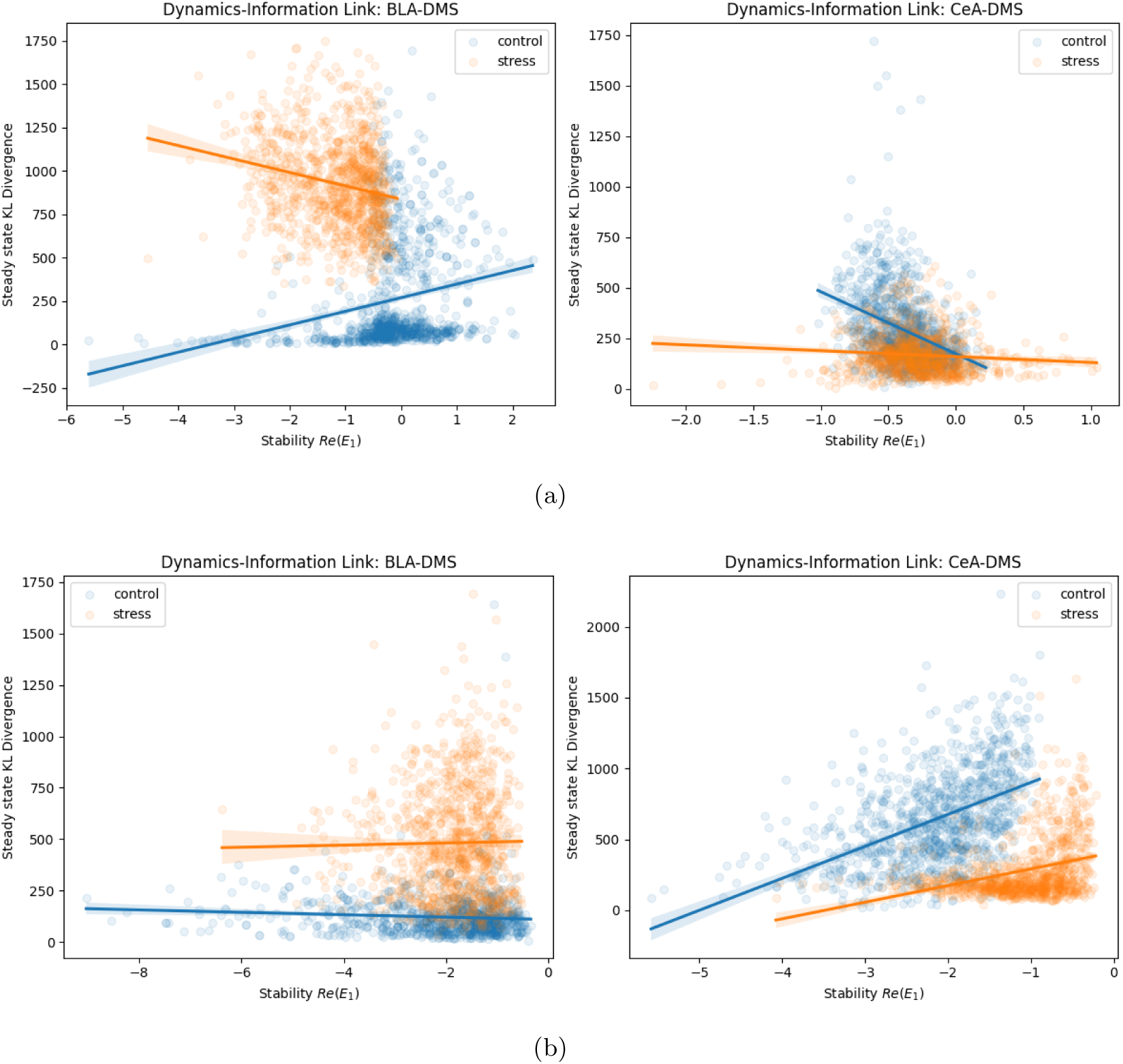
Empirical Alignment Between Informational and Dynamical Metrics. Relationship Between Steady-state KL Divergence and Principal Eigenvalues. Scatter plots showing the relationship between steady-state KL divergence and the principal eigenvalue *Re*(*E*_1_). Correlation coefficients and significance levels are reported in Table 7.

**Table 7:**
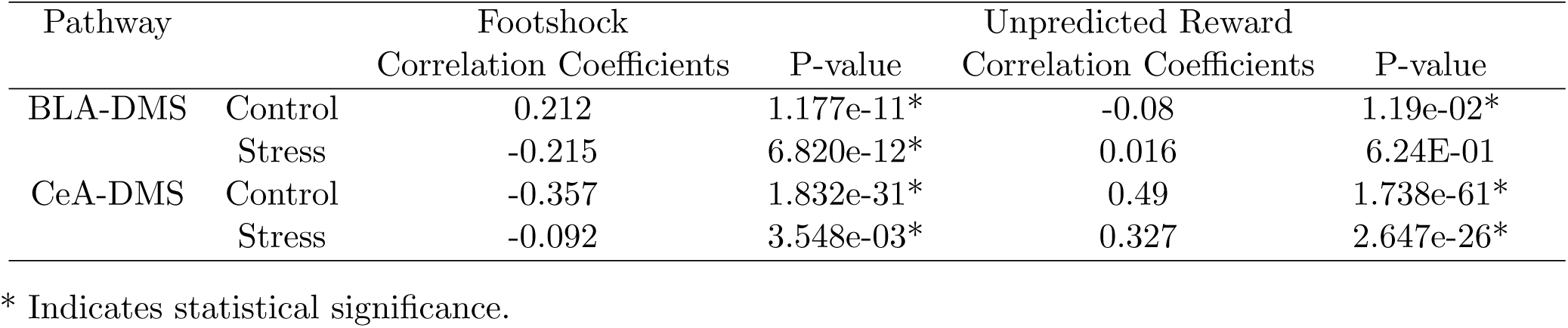
Correlation Coefficients Between Steady-state KL Divergence and Real Part of Principle Eigen-value.

**Figure 13:**
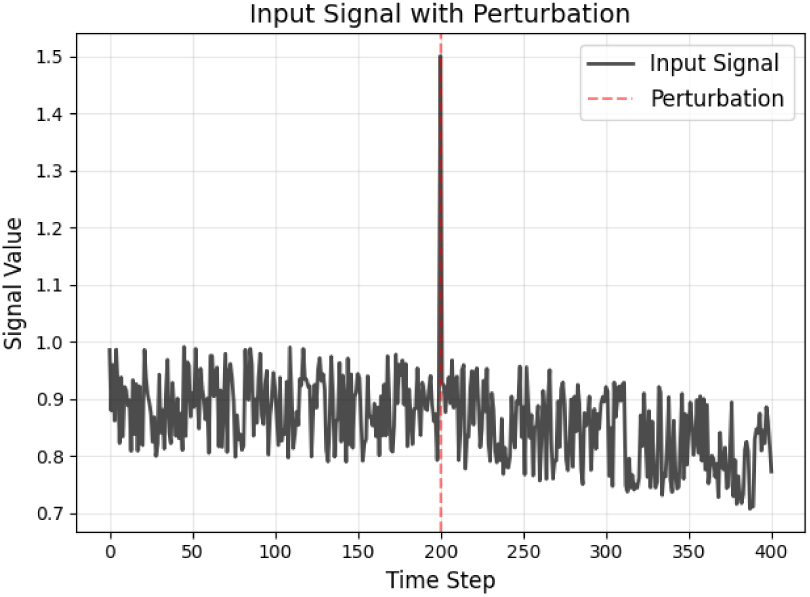
Simulation of test signals. The signal combines predictable periodicity with stochastic noise. The central spike mimics an acute high-impact perturbation (such as footshock), serving as a probe to test circuit stability.

These results indicate that both KL-based and dynamical measures exhibit condition-dependent structure, although they quantify different aspects of the data. The direction and magnitude of the KL-eigenvalue correlations varied across pathways and stimulus modalities (Table 7).

Remarks: It should be noted that the subject-level bootstrap procedure introduces stochastic variability across iterations, reflecting individual differences in neural responses. These analyses should not be inter-preted as implying a shared underlying generative model across statistical and dynamical representations. Despite these fluctuations, the overall statistical patterns are consistent across bootstrap samples. Specifically, the differences in regression coefficients between the footshock (Non-Learned) and reward/operant (Learned) modalities are maintained across resampling iterations. These patterns were consistently ob-served across bootstrap samples.

### 3.2 ANNs Model Simulation

Because both distributional and stability-related analyses revealed consistent stress-dependent alterations, we next asked whether a minimal set of asymmetric local constraints would be sufficient to reproduce selected qualitative signatures observed experimentally. The ANN was therefore used as a hypothesis-generating sufficiency test rather than a predictive or validation framework.

The Unified Control System (UCS) was constructed as a proof-of-concept computational framework motivated by two empirical observations from the preceding analyses. The model was not intended as a validation of the biological mechanisms, but rather as an exploratory framework for examining whether similar qualitative behaviors could emerge under predefined computational constraints:

Constraint Inspired by Distributional Persistence: The KL divergence analysis (Section 2.2) revealed persistent deviations from the pre-perturbation distribution. To represent such persistence in a simplified computational setting, we implemented adaptive LSTM units whose internal states allow information to be retained across time.

Constraint Inspired by Stability-Related Differences: The dynamical analyses (Section 3.1) identified consistent differences in regression coefficients and eigenvalue distributions between pathways and conditions. To incorporate these empirical asymmetries into the simulation framework, we assigned different objective functions to the two subnetworks. These objectives were selected to reproduce qualitative differences in response persistence and recovery observed in the experimental data and should not be interpreted as direct physical counterparts of the regression coefficients.

By incorporating these empirically motivated constraints, the UCS provides a computational framework for exploring how distinct optimization objectives may generate different response profiles under stochastic perturbations. This allows us to examine whether macroscopic differences in response profiles can be reproduced under distinct optimization objectives in a simplified computational setting.

To investigate the computational principles underlying stress-induced circuit remodeling, we trained independent instances of the Unified Control System (UCS). The models were tasked with adapting to distinct environmental statistics: a noise-buffered periodic baseline (Control) versus a regime characterized by frequent stochastic impulse perturbations (Stress).

The synthetic input sequences were designed to capture the core statistical features of the experimental environment (Figure 13): a periodic cosine component (*T* = 2*π/*80) representing predictable cycles, Gaussian noise (*σ* = 0.3) for internal fluctuations, and discrete anomalous pulses (amplitude 1.5) as computational analogs of acute footshocks.

The stable convergence of the total loss function (*L_T otal_*) across training epochs (Figure 14) confirms that both models successfully internalized their respective environmental statistics. Within the present simulation framework, reproducing the targeted response profiles was achieved using asymmetric optimization (*L_BLA_* vs. *L_CeA_*) objectives assigned to the two subnetworks.

**Figure 14:**
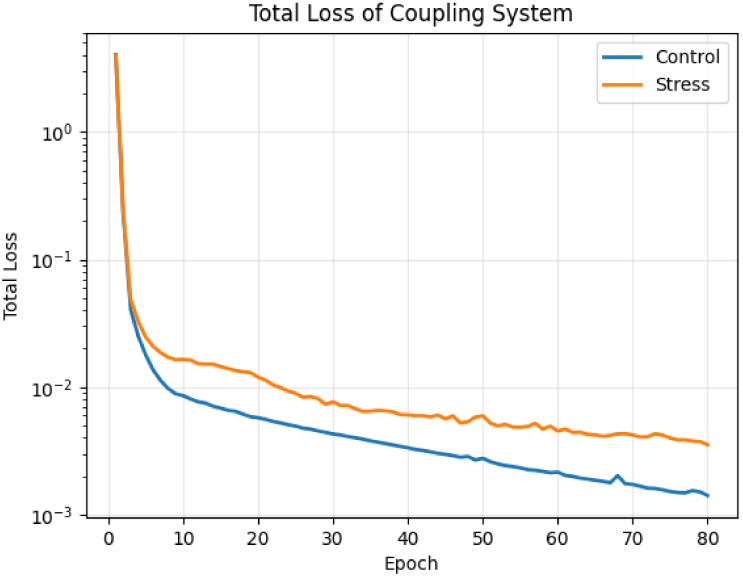
Training Convergence and Learning Stability. The trajectories of the total loss function for Control (blue) and Stress (orange) models demonstrate stable convergence across 80 epochs. The higher converged loss in the Stress group differences in optimization difficulty under distinct input statistics.

**Figure 15:**
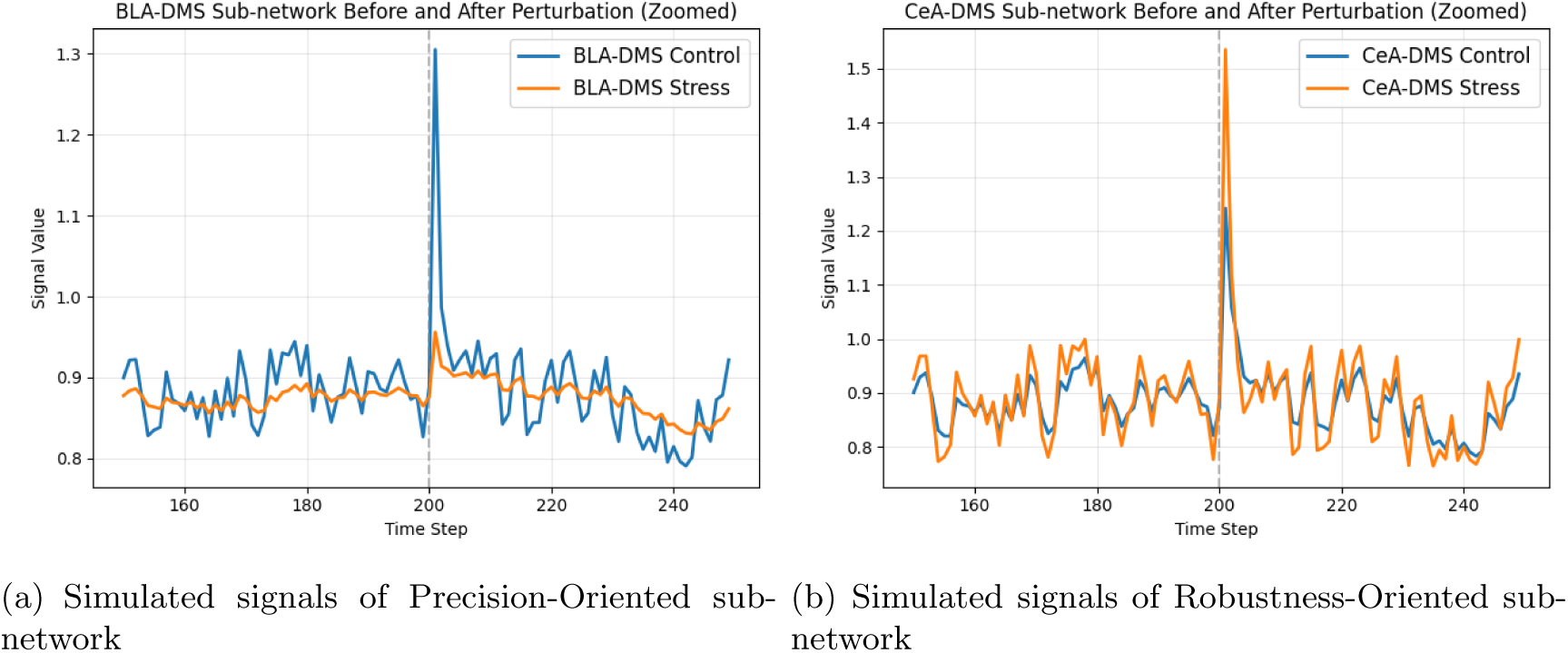
Simulated Subnetwork Responses to Acute Perturbation under the Computational Sufficiency-Testing Framework. Representative responses of the Precision-Oriented and Robustness-Oriented subnetworks under the two optimization regimes. Orange and blue lines correspond to the Stress and Control configurations, respectively.

To address the sensitivity of the model to initialization, we validated the architecture using 10 independent random seeds. The results demonstrate that the functional bifurcation is a consistent emergent pattern across random initializations of the asymmetric loss functions rather than a stochastic artifact.

We utilized the Max Deviation (Max*δ* = max*_t_ |δ_t_|*) as a proxy metric capturing peak deviation in internal state trajectories by each circuit axis (Figure 16).

**Figure 16:**
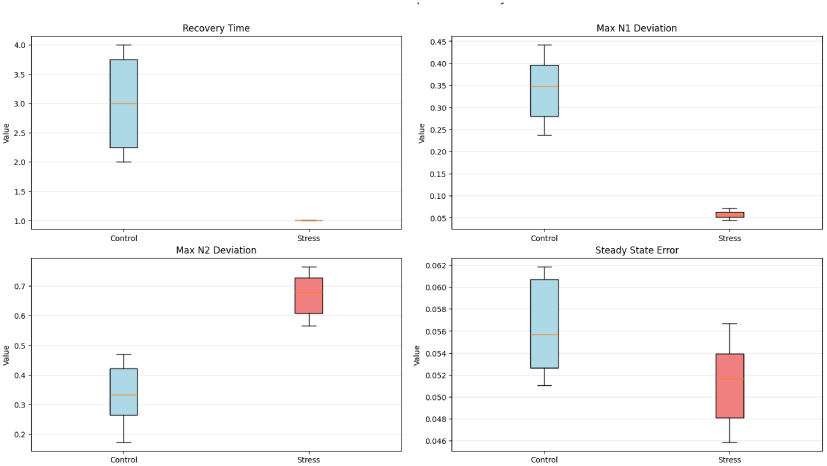
Robustness Metrics. Analysis across 10 independent random seeds suggests the stability of the UCS architecture. Top Left: Recovery time is significantly shorter in the Stress group, indicating more rapidly stabilizing regime. Top Right/Bottom Left: Max Deviation (Max*δ*) metrics illustrate a redistribution of peak deviation across the two sub-networks under different training regimes from the capacity-limited BLA-DMS sub-network in controls to the robust CeA-DMS sub-network in stressed models. Bottom Right: Lower steady-state error in the stress group is consistent with stronger stabilizing dynamics of the Robustness-Oriented sub-network in the model, resulting in reduced variability of output trajectories under this simulation setting.

Simulation outcomes differed between the two optimization regimes.

In the Control configuration, the Precision-Oriented subnetwork showed a larger Max Deviation (0.3390.07) and longer Recovery Time (3.000.77) steps.

In the Stress configuration, the Precision-Oriented subnetwork exhibited a lower Max Deviation (0.057), whereas the Robustness-Oriented subnetwork showed a larger Max Deviation (0.670). Recovery Time was reduced to 1.00 step.

Within the simulation framework, the two optimization regimes produced different distributions of Max Deviation across subnetworks. No direct inference regarding biological pathway function can be made from A key emergent feature of our ANN simulations is the decoupling of structural variability from functional stability. Across 10 random seeds, recovery time showed little variability (CV *≈* 40%, Supplementary Table C3), whereas the learned coupling weights varied substantially (Supplementary Table C3). This result indicates that similar behavioral outputs can be achieved under diverse parameter realizations within this simplified model.

## 4 Discussion and Limitations

### 4.1 Summary of Main Findings

The results suggest that chronic stress is associated with systematic alterations in the organization of neural activity patterns rather than a simple amplification or suppression of pathway activity. This alteration became most evident when neural circuits were challenged by external perturbations and was consistently reflected across both distributional and dynamical analyses.

Across behavioral modalities, chronic stress was associated with alterations in the temporal organization of activity distributions. These alterations were expressed not only through larger deviations from baseline distributions but also through changes in the variability of the bootstrap-derived KL trajectories. Under increasing reinforcement uncertainty, stressed animals exhibited broader distributional dispersion, suggesting greater heterogeneity in how pathway activity patterns evolved across subjects. (Figures 5,6)

The strongest differences between groups emerged following acute footshock. In the BLA-DMS pathway, control animals exhibited principal eigenvalue distributions centered closer to zero, with a substantial fraction of bootstrap estimates extending into the positive domain. Consistent with this observation, the reconstructed quasi-potential profiles were comparatively flatter, indicating weaker local confinement around the estimated equilibrium state. In contrast, the stressed BLA-DMS pathway showed principal eigenvalue distributions that remained predominantly negative and quasi-potential profiles with more pronounced curvature around the equilibrium. The CeA-DMS pathway displayed substantially smaller differences between groups, with largely overlapping eigenvalue distributions and qualitatively similar quasi-potential structures, although the control distributions were somewhat narrower. Although these analyses do not establish mechanistic circuit remodeling, they suggest that chronic stress alters how neural activity trajectories respond to external perturbations. (Tables 3 and Figures 3, 11(a))

During learned reward-related behaviors, both groups successfully acquired the lever press–reward association, consistent with the behavioral observations reported by Giovanniello et al. [5]. Principal eigen-values remained predominantly negative across reinforcement schedules, indicating locally stable recovery dynamics in both groups. At the same time, the pattern of statistically reliable coefficients differed across conditions, model formulations, and pathways. Importantly, these dynamical differences emerged during the acquisition phase, when overt behavioral performance remained largely comparable between groups. In the experimental study, behavioral divergence became evident only after changes in reinforcement contingencies. The present analyses therefore provide a complementary perspective: although the available data do not permit causal inference, the reconstructed dynamical models suggest that stress-related differences in neural activity organization may already be detectable before they become apparent at the behavioral level. The behavioral observations reported by Giovanniello et al. [5] indicate that stress-exposed animals eventually express altered action-selection strategies under changing reinforcement conditions. Our analyses suggest that signatures consistent with this later behavioral divergence may already be present in the statistical organization of pathway activity during the acquisition phase, even when overt task performance remains comparable. (Table D6, Figures D5, 7,8)

Across all behavioral modalities, derivative-dependent terms remained consistently reliable in both single-variable and population-level models. In contrast, the significance of positional and nonlinear terms varied across pathways and stress conditions. This recurring pattern suggests the presence of a conserved dynamical component that persists across contexts, whereas other aspects of the reconstructed dynamics remain sensitive to stress exposure and behavioral state. (Tables 2, 5, 6 and Supplementary D5, D9, D10, D13-D19)

We do not claim that KL divergence and Jacobian eigenvalues quantify the same dynamical quantity. Nevertheless, despite originating from distinct analytical frameworks, both measures revealed consistent stress-dependent alterations and exhibited non-random statistical associations across several experimental conditions. We therefore interpret these observations as an empirical alignment between distributional and stability-related descriptions rather than evidence of a formal theoretical correspondence.

Taken together, the results suggest that chronic stress is associated with a reorganization of neural activity patterns across multiple descriptive levels, from population-level activity distributions to local recovery dynamics. Although the present analyses do not establish causal mechanisms, they indicate that stress-related alterations become most apparent when neural circuits are challenged by external perturbations and may already be detectable before overt behavioral divergence emerges. These observations provided the rationale for the subsequent ANN sufficiency analyses, which were designed to examine whether a minimal set of asymmetric computational constraints could reproduce selected qualitative signatures identified in the experimental data.

### 4.2 Limitations

Several limitations should be acknowledged. The analyses rely on perievent recordings and non-simultaneous pathway measurements, preventing direct inference regarding real-time circuit interactions. Furthermore, no formal mathematical correspondence between KL divergence, dynamical parameters, and ANN representations is assumed. Future studies using simultaneous recordings and richer dynamical measurements may help determine whether these descriptive levels can be integrated within a more comprehensive theoretical framework. The population-level regression models should not be interpreted as estimates of real-time interactions between BLA-DMS and CeA-DMS pathways.

### 4.3 Future Directions

Although the present analyses were not designed to assess neural criticality, several observations motivate future investigation using frameworks specifically developed for criticality and near-critical dynamics. In particular, the tendency of control animals to exhibit eigenvalue distributions closer to the stability boundary, together with the shift toward more constrained recovery dynamics under chronic stress, raises the possibility that stress may alter the balance between dynamical flexibility and stability. Future studies combining simultaneous recordings, higher-dimensional state-space reconstruction, and criticality-oriented analyses may help determine whether these observations reflect broader changes in the operating regime of the underlying neural circuits [17, 18, 19].

## Declaration of generative AI in scientific writing

Authors declare no using of generative AI.

## Acknowledgements

We thank the authors of Giovanniello et al. (2025) [5] for sharing their datasets publicly.

This article is dedicated to the memory of my father, Weize Lin (born 1941-07-29 – passed away 2025-05-19). For a long time, I sought to understand his complex and challenging behaviors through the lens of psychological theory. However, at the beginning of 2025, I had a sudden epiphany: that these behavioral patterns, while appearing psychological on the surface, are essentially the outputs of a neural network trained and shaped by experience. This realization led me to develop a computational model to simulate these dynamics. Shortly thereafter, in February 2025, I encountered the relevant research published in Nature that provided the precise experimental datasets needed to validate this modeling approach. Although he passed away before the completion of this work, this study stands as a tribute to his profound influence on my thinking, representing the culmination of a journey from personal observation to scientific discovery during a time of great loss.

## Author contributions

Contribution of the authors: Feng Lin analyzed the data and constructed the framework, developed the analytical methodology, implemented the codes and wrote the manuscript.

## Data availability

The data used in this study were originally reported by Giovanniello et al. [5] and are publicly available in their supplementary materials. The datasets used for secondary analysis can be accessed through the publication and relevant repositories, subject to any access restrictions specified by the original authors.

## Ethics Statement

This article is based on a secondary analysis of data originally reported by Giovanniello et al. [5]. As this is a secondary data analysis, no new ethical approval or animal research was conducted for this study. All experimental procedures involving animals were performed in accordance with the ethical standards established by the original study.

## Statements

All authors have read and approved the final version of the manuscript. Feng Lin had access to the publicly available data used in this study and takes full responsibility for the integrity of the data analysis.

