## Supplementary material for "Stress-Associated Alterations in Amygdala-Striatal Activity: A Multi-Level Analysis of Distributional, Dynamical, and Computational Signatures": 2026-06-12

### A Single Variable Model Constructed by Ridged Regression

The variable $x$ in this section represents either $x_{BLA}$ or $x_{CeA}$ in the main text. Because the reconstructed system exhibits a single stable equilibrium, the curvature of the effective potential near the fixed point provides a geometric interpretation of the local stability quantified by the Jacobian eigenvalues. The experiments can be viewed as perturbations to the system. Before the perturbation, the signals are in a steady state; after the perturbation, the system returns to a steady state in the vicinity of the pre-perturbation state. In addition, calcium indicators introduce intrinsic temporal filtering, which can effectively generate higher-order temporal dynamics in the observed signals. This process was approximated using a second-order phenomenological dynamical model in which the observed acceleration was expressed as a function of signal amplitude and its first derivative:

$$\ddot{x} = a_0 + a_1 x + a_2 \dot{x} + a_3 x^2 + a_4 \dot{x}^2 + a_5 x \dot{x} + \text{higher-order terms} \tag{A.1}$$

In this equation, $x$ represents the signal recorded in the experimental data, and $\dot{x}$ can be estimated as the difference between consecutive data points, i.e., $\dot{x}_t \approx (x_{t+1} - x_t)/\Delta t$. The acceleration $\ddot{x}$ takes the form of a damping recovery equation. To construct this model, we apply ridge regression (python sklearn library, Ridge packet). The higher-order terms account for nonlinearities but contribute less significantly to the overall dynamics.

Because the reconstructed trajectories typically approached a stable post-perturbation state following stimulus onset, we sought to characterize the observed recovery dynamics using a low-dimensional phenomenological model. The experimental paradigm can be viewed as a transient perturbation to an underlying dynamical process: signals initially fluctuate away from baseline and subsequently evolve toward a new quasi-stationary state. In addition, calcium indicators introduce temporal filtering that may generate effective higher-order temporal structure in the recorded signals. To capture these empirical dynamics, we approximated the observed acceleration as a function of signal amplitude and its temporal derivative using the following second-order phenomenological equation (A.1):

$$\begin{aligned} \dot{x} &= v \\ \dot{v} &= a_0 + a_1 x + a_2 v + a_3 x^2 + a_4 v^2 + a_5 x v \end{aligned} \tag{A.2}$$

The coefficients in (A.1) were estimated using ridge regression and should be interpreted as parameters of a fitted phenomenological model rather than direct physical or biological quantities. When the nonlinear terms vanish, the model reduces to a linear second-order dynamical system whose local behavior can be characterized analytically through its Jacobian eigenvalues.

Within the reconstructed dynamical approximation, local behavior around the estimated equilibrium is characterized by the eigenvalues of the Jacobian matrix. Negative real parts correspond to trajectories that converge toward the equilibrium in the fitted model. These eigenvalues therefore provide a compact summary of local stability in the reconstructed dynamics, although they should not be interpreted as direct measurements of biological stability. The Jacobian ($J$) is given by:

$$J = \begin{pmatrix} 0 & 1 \\ a_1 + 2a_3 x + a_5 v & a_2 + 2a_4 v + a_5 x \end{pmatrix}_{x = x_s, v = \dot{x}_s} \tag{A.3}$$

In this equation, the steady-state value $x_s$ was estimated as the mean signal during the final 5 seconds of the recording window. Because the experimental recordings span 10 seconds following perturbation onset, neural activity typically stabilizes during the latter half of this interval. The final 5-second segment was therefore used as an empirical approximation of the post-perturbation steady state, providing a sufficient number of samples to obtain a stable estimate of the equilibrium signal level. So $\dot{x}_s \approx 0$. Therefore, the Jacobian simplifies to:

$$J = \begin{pmatrix} 0 & 1 \\ a_1 + 2a_3x & a_2 + a_5x \end{pmatrix}_{x=x_s} \tag{A.4}$$

The stability of the system around the steady state is determined by the eigenvalues of the Jacobian. If the real parts of all the eigenvalues are negative, the system is stable around the steady state. This simplified model neglects any oscillations or fluctuations around the steady state.

Because the full dynamical system contains velocity-dependent terms, a strict conservative potential does not formally exist. Therefore, we constructed a heuristic quasi-potential representation for visualization purposes by integrating only the position-dependent terms of the fitted model $U(x)$. This effective potential is a phenomenological and geometric construct derived from the fitted dynamics, and is not intended to represent physical energy or biological cost. By integrating the conservative force terms (those dependent only on position $x$), the effective potential function was defined as:

$$U(x) = -\int (a_0 + a_1x + a_3x^2)dx = -(a_0x + \frac{1}{2}a_1x^2 + \frac{1}{3}a_3x^3) \tag{A.5}$$

where $U(x)$ provides a qualitative geometric summary of the system. In this representation, steeper local curvature corresponds to larger position-dependent restoring components in the fitted dynamics. This visualization is intended as a descriptive summary and should not be interpreted as a physical energy landscape. The phase portraits were constructed in the $(x, \dot{x})$ phase space by visualizing the vector field $(\dot{x}, \ddot{x})$ to illustrate the state-space trajectories and damping characteristics. All stability estimates and landscape profiles were reported with 95% confidence intervals (CIs) derived from the bootstrap distribution.

### B Population-level Model for Pathway Associations

The authors did not specifically point out whether the signals from the two pathways in the same group were observed simultaneously. However, since the number of mice involved in each pathway are different, we can deduce that the signals were not observed synchronously. Thus, causality analysis by transfer entropy, which relies on synchronous time-series data, cannot be applied. To explore whether reproducible cross-pathway statistical associations are present under identical experimental conditions, we constructed a phenomenological two-variable model that includes cross-terms between BLA-DMS and CeA-DMS activity. Because recordings were obtained from different animals and were not temporally synchronized, the model is not intended to infer causal interactions or direct neural coupling. Instead, it provides a population-level description of how activity patterns in one pathway statistically relate to those observed in the other pathway under the same behavioral condition.

Our approach is based on the assumption that, although the data from different mice are not temporally synchronized, the underlying dynamical rules and interaction patterns of the neural circuits are reproducible under identical experimental conditions. The Ridge Regression statistical association model we use is designed to characterize reproducible cross-pathway statistical structure.

This population-level model is an extension of the model (A.1). Let $x_{BLA}$ represent the signal detected by the BLA-DMS pathway and $x_{CeA}$ the signal from the CeA-DMS pathway. The model accounts for both the self-dynamics and population-level cross-dynamics between the two pathways. The equations for each pathway are:

$$\begin{aligned} \ddot{x}_{BLA} &= a_0 + a_1 x_{BLA} + a_2 \dot{x}_{BLA} + a_3 x_{BLA}^2 + a_4 \dot{x}_{BLA}^2 + a_5 x_{BLA} \dot{x}_{BLA} \\ &\quad + b_1 x_{CeA} + b_2 \dot{x}_{CeA} + b_3 x_{BLA} x_{CeA} + b_4 x_{BLA} \dot{x}_{CeA} + b_5 \dot{x}_{BLA} x_{CeA} + b_6 \dot{x}_{BLA} \dot{x}_{CeA} \\ &\quad + \text{higher-order terms} \end{aligned} \tag{A.6}$$

$$\begin{aligned} \ddot{x}_{CeA} &= c_0 + c_1 x_{CeA} + c_2 \dot{x}_{CeA} + c_3 x_{CeA}^2 + c_4 \dot{x}_{CeA}^2 + c_5 x_{CeA} \dot{x}_{CeA} \\ &\quad + d_1 x_{BLA} + d_2 \dot{x}_{BLA} + d_3 x_{BLA} x_{CeA} + d_4 x_{CeA} \dot{x}_{BLA} + d_5 \dot{x}_{CeA} x_{BLA} + d_6 \dot{x}_{BLA} \dot{x}_{CeA} \\ &\quad + \text{higher-order terms} \end{aligned} \tag{A.7}$$

The coefficients $a$ and $c$ (from 0 to 5) describe the reconstructed within-pathway dynamics for the BLA-DMS and CeA-DMS signals, respectively, whereas the coefficients $b$ and $d$ (from 1 to 6) quantify cross-pathway terms included in the phenomenological model. Because the two pathways were recorded in different animals and were not measured simultaneously, these cross-pathway terms should be interpreted as population-level statistical associations under matched experimental conditions rather than direct measures of causal interactions or neural coupling. The model incorporates both within-pathway and cross-pathway components to characterize reproducible patterns in the reconstructed dynamics.

This model was developed as an exploratory analytical framework rather than a predictive or mechanistic model. Its primary purpose is to characterize how the fitted cross-pathway association terms vary across experimental conditions (e.g., Control versus Stress) and behavioral paradigms. Accordingly, changes in the coefficients $b$ and $d$ are interpreted as changes in the statistical structure of the reconstructed phenomenological model and not as direct evidence of altered causal interactions between neural circuits.

To characterize local stability properties of the reconstructed phenomenological model Eq. (A.6) and (A.7), we linearized the system around its empirical equilibrium $\mathbf{u}^* = [x_{BLA}^*, v_{BLA}^*, x_{CeA}^*, v_{CeA}^*]^T$, where $v^* = \dot{x} = 0$ at steady state. The global stability is governed by the $4 \times 4$ Jacobian matrix $\mathbf{J}$. Because the model was fit to population-averaged recordings obtained from different animals, the resulting Jacobian describes local properties of the reconstructed phenomenological representation and should not be interpreted as a direct estimate of circuit-level interactions.

$$\mathbf{J} = \begin{bmatrix} 0 & 1 & 0 & 0 \\ \frac{\partial \ddot{x}_{BLA}}{\partial x_{BLA}} & \frac{\partial \ddot{x}_{BLA}}{\partial \dot{x}_{BLA}} & \frac{\partial \ddot{x}_{BLA}}{\partial x_{CeA}} & \frac{\partial \ddot{x}_{BLA}}{\partial \dot{x}_{CeA}} \\ 0 & 0 & 0 & 1 \\ \frac{\partial \ddot{x}_{CeA}}{\partial x_{BLA}} & \frac{\partial \ddot{x}_{CeA}}{\partial \dot{x}_{BLA}} & \frac{\partial \ddot{x}_{CeA}}{\partial x_{CeA}} & \frac{\partial \ddot{x}_{CeA}}{\partial \dot{x}_{CeA}} \end{bmatrix}_{\mathbf{u}^*} \tag{A.8}$$

By differentiating the cross-pathway equations, we obtain the explicit forms of the partial derivatives. Position-dependent local sensitivity of BLA:

$$\begin{aligned} \frac{\partial \ddot{x}_{BLA}}{\partial x_{BLA}} &= a_1 + 2a_3 x^*_{BLA} + a_5 v^*_{BLA} + b_3 x^*_{CeA} + b_4 v^*_{CeA} \\ &= a_1 + 2a_3 x^*_{BLA} + b_3 x^*_{CeA} \end{aligned} \tag{A.9}$$

Rate-dependent local sensitivity of BLA:

$$\begin{aligned} \frac{\partial \ddot{x}_{BLA}}{\partial \dot{x}_{BLA}} &= a_2 + 2a_4 v^*_{BLA} + a_5 x^*_{BLA} + b_5 x^*_{CeA} + b_6 v^*_{CeA} \\ &= a_2 + a_5 x^*_{BLA} + b_5 x^*_{CeA} \end{aligned} \tag{A.10}$$

Cross-pathway position/rate dependence from CeA:

$$\begin{aligned} \frac{\partial \ddot{x}_{BLA}}{\partial x_{CeA}} &= b_1 + b_3 x^*_{BLA} + b_5 v^*_{BLA} \\ &= b_1 + b_3 x^*_{BLA} \end{aligned} \tag{A.11}$$

$$\begin{aligned} \frac{\partial \ddot{x}_{BLA}}{\partial \dot{x}_{CeA}} &= b_2 + b_4 x^*_{BLA} + b_6 v^*_{BLA} \\ &= b_2 + b_4 x^*_{BLA} \end{aligned} \tag{A.12}$$

Position-dependent local sensitivity of CeA:

$$\begin{aligned} \frac{\partial \ddot{x}_{CeA}}{\partial x_{CeA}} &= c_1 + 2c_3 x^*_{CeA} + c_5 v^*_{CeA} + d_3 x^*_{BLA} + d_4 v^*_{BLA} \\ &= c_1 + 2c_3 x^*_{CeA} + d_3 x^*_{BLA} \end{aligned} \tag{A.13}$$

Rate-dependent local sensitivity of CeA:

$$\begin{aligned} \frac{\partial \ddot{x}_{CeA}}{\partial \dot{x}_{CeA}} &= c_2 + 2c_4 v^*_{CeA} + c_5 x^*_{CeA} + d_5 x^*_{BLA} + d_6 v^*_{BLA} \\ &= c_2 + c_5 x^*_{CeA} + d_5 x^*_{BLA} \end{aligned} \tag{A.14}$$

Cross-pathway position/rate dependence from BLA:

$$\begin{aligned} \frac{\partial \ddot{x}_{CeA}}{\partial x_{BLA}} &= d_1 + d_3 x^*_{CeA} + d_5 v^*_{CeA} \\ &= d_1 + d_3 x^*_{CeA} \end{aligned} \tag{A.15}$$

$$\begin{aligned} \frac{\partial \ddot{x}_{CeA}}{\partial \dot{x}_{BLA}} &= d_2 + d_4 x^*_{CeA} + d_6 v^*_{CeA} \\ &= d_2 + d_4 x^*_{CeA} \end{aligned} \tag{A.16}$$

Constant terms contribute to the equilibrium location but do not enter the Jacobian because their derivatives vanish.

Within the fitted phenomenological model, local stability and transient behavior are characterized through the combined contribution of position-dependent, rate-dependent, nonlinear, and cross-pathway terms through their effects on the Jacobian matrix.

The sign and statistical significance of individual coefficients were evaluated separately for each experimental condition and are reported in the corresponding Results and Supplementary Tables.

The Jacobian matrix provides a local linear approximation of the reconstructed dynamics around the estimated equilibrium. Eigenvalues of the Jacobian were therefore used as summary measures of local stability within the fitted model. Because the model was derived from ridge-regression reconstruction of non-simultaneous recordings rather than mechanistic system identification, these quantities should be interpreted as descriptors of the reconstructed dynamical representation rather than direct measurements of biological stability, causal interactions, or circuit-level coupling. Accordingly, Jacobian-based analyses were used to compare relative changes in the reconstructed dynamics across experimental conditions rather than to infer mechanistic properties of the underlying neural circuitry.

### C Neural Network Model Architecture

The artificial neural network (ANN) model was constructed as a conceptual computational framework rather than a biologically realistic neural circuit model. The model consists of two interacting LSTM-based modules, denoted as $x$ and $y$. These modules were designed to represent two functionally distinct dynamical subsystems motivated by the experimental observations. Throughout this section, the labels "BLA-DMS" and "CeA-DMS" are used as convenient shorthand for these computational modules and do not imply anatomical equivalence, direct circuit mapping, or biological validation.

LSTM units are selected because their gated cell state naturally implements a controllable memory process, allowing the model to represent persistent latent deviations that accumulate over time following perturbations. These features enable the network to capture the temporal dependencies and persistence of neural activity. When subjected to external perturbations, the network outputs ($x(t)$ and $y(t)$) exhibit an instantaneous increase (response) followed by an exponential decay (returning to baseline), which aligns with the temporal dynamics observed in real neural pathways.

The $x$ and $y$ modules were assigned different computational roles. The $x$ module was designed to be more sensitive to incoming perturbations, whereas the $y$ module was designed to maintain longer-lived internal states and provide stabilizing feedback. These asymmetries were introduced as modeling hypotheses motivated by the phenomenological differences observed in the experimental recordings. They should not be interpreted as direct evidence that the corresponding biological pathways possess identical computational architectures.

Main components of the neural network model are shown in Table C1. There are 7 statistical features of the input signal and target signals to gain "outside" the neural network model. The seven statistical features include mean, standard deviation, skewness, kurtosis, and three difference metrics between input and target signals. The first four features are used for the parameter predictor to judge whether the system is exposed in chronic stress environment. The statistical difference of input signals and the target signals are taking into account by the neural network to judge whether the optimization reaches the aim.

Table C1: Model Components

| Components | Class | Input Dimension | Hidden Dimension | Output Dimension | Key Functions |
|---|---|---|---|---|---|
| Parameter Predictor | Feedforward Neural Networks | 7 (statistical feature) | 64 | 6 ($\phi$) | Predict system's macroscopic parameters |
| $x$ (BLA-DMS) | Adaptive LSTM Predictor | 1 (signal) | 32 | 1 (output) | exploration_bias = 0.2 |
| $y$ CeA-DMS | Adaptive LSTM Predictor | 1 (signal) | 32 | 1 (output) | exploration_bias = $-1$ |
| $x/y$ LSTM core | LSTM | 1 | 32 | 32 (hidden) | Predict internal state of baseline signal |
| $x/y$ predictor | Linear Layer | 32 (hidden) | - | 1 | Predict baseline signal |
| $x/y$ threshold net | Feedforward Neural Networks | 33 (hidden+1) | 16 | 1 (base threshold) | Predict baseline of dynamical threshold |
| $x/y$ weight net | Feedforward Neural Networks | 33 (hidden+1) | 16 | 1 (base weight) | Predict baseline weights of bias signal |

The macroscopic parameters ($\phi$) are used to represent the interaction scale and nonlinearity of the system consisted by $x$ and $y$ nets, the decay modulation of $x$ and $y$ nets, the excitation and inhibition of $y$ to $x$ net. These parameters are predicted by parameter predictor. The key algorithm of this component is in Algorithm 1. The input variables are from the statistical feature component which forms a $R^7$ Euclidean space. By mapping of parameter predictor the output variables are 6, forms a $R^6$. These low-order statistical moments are chosen as compact descriptors of environmental volatility, allowing the model to adapt its internal control parameters according to coarse statistical properties of the training data.

```
Parameter_Predictor
// 1. MLP Forward propagation
Raw_Params ← MLP(S) // R^7 → R^6
// 2. Normalization (Sigmoid)
Scaled_Params ← Sigmoid(Raw_Params)
// 3. Map to a bounded interval (φ)
φ_scale ← Scaled_Params[0] * 1.7 + 0.3 // interaction scale ∈ [0.3, 2.0]
φ_power ← Scaled_Params[1] * 1.0 + 0.5 // suppression power ∈ [0.5, 1.5]
φ_dec1 ← Scaled_Params[2] * 1.7 + 0.3 // decay modulation of x net ∈ [0.3, 2.0]
φ_dec2 ← Scaled_Params[3] * 1.7 + 0.3 // decay modulation of y net ∈ [0.3, 2.0]
φ_exc ← Scaled_Params[4] // excitation of y to x ∈ [0.0, 1.0]
φ_inh ← Scaled_Params[5] // inhibition of y of x ∈ [0.0, 1.0]
Return φ
```

**Algorithm 1:** Parameter predictor

The adaptive LSTM predictor is used to determine the inner self-adaptive mechanisms of $x$ and $y$ nets. The key point is to dynamically calculate the deviation. In Algorithm 2 the input values of adaptive LSTM predictor at time $t$ is represented by $I_t$, $H_{t-1}$ represents the hidden state at time $t-1$. The accumulated deviations are represented by $\delta_t$. The output of adaptive LSTM predictor is the predicted value $P_t$ at time $t$, the contribution of deviation $\Delta\delta$, $H_t$. The information is used to dictate the system.

```
Adaptive_LSTM_Predictor(I_t, H_{t-1}, Bias_k)
// 1. LSTM calculation
H_t ← LSTM_Core(I_t, H_{t-1})
P_t ← Predictor_Linear(H_t) // calibration prediction
// 2. Calculation of dynamical parameter
Features ← Concat(H_t, I_t)
Base_Threshold ← Threshold_Net(Features)
Base_Weight ← Weight_Net(Features)
// 3. Application exploration bias (Bias_k)
Dynamic_Threshold ← Base_Threshold · exp(−Bias_k)
Deviation_Weight ← Base_Weight · (1.0 + Bias_k · 0.5)
// 4. Calculation of contribution of deviation
E_t ← I_t − P_t
|E_t| ← Absolute(E_t)
// 5. Soft Mask to determine the stimulating degree
Soft_Mask ← Sigmoid((|E_t| - Dynamic_Threshold) · 5.0)
// 6. Final contribution of deviation
Δδ ← E_t · Deviation_Weight · Soft_Mask
Return P_t, Δδ, H_t
```

**Algorithm 2:** Adaptive LSTM Predictor

Though the coordination of $x$ and $y$ is mainly determined by Parameter_Predictor, the output of macroscopic parameters $\phi$ (for instance $\phi_{\text{scale}}, \phi_{\text{inh}}$), Meta_Controller in Algorithm 3 is responsible for the adaptation based on instant states. Meta_Controller accepts the instant deviations $(\delta_x, \delta_y)$ of $x$ and $y$ and threshold information to predict the instant interaction scale and decay modulations. Its output signals, Uncertainty (uncertainty), are used to dynamically weighting the coordination terms of $x$ and $y$. This makes the system's dynamics not only depending on the macroscopic environment (statistical features), but also the microscopic environment which makes the "brain" control its dynamical behavior in a refined manner.

Meta_Controller
// 1. input is a combined state
Combined_State ← Concat($\delta_x, \delta_y, P_x, P_y, \text{Threshold}_x, \text{Threshold}_y$)
// 2. Calculation of uncertainty (core signal)
Uncertainty ← Sigmoid(FNN_Uncertainty(Combined_State)) // [0, 1]
// 3. Control parameters estimation based on uncertainty
Interaction_Input ← Concat(Uncertainty, Combined_State)
Control_Params ← FNN_Interaction(Interaction_Input) // $R^4$
// 4. Mapping to control signals
Decay_$x$ ← Sigmoid(Control_Params[0]) · 0.3 + 0.01 // Instant decay modulation of $x$
Interact$_x$ ← Sigmoid(Control_Params[1]) · Uncertainty // interaction scale of $x$
Decay$_y$ ← Sigmoid(Control_Params[2]) · 0.3 + 0.01 // Instant decay modulation of $y$
Interact$_y$ ← Sigmoid(Control_Params[3]) · (1.0 - Uncertainty · 0.5) // interaction scale of $y$
// 5. Returned Control signals
Return Decay_$x$, Interact$_x$, Decay$_y$, Interact$_y$, Uncertainty

**Algorithm 3:** Meta Controller

The Algorithm in 4 is to exhibit in each time step $t$, how the macroscopic parameters $\phi$ to adjust the coordination of $x$ and $y$ and how the updating of $\delta$. Input: $I_t$, Targets $T_t$. Variables: $\phi$ at time $t$, $\delta_x, \delta_y$ (accumulated deviations) $P_x, P_y$ (predicted calibration).

Unified_Control_System($I_t, \phi, \delta_x, \delta_y$)
// 1. LSTM prediction (By Algorithm 2)
$P_1, \Delta\delta_1, H_1$ ← Adaptive_LSTM_Predictor($I_t$, Bias$_x$)
$P_2, \Delta\delta_2, H_2$ ← Adaptive_LSTM_Predictor($I_t$, Bias$_y$)
// 2. Calculation of coordination term (using macroscopic parameters $\phi$)
$\phi_{\text{scale}}, \phi_{\text{power}}, \ldots, \phi_{\text{inh}} \leftarrow \phi$
// $y \to x$ signal (excitation and inhibition)
Signal$_{y\to x}$ ← Sign($\delta_y$) · Power($|\delta_y|$, $\phi_{\text{power}}$) · $\phi_{\text{scale}} \cdot \phi_{\text{exc}}$ - Sign($\delta_x$) · Power($|\delta_x|$, $\phi_{\text{power}}$) · $\phi_{\text{scale}} \cdot \phi_{\text{inh}}$
// cost of inhibition
Cost$_{\text{2Supp}}$ ← Absolute(Signal$_{y\to x,\text{Inh}}$) · 2.0
// $y \to x$ signal (difference shirk)
Signal$_{x\to y} \leftarrow \delta_x \cdot \phi_{\text{scale}} \cdot 0.1 \cdot (1.0 + \text{Capacity}_x)$
// 3. Decay of difference (Decay)
$\delta_x^{\text{decay}} \leftarrow \delta_x \cdot (x - \text{Decay}_{\text{Meta}} \cdot \phi_{\text{decx}})$ // Decay$_{\text{Meta}}$ from Algorithm 3
$\delta_y^{\text{decay}} \leftarrow \delta_y \cdot (x - \text{Decay}_{\text{Meta}} \cdot \phi_{\text{decy}})$
// 4. Updating of deviation (Key point: cost and coordination terms are added)
$\delta_x^{\text{new}} \leftarrow \delta_x^{\text{decay}} + \Delta\delta_x + \text{Signal}_{y\to x}$
$\delta_y^{\text{new}} \leftarrow \delta_y^{\text{decay}} + \Delta\delta_y + \text{Signal}_{x\to y} + \text{Cost}_{y\text{Supp}}$
// 5. Capacity limitation and punishment (hard constraint)
Penalty$_x$ ← ReLU($|\delta_x^{\text{new}}| - 0.2$)
Penalty$_y$ ← ReLU($|\delta_y^{\text{new}}| - 3.0$)
$\delta_x$ ← Clamp($\delta_x^{\text{new}}$, [-0.2, 0.2])
$\delta_y$ ← Clamp($\delta_y^{\text{new}}$, [-3.0, 3.0])
// 6. Output
Output$_x \leftarrow P_x + \delta_x$
Output$_y \leftarrow P_y + \delta_y$
Return Output$_x$, Output$_y$, Penalty$_x$, Penalty$_y$

**Algorithm 4:** Unified control system

The loss function (driving force of function differentiation)
In formation of key loss terms are exhibited in Table C2

Table C2: Key loss terms

| Loss terms | Aim | Calculation |
|---|---|---|
| $L_{\text{Buffer}}$ | Reward | Deviations of $y$ under perturbation to buffer $x$. |
| $L_{\text{Calibration}}$ | Guarantee $y$ is calibrated baseline when there is no perturbation | $L_{\text{Calibration}} = \text{MSE}(O_2 \odot \mathbf{M}_{\text{Non-Pert}}, T \odot \mathbf{M}_{\text{Non-Pert}})$ |
| $L_{\text{Saturation}}$ | punishment | deviation of $x$ that out of capacity. $L_{\text{Saturation}} = \text{Mean}(\text{Penalty}_x)$ |
| $L_{\text{Persistence}}$ | reward | persistence of $y$ $\delta$ (negative weights). |

The loss function is the key point of driving the network $x$ and $y$'s function differentiation. $L_{Total}$ is the weighted addition of all loss functions. Here $\mathbf{M}_{\text{Pert}}$ is the masked perturbation. $\mathbb{I}(|I_t - T_t| > 0.1)$ and $\mathbf{M}_{\text{Non-Pert}}$ is the non-masked perturbation in which $1 - \mathbf{M}_{\text{Pert}}$ and $W$ is the weight. The total weighted loss function is

$$L_{Total} = L_x + L_y$$

$$L_x = W_{\text{MSE}} \cdot L_{\text{MSE}}(O_1, T) + W_{\text{Sat}} \cdot L_{\text{Saturation}}$$

$$L_y = W_{\text{Buf}} \cdot L_{\text{Buffer}} + W_{\text{Cal}} \cdot L_{\text{Calibration}} + W_{\text{Inst}} \cdot L_{\text{Instability}} + W_{\text{Pers}} \cdot L_{\text{Persistence}}$$

$$L_{\text{Buffer}} = -\frac{1}{|\mathcal{T}|} \sum_{t \in \mathcal{T}} (|O_{x,t} - O_{y,t}| \cdot \mathbf{M}_{\text{Pert},t})$$

$O_{x,t}$ and $O_{y,t}$ are the output of $x$ and $y$ at time $t$. $|\mathcal{T}|$ is the total time step. $\mathbf{M}_{\text{Pert},t}$ is the masked perturbation, when $|I_t - T_t| > \theta_{\text{pert}} = 1 \quad or \quad 0.$(in our code $\theta_{\text{pert}} = 0.1$)

$$L_{\text{Persistence}} = -\frac{1}{|\mathcal{T}|} \sum_{t \in \mathcal{T}} |\delta_{y,t}|$$

where $\delta_{y,t}$ is the cumulated deviations of $y$ at time $t$. The weights $W_{\text{Pers}}$ in our code is negative $(-2.0)$, meaning that minimizing this term is equivalent to maxmize $|\delta_{y,t}|$.

#### C.1 Robustness Test for ANN

To ensure the statistical robustness and reproducibility of the mechanisms learned by the Unified Control System, we conducted a stability analysis using $N = 10$ independent random seeds (specifically, seeds $0, 10, 20, \ldots, 90$).

Table C3 presents the statistical characteristics (Mean, Standard Deviation, Range, and Coefficient of Variation) for both the learned macro-parameters (e.g., Suppression Power, Interaction Scale, etc) and the functional response metrics (e.g., Recovery Time, Max N1 Deviation, etc).

The distribution and variability of the learned parameters and functional metrics listed in Table C3 are visually represented by the box plots in Figure C1. Our stability analysis provides insight into the trade-off between stability and flexibility enforced by chronic stress. While the Stress models achieved near-zero CV for Recovery Time, confirming a highly rigid, robust strategy to perturbation, the Control models retained a higher functional CV (e.g., 25.8% CV in Recovery Time). This elevated functional variance in the Control group may be interpreted as a higher degree of behavioral or functional flexibility in response to acute challenges. Furthermore, the highest CVs were observed in the inter-pathway connection weights (CeA-DMS $\rightarrow$ BLA-DMS coordination), indicating that the system preserves structural flexibility (functional compensation) as a core mechanism for achieving functional stability. Within this computational framework, the coexistence of stable outputs and variable internal coordination parameters suggests that the model favors a balance between output robustness (for absolute stability CV $\ll 0.01$) and internal flexibility.

Table C3: Statistical Robustness of Learned Parameters and Functional Metrics Across 10 Independent Seeds

| | | Control | | | | Stress | | | |
|---|---|---|---|---|---|---|---|---|---|
| | Parameters | Mean | Standard Error | Range | Coefficient of Variation | Mean | Standard Error | Range | Coefficient of Variation |
| Training Index | final loss | 0.001507 | 0.000282 | [0.001107, 0.002091] | 0.1871 | 0.003581 | 0.000477 | [0.002855, 0.004457] | 0.1333 |
| | mse | 0.000701 | 0.000139 | [0.000490, 0.000987] | 0.1979 | 0.002735 | 0.000201 | [0.002381, 0.003043] | 0.0735 |
| Learned Parameters | interactions cale | 0.5186 | 0.063 | [0.4094, 0.6249] | 0.1214 | 0.5182 | 0.0531 | [0.4407, 0.6254] | 0.1024 |
| | suppression power | 1.2318 | 0.0795 | [1.0965, 1.3718] | 0.0645 | 1.246 | 0.0591 | [1.1563, 1.3519] | 0.0474 |
| | BLA-DMS decay modulation | 1.7571 | 0.087 | [1.5809, 1.8989] | 0.0495 | 1.5451 | 0.2172 | [1.0864, 1.7548] | 0.1405 |
| | CeA-DMS decay modulation | 1.7591 | 0.0421 | [1.7027, 1.8405] | 0.0239 | 1.7429 | 0.0599 | [1.6647, 1.8516] | 0.0344 |
| | CeA-DMS $\rightarrow$ BLA-DMS excitatory | 0.3634 | 0.0896 | [0.1933, 0.5055] | 0.2465 | 0.3527 | 0.0987 | [0.1995, 0.5364] | 0.2798 |
| | CeA-DMS $\rightarrow$ BLA-DMS inhibitory | 0.1234 | 0.0501 | [0.0529, 0.2376] | 0.4062 | 0.1387 | 0.0468 | [0.0804, 0.2549] | 0.3375 |
| Response to Perturbation | recovery time | 3 | 0.7746 | [2.0000, 4.0000] | 0.2582 | 1 | 0 | [1.0000, 1.0000] | 0 |
| | max BLA-DMS deviation | 0.3389 | 0.0688 | [0.2373, 0.4416] | 0.203 | 0.057 | 0.0089 | [0.0438, 0.0715] | 0.1558 |
| | max CeA-DMS deviation | 0.3389 | 0.0953 | [0.1729, 0.4691] | 0.2813 | 0.6698 | 0.0722 | [0.5650, 0.7655] | 0.1077 |
| | steady state error | 0.0564 | 0.0041 | [0.0510, 0.0619] | 0.0724 | 0.0512 | 0.0037 | [0.0459, 0.0567] | 0.0718 |

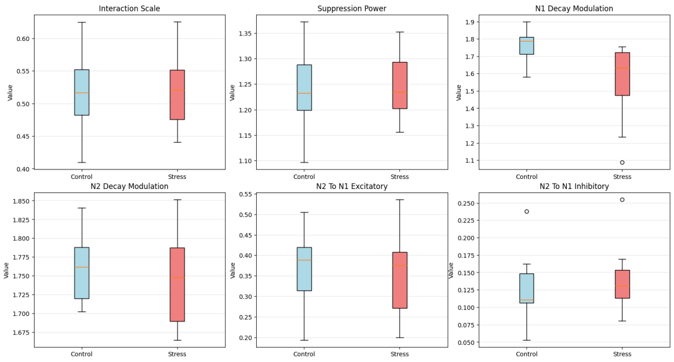


(a) Learned parameters

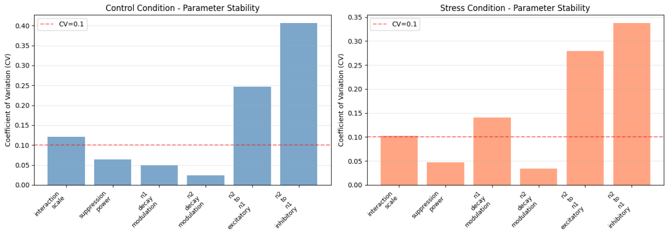


(b) Coefficient of Variation

Figure C1: Robustness test for the ANN model.

#### C.2 Illustrative Comparison of Symmetric and Asymmetric Objective Functions

The ANN framework was not intended as an independent validation of the dynamical analyses presented in the main text. Instead, it serves as a proof-of-concept computational illustration showing that distinct dynamical response patterns qualitatively resembling the experimental observations can emerge when different adaptive objectives are imposed on interacting computational modules. Because the model is hypothesis-driven, several architectural and objective-function asymmetries were deliberately specified a priori. These assumptions should therefore be viewed as explicit encodings of the proposed mechanism rather than discoveries emerging from the optimization process itself.

To investigate whether the observed valence-specific dynamics could emerge from a simpler, non-specific architecture, we conducted a control simulation using a Unified Loss Framework. In this configuration, both the BLA-DMS and CeA-DMS units were optimized using an identical objective function:The neural network architecture consists of two interacting units, $x$ (representing the BLA-DMS pathway) and $y$ (representing the CeA-DMS pathway). The state of each unit is defined by its cumulative informational deviation $\delta_i$, which evolves according to the following coupled stochastic differential logic:

For the $x$ Unit (Controlled Pathway):

$$\delta_x(t) = \delta_x(t-1) + \Delta_x(t) + \Phi(t) \tag{A.17}$$

Where $\Phi(t)$ is the active inhibitory control signal from $y$ designed to counteract excessive fluctuations in $x$:

$$\Phi(t) = -\text{sgn}(\delta_x) \cdot |\delta_x|^\gamma \cdot (\theta_{scale} \cdot \lambda_{x,t}) \cdot (1 + \tanh(2|\delta_y|)) \tag{A.18}$$

Here, $\gamma$ is the suppression power, $\theta_{scale}$ is the global interaction scale, and $\lambda_{x,t}$ is the dynamic gain provided by the meta-controller.

For the $y$ Unit (Buffering Pathway):

$$\delta_y(t) = \delta_y(t-1) + \Delta_y(t) + 0.3 \cdot \lambda_{y,t} \cdot \delta_x(t-1) \tag{A.19}$$

The term $0.3 \cdot \lambda_{y,t} \cdot \delta_x$ represents the passive absorption of informational overflow from the BLA to the CeA. Global Dissipation:To ensure physical realism and energy dissipation, both deviations are subjected to a shared decay modulation $\theta_{decay}$:

$$\delta_i(t) \leftarrow \delta_i(t) \cdot (1 - \eta_i \cdot \theta_{decay}), \quad i \in x, y \tag{A.20}$$

Unlike standard black-box models, our system is optimized using a Unified Stability-Constrained Loss. This objective forces the network to find a balance between signal fidelity and dynamical stability:

$$L_{total} = L_{MSE} + \alpha \cdot L_{stability} \tag{A.21}$$

In which $\alpha = 0.1$ is a given parameter. Trajectory Reconstruction (MSE):

$$L_{MSE} = \frac{1}{2T} \sum_{t=1}^{T} \left( |M_x(t) - M_{target}(t)|^2 + |M_y(t) - M_{target}(t)|^2 \right) \tag{A.22}$$

Physical Stability Penalty: The stability term penalizes deviations that exceed a biologically plausible range relative to the input signal magnitude $|P(t)|$:

$$L_{stability} = \frac{1}{T} \sum_{t=1}^{T} \text{ReLU}\left( (|\delta_x(t)| + |\delta_y(t)|) - \Omega \cdot |P(t)| \right) \tag{A.23}$$

Where $\Omega = 2.0$ is the maximum deviation ratio and $\alpha = 0.1$ is the penalty weight.

Data-Driven Parameter Discovery: The parameters $\Theta = \{\theta_{scale}, \gamma, \theta_{decay}\}$ are not learned as static weights. Instead, they are generated by a Parameter Predictor Network $\mathcal{F}$ mapping from 7-dimensional statistical features of the input ensemble:

$$\Theta = \mathcal{F}(\text{Stats}_{ensemble}) \tag{A.24}$$

This ensures that the divergent dynamics (e.g., the higher interaction scale in the Stress group) are emergent properties required to minimize the stability loss under high-volatility conditions.

In Figure C2, under a single perturbation (e.g., footshock), both the Control and Stress groups in the BLA-DMS pathway show stronger responses than observed experimentally. This discrepancy arises because the neural network design oversimplifies the biological mechanisms. Specifically, when BLA ($x$) deviates from baseline, CeA ($y$) generates a reverse signal based on preset feedback parameters:

Negative Feedback: When $x$ deviates, CeA produces a suppressive signal to "pull" $x$ back, with the suppression strength increasing non-linearly as the deviation grows.

Passive Activation of $y$ by $x$: The deviation in BLA's signal directly activates CeA, but this interaction is linear and one-way, meaning the stronger the deviation in BLA, the more CeA gets activated.

Shared Decay Modulation: The model uses a global decay parameter that affects both systems simultaneously, but this fails to account for their individual dynamic behaviors.

Identical MSE Loss Function: The loss function used for training both sub-networks is the same (Mean Squared Error, MSE), which does not account for the potential differences in how these two pathways should adapt to stress.

As a result of these simplifications, the neural network outputs stronger responses than what is seen in the experimental data under chronic stress.

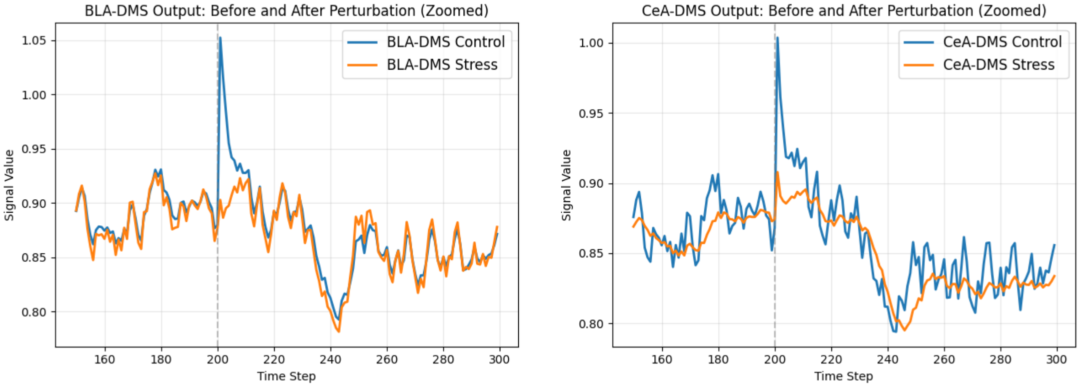


(a) Simulated signals of BLA-DMS sub-network (b) Simulated signals of CeA-DMS sub-network

Figure C2: Simulated subnetwork responses to acute perturbation sharing the Same Loss Function. Three prediction parameters represent subnetworks' coupling. 2(a) The BLA-DMS unit shows high-fidelity tracking in controls but is suppressed in the stress group to prevent saturation. 2(b) The CeA-DMS unit exhibits similar tendency as BLA-DMS unit which fails to reproduce several qualitative features observed in the experimental recordings

If the coupling between the two subnetworks is modified according to the scheme in Algorithm 4, and only the identical loss function as in Equation (A.21) is retained for training, the simulation results, shown in Figure C3, which fails to reproduce several qualitative features observed in the experimental recordings.

Importantly, the ANN framework should be interpreted as an illustrative computational model rather than a validated mechanistic account of the underlying neural circuits. The architecture, coupling rules, and objective functions were specified a priori to explore whether distinct adaptive objectives could generate qualitatively different dynamical regimes. Consequently, successful reproduction of selected experimental features does not constitute evidence that the biological system implements the same computational strategy. The ANN results are therefore used only as a complementary conceptual demonstration alongside the statistical and phenomenological dynamical analyses.

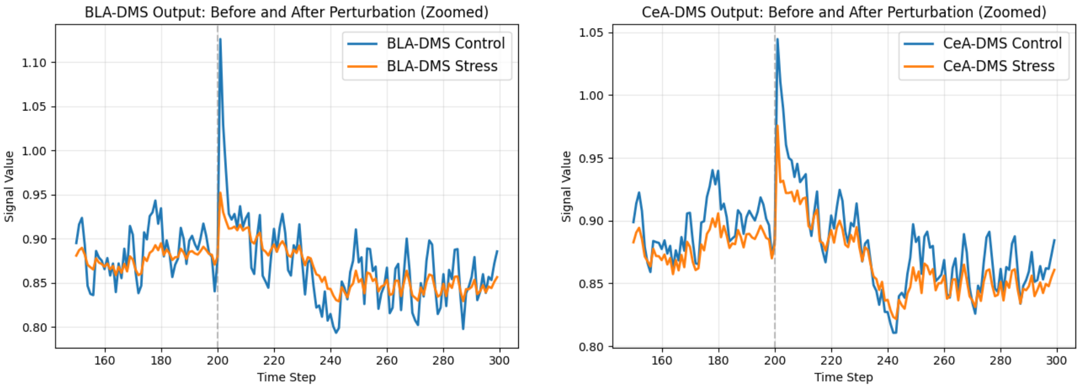


(a) Simulated signals of BLA-DMS sub-network (b) Simulated signals of CeA-DMS sub-network

Figure C3: Simulated subnetwork responses to acute perturbation sharing the Same Loss Function. The subnetworks' coupling are the same as the ANN architecture. 3(a) The BLA-DMS unit shows high-fidelity tracking in controls but is suppressed in the stress group to prevent saturation. 3(b) The CeA-DMS unit exhibits similar tendency as BLA-DMS unit which fails to reproduce several qualitative features observed in the experimental recordings.

### D Supplementary Graphs, Tables and Results

#### D.1 Unpredicted Reward

Learned modality includes task-related events during the operant lever-press paradigm in which mice learned to associate lever pressing with food rewards. Unpredicted rewards were occasionally delivered during the task. Although these rewards were not contingent on the mouse's immediate action, they occurred within a context where mice had already learned lever pressing-reward associations.

##### D.1.1 Statistical Perspective

Figure D4 depicts the temporal evolution of KL divergence following unpredicted reward delivery. Both groups exhibited an initial increase in KL divergence, indicating a transient shift in the signal distribution relative to the pre-event baseline. In the control group, the CeA-DMS pathway (green dashed line) maintained elevated KL divergence for a longer duration during the post-reward period, whereas the stressed group returned more rapidly toward baseline levels. These results suggest that the distributional effects of the unexpected reward persisted longer in the control condition.

Table D4: Statistical Robustness of Integrated KL Divergence Following Unpredicted Reward

| Pathway | Group | Mean Cumulative $D_{KL}$ | 95% CI |
|---|---|---|---|
| BLA-DMS | Control | 11577.6141 | [4737.6294, 23706.7800] |
| BLA-DMS | Stress | 41112.3110 | [14561.3068, 82247.0229] |
| CeA-DMS | Control | 39840.0068 | [10055.5033, 84940.6982] |
| CeA-DMS | Stress | 22554.1092 | [9437.3740, 60992.8522] |

##### D.1.2 Dynamical Perspective

Parallel to the ridge regression analysis for footshock presented in the main text, Table D5 reports the mean coefficients and their corresponding 95% confidence intervals (CIs) derived from 1000 bootstrap iterations

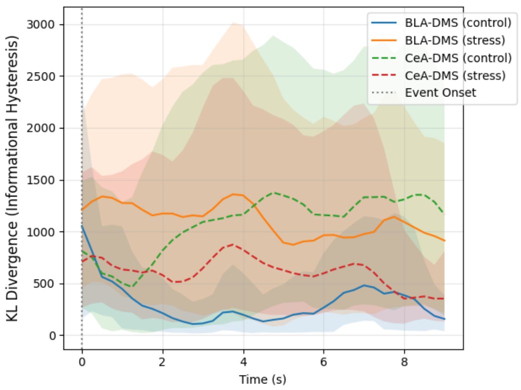


Figure D4: KL divergence of BLA-DMS and CeA-DMS response to unpredicted reward

for the unpredicted reward response.

Table D5: Single Variable Model Statistics of Governing Dynamical Equations for Unpredicted Reward Response

| Pathway | Term | Control Mean | 95% CI | Stress Mean | 95% CI |
|---|---|---|---|---|---|
| BLA-DMS | Intercept | 7.1891 | [ -3.7519, 33.7591] | -14.8515 | [ -32.6979, -3.6618]* |
| | $x_{BLA}$ | -56.5409 | [-150.8085, -15.0722]* | -34.1524 | [ -67.7800, -11.8236]* |
| | $\dot{x}_{BLA}$ | -29.6876 | [ -33.3740, -26.2567]* | -28.0101 | [ -33.3118, -21.6534]* |
| | $x^2_{BLA}$ | 2.2001 | [ -51.2579, 47.9127] | 25.0538 | [ 3.7402, 48.9843]* |
| | $x_{BLA}\dot{x}_{BLA}$ | 1.6282 | [ -10.1931, 13.7740] | 4.3439 | [ -10.3499, 19.9228] |
| | $\dot{x}^2_{BLA}$ | -0.2874 | [ -1.0894, 0.6347] | 0.3357 | [ -0.8460, 1.5980] |
| CeA-DMS | Intercept | -4.2516 | [ -16.0314, 13.0788] | 12.6131 | [ 3.2989, 28.6692]* |
| | $x_{CeA}$ | -54.8347 | [-101.7888, -22.9360]* | -14.4596 | [ -42.7331, -1.7253]* |
| | $\dot{x}_{CeA}$ | -29.6274 | [ -32.8554, -25.6176]* | -25.5327 | [ -32.0388, -19.3126]* |
| | $x^2_{CeA}$ | 8.6733 | [ -24.8204, 35.2159] | -13.2403 | [ -34.1455, 2.1735] |
| | $x_{CeA}\dot{x}_{CeA}$ | 1.5517 | [ -14.8011, 17.6579] | -2.7992 | [ -13.8325, 7.5916] |
| | $\dot{x}^2_{CeA}$ | -0.1288 | [ -1.2116, 1.0444] | -0.3241 | [ -2.0143, 1.2905] |

* Indicates statistical significance (95% CI does not overlap with zero).

Table D6: Statistics of Principal Jacobian Eigenvalues

| Pathway | Group | Mean Principal $Re(E_1)$ | 95% CI |
|---|---|---|---|
| BLA-DMS | Control | -2.3394 | [-6.4732, -0.5823]* |
| BLA-DMS | Stress | -1.9031 | [-3.9257, -0.7885]* |
| CeA-DMS | Control | -2.1398 | [-3.8837, -1.1352]* |
| CeA-DMS | Stress | -1.1651 | [-2.5981, -0.3551]* |

* Indicates statistical significance (95% CI does not overlap with zero).

While the bootstrapped distributions provide a statistical summary of signal variability across the population, the reconstructed dynamical models provide a separate phenomenological description of local trajectory behavior near the estimated equilibrium. As summarized in Table D7, eigenvalues derived from the ridge-regression models were used as descriptive measures of local stability within the reconstructed phase-space representation. These quantities should be interpreted independently from the distributional

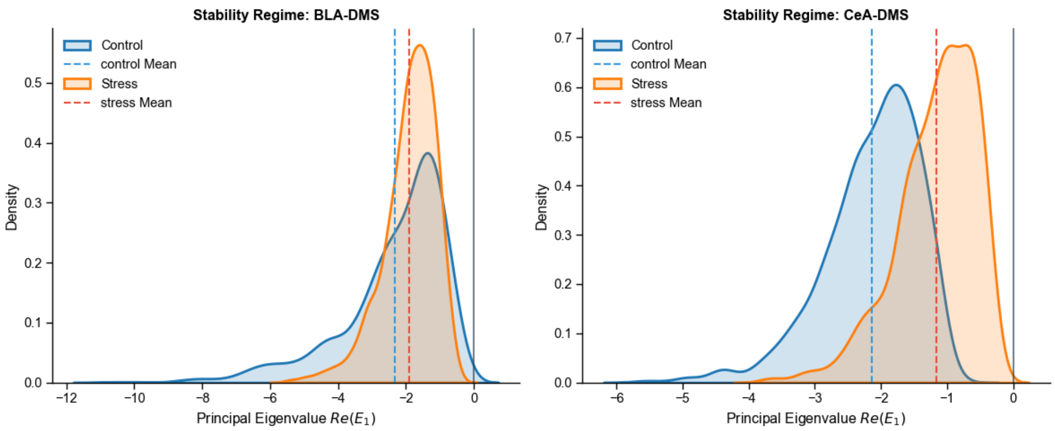


Figure D5: Population-level recovery-rate characteristics analysis of neural trajectories. The probability density functions (PDFs) of the principal Jacobian eigenvalues ($Re(E_1)$) were estimated via subject-level bootstrapping ($n = 1000$ iterations) following unpredicted reward. The consistently negative real parts of the eigenvalues across both pathways confirm the global asymptotic stability of the underlying dynamical regimes.

analyses, as no direct theoretical mapping between the two measures is assumed in the present study.

Table D7: Deterministic Stability Parameters and Recovery Kinetics Post Unpredicted Reward

| Group | Pathway | Steady State | | Eigenvalues | | Total Recovery Speed |
|---|---|---|---|---|---|---|
| | | $x_s$ | $\dot{x}_s$ | $E_1$ | $E_2$ | |
| Control | BLA-DMS | 0.0331 | 0 | -1.5011 | -27.7207 | 27.761 |
| | CeA-DMS | -0.1326 | 0 | -1.7699 | -27.7119 | 27.768 |
| Stress | BLA-DMS | -0.3153 | 0 | -1.7012 | -27.2792 | 27.332 |
| | CeA-DMS | 0.5462 | 0 | -1.0187 | -25.9814 | 26.001 |

Table D8: Mean and Standard Deviation Change for Each Group and Pathway

| Case | Group | Pathway | Baseline Mean (Pre) | Baseline Mean (Post) | ΔMean (Post-Pre) | ΔStd(Post-Pre) |
|---|---|---|---|---|---|---|
| FootShock | Control | BLA-DMS | 0.0252 | -0.397 | -0.4222 | 0.0832 |
| | | CeA-DMS | 0.2392 | -0.3924 | -0.6316 | 0.1804 |
| | Stress | BLA-DMS | 0.2294 | -0.6465 | -0.8759 | 0.0884 |
| | | CeA-DMS | 0.1392 | -0.1597 | -0.2989 | 0.2692 |
| Unpredicted Reward | Control | BLA-DMS | 0.0474 | 0.0331 | -0.0143 | 0.1324 |
| | | CeA-DMS | 0.0849 | -0.1326 | -0.2175 | 0.3418 |
| | Stress | BLA-DMS | 0.1182 | -0.3153 | -0.4335 | 0.2384 |
| | | CeA-DMS | -0.1213 | 0.5462 | 0.6675 | 0.0541 |

The population-level dynamical model of the Unpredicted Reward phase (Tables D9-D10).

#### D.2 FR1 to RR10 Training Sessions

##### D.2.1 KL Divergence: Tables D11 and D12

Table D9: Coefficients of Population-level Regression Model (BLA-DMS as Target pathway)

| Variables | Control Mean Coefficients | 95% CI | Stress Mean Coefficients | 95% CI |
|---|---|---|---|---|
| $x_{BLA}$ | -98.7578 | [-245.0543, -14.1904]* | -25.7685 | [-93.2425, 27.6881] |
| $\dot{x}_{BLA}$ | -31.9542 | [-38.8487, -26.1135]* | -25.7713 | [-38.1226, -11.4908]* |
| $x^2_{BLA}$ | 8.4001 | [-100.0106, 150.9124] | 55.2617 | [-28.3793, 149.3794] |
| $x_{BLA}\dot{x}_{BLA}$ | 4.7084 | [-12.9674, 22.9049] | 3.4384 | [-19.5655, 25.3180] |
| $\dot{x}^2_{BLA}$ | -0.1834 | [-1.1506, 0.6457] | 0.2197 | [-1.1821, 1.4561] |
| $x_{CeA}$ | 28.0601 | [-38.2279, 111.6203] | -18.1017 | [-63.2768, 34.2242] |
| $\dot{x}_{CeA}$ | 1.0346 | [-3.0592, 6.5920] | 1.2395 | [-8.2394, 14.6147] |
| $x_{BLA}x_{CeA}$ | 68.9795 | [-22.1482, 213.8133] | -38.6598 | [-125.6813, 41.5800] |
| $x_{BLA}\dot{x}_{CeA}$ | -0.2961 | [-27.5750, 28.3352] | 2.5395 | [-25.1237, 36.5010] |
| $\dot{x}_{BLA}x_{CeA}$ | -5.3197 | [-29.2786, 13.9949] | -6.7713 | [-26.9807, 11.4020] |
| $\dot{x}_{BLA}\dot{x}_{CeA}$ | 0.1295 | [-1.6524, 1.8954] | -0.4911 | [-2.7676, 1.7339] |

* Indicates statistical significance (95% CI does not overlap with zero).

Table D10: Coefficients of Population-level Regression Model (CeA-DMS as target pathway)

| Variables | Control Mean Coefficients | 95% CI | Stress Mean Coefficients | 95% CI |
|---|---|---|---|---|
| $x_{CeA}$ | -92.0185 | [-198.2043, -13.2261]* | -18.9510 | [-82.9084, 20.4310] |
| $\dot{x}_{CeA}$ | -30.4950 | [-35.7646, -24.4500]* | -26.9429 | [-37.0648, -16.6963]* |
| $x^2_{CeA}$ | 38.2353 | [-60.4707, 121.2375] | -13.6061 | [-57.1711, 41.8892] |
| $x_{CeA}\dot{x}_{CeA}$ | 3.9223 | [-23.3504, 26.5724] | -1.2625 | [-16.5882, 13.6153] |
| $\dot{x}^2_{CeA}$ | -0.0640 | [-1.3730, 1.2717] | -0.3545 | [-2.1591, 1.4021] |
| $x_{BLA}$ | 15.0593 | [-28.7133, 68.5997] | 7.7932 | [-31.3337, 43.1903] |
| $\dot{x}_{BLA}$ | 0.1626 | [-3.1432, 3.2865] | -0.3904 | [-14.0078, 10.6121] |
| $x_{CeA}x_{BLA}$ | 8.8987 | [-64.5687, 104.4596] | 13.3426 | [-38.6467, 84.2520] |
| $x_{CeA}\dot{x}_{BLA}$ | -1.1719 | [-21.3051, 20.1282] | 2.4396 | [-16.0964, 27.5882] |
| $\dot{x}_{CeA}x_{BLA}$ | 0.0219 | [-18.1112, 21.1292] | -0.6001 | [-22.7219, 22.4070] |
| $\dot{x}_{CeA}\dot{x}_{BLA}$ | -0.0157 | [-1.5400, 1.4706] | -0.3922 | [-2.5207, 1.5418] |

* Indicates statistical significance (95% CI does not overlap with zero).

##### D.2.2 Dynamical Perspective: Tables D13-D19

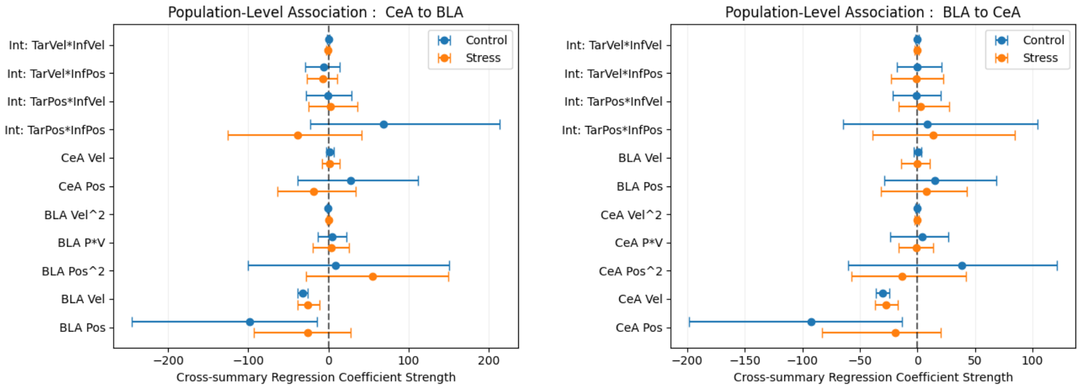


(a) Regression configuration: CeA-DMS to BLA-DMS

(b) Regression configuration: BLA-DMS to CeA-DMS

Figure D6: Forest plots show cross-summary regression coefficients from the population-level dynamical model.

Table D11: Statistical Robustness of Integrated KL Divergence Following Pressing of Each Session

| Pathway | Group | Session | Mean Cumulative $D_{KL}$ | 95% CI |
|---|---|---|---|---|
| BLA-DMS | Control | FR1 | 16816.1492 | [2372.1439,82475.2567] |
| BLA-DMS | Control | RR2 | 19767.3561 | [5853.7709,73573.8874] |
| BLA-DMS | Control | RR5 | 22076.9192 | [6430.3595,58165.5443] |
| BLA-DMS | Control | RR10 | 8477.2925 | [3717.3232,18611.9694] |
| BLA-DMS | Stress | FR1 | 43024.1266 | [21415.111,104372.608] |
| BLA-DMS | Stress | RR2 | 20961.4737 | [10326.7752,58847.6864] |
| BLA-DMS | Stress | RR5 | 16932.2606 | [5329.422,55730.6548] |
| BLA-DMS | Stress | RR10 | 55011.6805 | [16479.7293,161800.8385] |
| CeA-DMS | Control | FR1 | 2699.0721 | [958.0845,6744.1825] |
| CeA-DMS | Control | RR2 | 16873.9109 | [7784.1484,40107.5754] |
| CeA-DMS | Control | RR5 | 32465.1389 | [9600.4193,67246.1506] |
| CeA-DMS | Control | RR10 | 10021.7906 | [2190.8027,30358.8264] |
| CeA-DMS | Stress | FR1 | 5651.2204 | [2188.9462,14328.1062] |
| CeA-DMS | Stress | RR2 | 13516.2779 | [5099.8194,29266.544] |
| CeA-DMS | Stress | RR5 | 19631.6303 | [8593.419,33346.937] |
| CeA-DMS | Stress | RR10 | 21425.9159 | [5357.5366,70988.8337] |

#### D.3 Sensitivity Analysis of KDE Bandwidth Choice

To evaluate whether the observed KL-divergence patterns were sensitive to kernel-density estimation (KDE) parameters, we performed a bandwidth sensitivity analysis. The default bandwidth was selected using Scott's Rule, a data-driven estimator based on sample size and variance.

We then perturbed the optimal bandwidth($0.9 \times h_{opt}$ [narrow] and $1.1 \times h_{opt}$ [wide]) and recomputed the cumulative KL-divergence trajectories for the BLA-DMS pathway during footshock. As shown in Figure D9, changing the bandwidth affected the absolute magnitude of the cumulative KL-divergence values, as expected from differences in smoothing. However, the relative separation between the Control and Stress groups remained qualitatively unchanged across bandwidth settings. Corresponding 95% confidence intervals are reported in Table D20.

Table D12: Statistical Robustness of Integrated KL Divergence Following Reward of Each Session

| Pathway | Group | Session | Mean Cumulative $D_{KL}$ | 95% CI |
|---|---|---|---|---|
| BLA-DMS | Control | FR1 | 20779.2454 | [7996.1546,51824.8836] |
| BLA-DMS | Control | RR2 | 40175.1672 | [10775.304,79629.8986] |
| BLA-DMS | Control | RR5 | 23159.2532 | [3772.7712,78079.9234] |
| BLA-DMS | Control | RR10 | 34616.4245 | [17385.6191,54192.0772] |
| BLA-DMS | Stress | FR1 | 61375.3241 | [37042.1089,122405.9986] |
| BLA-DMS | Stress | RR2 | 24642.2449 | [13428.5497,45501.857] |
| BLA-DMS | Stress | RR5 | 18014.9215 | [3266.036,49379.3554] |
| BLA-DMS | Stress | RR10 | 37260.1647 | [8272.4422,103366.8172] |
| CeA-DMS | Control | FR1 | 19560.1202 | [3830.0245,71783.6601] |
| CeA-DMS | Control | RR2 | 4772.6126 | [2161.7166,14189.1465] |
| CeA-DMS | Control | RR5 | 18483.5742 | [5162.3285,45015.4785] |
| CeA-DMS | Control | RR10 | 31912.9089 | [10617.6616,78462.0686] |
| CeA-DMS | Stress | FR1 | 7464.6371 | [3850.7582,13130.0388] |
| CeA-DMS | Stress | RR2 | 27528.9806 | [11352.1426,64529.4044] |
| CeA-DMS | Stress | RR5 | 58867.3343 | [24396.8574,107702.6232] |
| CeA-DMS | Stress | RR10 | 52516.3992 | [27365.565,124927.3165] |

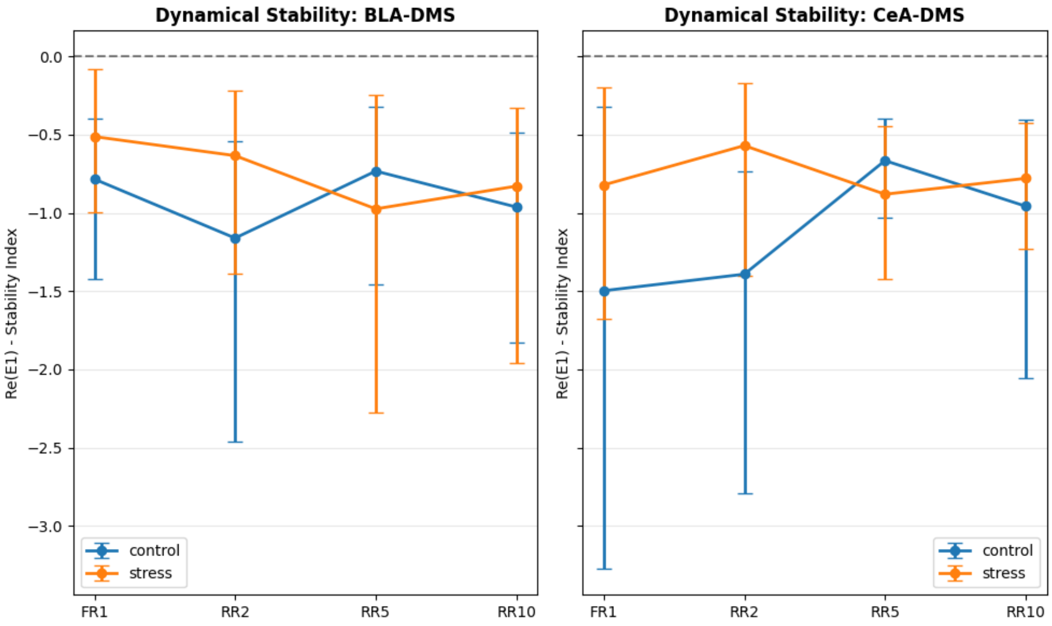


Figure D7: Statistical Distribution of Dominant Jacobian Eigenvalues After Bout-Initiating Pressing. Forest plot displaying the mean and 95% confidence intervals of the real part of the dominant Jacobian eigenvalue ($Re(E_1)$) obtained from the reconstructed univariate dynamical model for the BLA-DMS and CeA-DMS pathways during lever pressing. The distributions were estimated from $n = 1000$ bootstrap iterations. Across sessions (FR1-RR10), the estimated eigenvalue distributions remained predominantly negative, consistent with local stability of the reconstructed trajectories around the empirical equilibrium. The substantial overlap between control (blue) and stress (orange) groups indicates broadly similar stability estimates within the fitted phenomenological model.

Table D13: Regression Coefficients of the Reconstructed Dynamical Models Across Learning Stages (Operant Lever Pressing)

| Pathway | Session | Term | Control Mean | 95% CI | Stress Mean | 95% CI |
|---|---|---|---|---|---|---|
| BLA-DMS | FR1 | Intercept | 1.2712 | [ -2.3786, 7.5848] | -4.5539 | [ -9.7132, -0.8905] * |
| | | $x_{BLA}$ | -16.7932 | [ -26.2511, -8.7176] * | -6.8968 | [ -14.5424, -0.8534] * |
| | | $\dot{x}_{BLA}$ | -23.9413 | [ -27.3359, -20.6387] * | -26.7075 | [ -30.3255, -22.7223] * |
| | | $x^2_{BLA}$ | -6.3576 | [ -37.5373, 5.7211] | 7.9442 | [ 1.5195, 15.6654] * |
| | | $x_{BLA}\dot{x}_{BLA}$ | -8.2205 | [ -17.2998, 2.1525] | 4.4405 | [ -1.7644, 11.9634] |
| | | $\dot{x}^2_{BLA}$ | -0.7182 | [ -3.1023, 2.1449] | -1.5148 | [ -4.4279, 0.9394] |
| CeA-DMS | FR1 | Intercept | -7.2434 | [ -23.4901, 1.2669] | 0.1446 | [ -3.9513, 5.3153] |
| | | $x_{CeA}$ | -30.0133 | [ -57.4265, -7.9078] * | -22.7966 | [ -47.7347, -6.1556] * |
| | | $\dot{x}_{CeA}$ | -25.8503 | [ -29.5162, -20.6607] * | -28.5185 | [ -31.3561, -25.1919] * |
| | | $x^2_{CeA}$ | 15.9658 | [ 0.1044, 43.4168] * | 0.1724 | [ -11.7799, 8.7144] |
| | | $x_{CeA}\dot{x}_{CeA}$ | 10.5297 | [ -2.1263, 23.3268] | 0.4284 | [ -11.5630, 13.5678] |
| | | $\dot{x}^2_{CeA}$ | -0.4998 | [ -2.0890, 2.8188] | -0.138 | [ -2.3139, 1.4985] |
| BLA-DMS | RR2 | Intercept | 5.4888 | [ -2.4378, 18.3767] | -6.4581 | [ -13.3615, -2.5967] * |
| | | $x_{BLA}$ | -23.3321 | [ -48.2440, -8.2264] * | -8.4262 | [ -22.9443, -1.0713] * |
| | | $\dot{x}_{BLA}$ | -26.9326 | [ -32.8096, -21.6539] * | -24.1228 | [ -28.5033, -19.4136] * |
| | | $x^2_{BLA}$ | -10.3613 | [ -32.5278, 5.2695] | 9.1015 | [ 3.8026, 18.5210] * |
| | | $x_{BLA}\dot{x}_{BLA}$ | 1.9898 | [ -10.1158, 16.2188] | 7.9266 | [ -1.3280, 14.3093] |
| | | $\dot{x}^2_{BLA}$ | -0.5516 | [ -3.2334, 1.8200] | 0.9359 | [ -1.5046, 4.3840] |
| CeA-DMS | RR2 | Intercept | -7.7089 | [ -15.0675, -2.3021] * | 0.7761 | [ -2.1409, 4.9105] |
| | | $x_{CeA}$ | -30.0102 | [ -66.0351, -12.3878] * | -14.0464 | [ -36.9369, -3.8021] * |
| | | $\dot{x}_{CeA}$ | -26.9752 | [ -31.9238, -21.4754] * | -27.607 | [ -31.7346, -22.7773] * |
| | | $x^2_{CeA}$ | 12.8896 | [ 3.2139, 26.4671] * | -2.2253 | [ -11.1252, 4.8487] |
| | | $x_{CeA}\dot{x}_{CeA}$ | 5.1413 | [ -12.2320, 17.2621] | 1.562 | [ -8.2181, 11.1112] |
| | | $\dot{x}^2_{CeA}$ | -1.1222 | [ -2.8875, 0.9589] | -0.2806 | [ -3.1200, 2.4826] |
| BLA-DMS | RR5 | Intercept | 1.5048 | [ -1.9884, 4.8024] | -7.9965 | [ -17.5491, -1.8634] * |
| | | $x_{BLA}$ | -17.8966 | [ -31.7472, -8.1769] * | -15.9429 | [ -37.5970, -1.6101] * |
| | | $\dot{x}_{BLA}$ | -26.6403 | [ -29.4009, -23.3475] * | -26.3634 | [ -31.4469, -21.5874] * |
| | | $x^2_{BLA}$ | -3.1367 | [ -17.3952, 4.1346] | 12.9722 | [ 3.9089, 27.7909] * |
| | | $x_{BLA}\dot{x}_{BLA}$ | -5.2615 | [ -18.1205, 4.8795] | 5.6949 | [ -7.6435, 15.2372] |
| | | $\dot{x}^2_{BLA}$ | -1.2047 | [ -3.6665, 1.1672] | -0.3694 | [ -3.0950, 1.9622] |
| CeA-DMS | RR5 | Intercept | -3.7817 | [ -7.2668, -0.5990] * | 2.3344 | [ -0.4139, 5.6975] |
| | | $x_{CeA}$ | -16.627 | [ -25.5381, -8.8942] * | -21.4448 | [ -32.8877, -10.1296] * |
| | | $\dot{x}_{CeA}$ | -28.4015 | [ -32.1625, -24.5586] * | -26.6243 | [ -29.4545, -22.7915] * |
| | | $x^2_{CeA}$ | 4.7267 | [ 0.0896, 9.9620] * | -5.1593 | [ -15.6078, 0.8026] |
| | | $x_{CeA}\dot{x}_{CeA}$ | 1.9315 | [ -7.7722, 13.8645] | -2.6786 | [ -13.5873, 10.1936] |
| | | $\dot{x}^2_{CeA}$ | 0.7566 | [ -1.8281, 3.3057] | 0.7404 | [ -1.4535, 3.6630] |
| BLA-DMS | RR10 | Intercept | 0.9222 | [ -4.3107, 5.7367] | -6.8144 | [ -14.5928, -2.5355] * |
| | | $x_{BLA}$ | -23.5777 | [ -45.2631, -10.0863] * | -15.5389 | [ -34.6631, -4.3950] * |
| | | $\dot{x}_{BLA}$ | -26.4676 | [ -29.1329, -23.0011] * | -25.5338 | [ -29.9856, -21.2664] * |
| | | $x^2_{BLA}$ | -2.2196 | [ -20.4440, 9.8049] | 9.9562 | [ 3.2377, 23.0684] * |
| | | $x_{BLA}\dot{x}_{BLA}$ | -0.0435 | [ -13.7765, 11.8675] | 6.5996 | [ -8.2139, 15.9745] |
| | | $\dot{x}^2_{BLA}$ | -0.5917 | [ -2.2264, 1.4093] | 0.4519 | [ -2.2080, 2.8794] |
| CeA-DMS | RR10 | Intercept | -5.2061 | [ -12.9868, -0.0624] * | 3.3342 | [ -0.6833, 7.8437] |
| | | $x_{CeA}$ | -25.3961 | [ -57.9887, -10.5250] * | -17.6322 | [ -25.7508, -9.4809] * |
| | | $\dot{x}_{CeA}$ | -29.1342 | [ -32.3675, -25.3054] * | -27.5671 | [ -30.0983, -24.1701] * |
| | | $x^2_{CeA}$ | 7.7333 | [ -3.4088, 28.9279] | -7.205 | [ -22.5222, 5.5463] |
| | | $x_{CeA}\dot{x}_{CeA}$ | -0.3174 | [ -12.1224, 10.8408] | -1.0816 | [ -11.1387, 9.6965] |
| | | $\dot{x}^2_{CeA}$ | 0.1427 | [ -2.8908, 2.1151] | 0.1553 | [ -2.1756, 2.8886] |

* Indicates statistical significance (95% CI does not overlap with zero).

These analyses indicate that the reported KL-divergence differences are not driven by a specific bandwidth choice and are robust within a reasonable range of KDE smoothing parameters. Consistent with the

Table D14: Regression Coefficients of the Reconstructed Dynamical Models Across Learning Stages (Contingent Reward Delivery)

| Pathway | Session | Term | Control Mean | 95% CI | Stress Mean | 95% CI |
|---|---|---|---|---|---|---|
| BLA-DMS | FR1 | Intercept | -0.3843 | [ -5.7995, 9.1509] | -9.677 | [ -21.1479, 0.2909] |
| | | $x_{BLA}$ | -16.9427 | [ -41.5624, -6.1998] * | -10.3041 | [ -25.5124, -0.4477] * |
| | | $\dot{x}_{BLA}$ | -27.6025 | [ -31.8651, -22.9625] * | -23.7977 | [ -32.6377, -16.2796] * |
| | | $x^2_{BLA}$ | 5.8881 | [ -13.3293, 23.5769] | 10.6416 | [ 1.7467, 23.9370] * |
| | | $x_{BLA}\dot{x}_{BLA}$ | 3.3408 | [ -21.2317, 17.0397] | 6.2346 | [ -7.9675, 16.7377] |
| | | $\dot{x}^2_{BLA}$ | 0.5563 | [ -1.8016, 2.7815] | 0.4149 | [ -2.5510, 3.8662] |
| CeA-DMS | FR1 | Intercept | -3.4226 | [ -10.7325, 4.9361] | 0.4525 | [ -2.4295, 4.2184] |
| | | $x_{CeA}$ | -27.234 | [ -55.9397, -10.1359] * | -22.8367 | [ -66.3475, -8.5566] * |
| | | $\dot{x}_{CeA}$ | -28.6038 | [ -31.5329, -24.9884] * | -28.3777 | [ -30.9853, -25.6004] * |
| | | $x^2_{CeA}$ | 6.561 | [ -5.8795, 17.3110] | -6.9012 | [ -24.5523, 1.6153] |
| | | $x_{CeA}\dot{x}_{CeA}$ | 2.1403 | [ -13.1078, 21.5605] | -4.1301 | [ -15.9557, 7.0743] |
| | | $\dot{x}^2_{CeA}$ | -0.0967 | [ -2.3966, 2.8164] | 0.3541 | [ -1.3242, 2.3404] |
| BLA-DMS | RR2 | Intercept | 2.5964 | [ -6.8191, 21.6276] | -8.8028 | [ -19.5706, -1.4069] * |
| | | $x_{BLA}$ | -23.1327 | [ -52.8191, -9.2973] * | -11.8323 | [ -27.3396, -1.5943] * |
| | | $\dot{x}_{BLA}$ | -26.7117 | [ -30.2458, -23.0865] * | -26.0786 | [ -34.1990, -19.5727] * |
| | | $x^2_{BLA}$ | 7.7317 | [ -24.4306, 33.4196] | 11.2511 | [ 1.7957, 24.5899] * |
| | | $x_{BLA}\dot{x}_{BLA}$ | 1.8771 | [ -12.2609, 12.4772] | 5.1322 | [ -10.8490, 16.1347] |
| | | $\dot{x}^2_{BLA}$ | -0.7187 | [ -2.3260, 1.5908] | 0.7929 | [ -1.6623, 3.4407] |
| CeA-DMS | RR2 | Intercept | -0.5242 | [ -5.7302, 6.1979] | 2.3015 | [ -2.8163, 10.5868] |
| | | $x_{CeA}$ | -24.3698 | [ -54.7970, -7.0123] * | -22.2851 | [ -51.9297, -9.3713] * |
| | | $\dot{x}_{CeA}$ | -27.1574 | [ -29.8181, -24.1091] * | -27.8106 | [ -30.4389, -24.2136] * |
| | | $x^2_{CeA}$ | -3.0785 | [ -32.3723, 11.0217] | -3.2122 | [ -17.2392, 5.4550] |
| | | $x_{CeA}\dot{x}_{CeA}$ | -0.2573 | [ -14.9042, 11.2403] | -2.4768 | [ -13.6383, 9.4297] |
| | | $\dot{x}^2_{CeA}$ | -0.9931 | [ -2.2543, 0.9182] | 0.1654 | [ -1.5824, 2.9033] |
| BLA-DMS | RR5 | Intercept | 1.4335 | [ -5.4506, 10.7005] | -10.1086 | [ -21.3729, -2.4910] * |
| | | $x_{BLA}$ | -24.9842 | [ -54.5820, -9.3385] * | -18.0072 | [ -41.2761, -4.5856] * |
| | | $\dot{x}_{BLA}$ | -27.7398 | [ -31.6671, -24.1157] * | -25.8395 | [ -33.1486, -18.2584] * |
| | | $x^2_{BLA}$ | 2.3427 | [ -16.5419, 20.7000] | 14.4788 | [ 0.9297, 31.7778] * |
| | | $x_{BLA}\dot{x}_{BLA}$ | 1.361 | [ -10.4464, 14.9628] | 3.5286 | [ -13.6324, 18.4408] |
| | | $\dot{x}^2_{BLA}$ | -0.1914 | [ -2.3885, 2.4886] | 0.565 | [ -1.8618, 4.2813] |
| CeA-DMS | RR5 | Intercept | -0.2492 | [ -3.6396, 4.2873] | 4.508 | [ 0.5930, 12.0949] * |
| | | $x_{CeA}$ | -15.0153 | [ -31.8495, -2.1089] * | -16.7161 | [ -36.6164, -3.2420] * |
| | | $\dot{x}_{CeA}$ | -29.014 | [ -32.3352, -25.7642] * | -28.636 | [ -33.6542, -24.7766] * |
| | | $x^2_{CeA}$ | -1.328 | [ -15.7100, 7.9578] | -10.0115 | [ -25.3317, -2.4162] * |
| | | $x_{CeA}\dot{x}_{CeA}$ | -2.6551 | [ -13.2258, 7.9731] | -2.1497 | [ -13.3518, 12.7232] |
| | | $\dot{x}^2_{CeA}$ | 0.1937 | [ -2.2669, 1.9984] | 0.2958 | [ -1.6203, 2.1437] |
| BLA-DMS | RR10 | Intercept | 0.9619 | [ -7.9330, 15.6095] | -8.8394 | [ -22.0320, -2.1155] * |
| | | $x_{BLA}$ | -30.3819 | [ -71.3813, -10.2099] * | -16.796 | [ -45.6861, -5.3380] * |
| | | $\dot{x}_{BLA}$ | -27.5751 | [ -31.1033, -24.2332] * | -24.3471 | [ -30.4648, -19.7917] * |
| | | $x^2_{BLA}$ | 6.7833 | [ -16.8917, 27.7254] | 12.8132 | [ 2.2670, 31.2512] * |
| | | $x_{BLA}\dot{x}_{BLA}$ | 3.6874 | [ -10.4825, 13.7598] | 7.2731 | [ -6.5339, 15.0526] |
| | | $\dot{x}^2_{BLA}$ | -0.3529 | [ -2.7521, 1.3318] | 0.0325 | [ -2.3372, 2.6915] |
| CeA-DMS | RR10 | Intercept | -0.0773 | [ -6.9297, 7.7160] | 12.4221 | [ 4.5612, 23.3921] * |
| | | $x_{CeA}$ | -24.9047 | [ -58.6285, -5.1298] * | -24.0998 | [ -50.3895, -6.8594] * |
| | | $\dot{x}_{CeA}$ | -29.056 | [ -31.9831, -24.7099] * | -26.1875 | [ -30.4673, -22.4364] * |
| | | $x^2_{CeA}$ | -6.3689 | [ -48.8102, 14.4061] | -19.8416 | [ -38.4165, -3.6166] * |
| | | $x_{CeA}\dot{x}_{CeA}$ | 1.3837 | [ -12.1729, 15.8929] | -3.8858 | [ -14.4820, 6.3183] |
| | | $\dot{x}^2_{CeA}$ | -0.2068 | [ -1.8317, 1.1353] | -0.5434 | [ -2.0405, 1.6323] |

* Indicates statistical significance (95% CI does not overlap with zero).

framework of McNealis and Léger (2026), kernel smoothing was applied across all bandwidth conditions to provide comparable uncertainty estimates for the non-Gaussian calcium-signal distributions.

Table D15: Statistical Robustness of Largest Real Part of Eigenvalue of Each Session

| Pathway | Group | Session | Mean $ReE_1$ | 95% CI |
|---|---|---|---|---|
| BLA-DMS | Control | FR1 | -0.7845 | [ -1.422, -0.399] |
| BLA-DMS | Control | RR2 | -1.1610 | [ -2.462, -0.540] |
| BLA-DMS | Control | RR5 | -0.7331 | [ -1.454, -0.320] |
| BLA-DMS | control | RR10 | -0.9628 | [ -1.829, -0.485] |
| BLA-DMS | Stress | FR1 | -0.5128 | [ -0.998, -0.083] |
| BLA-DMS | Stress | RR2 | -0.6345 | [ -1.390, -0.219] |
| BLA-DMS | Stress | RR5 | -0.9744 | [ -2.277, -0.250] |
| BLA-DMS | Stress | RR10 | -0.8291 | [ -1.959, -0.331] |
| CeA-DMS | Control | FR1 | -1.4973 | [ -3.276, -0.319] |
| CeA-DMS | Control | RR2 | -1.3917 | [ -2.795, -0.734] |
| CeA-DMS | Control | RR5 | -0.6661 | [ -1.033, -0.399] |
| CeA-DMS | Control | RR10 | -0.9573 | [ -2.059, -0.408] |
| CeA-DMS | Stress | FR1 | -0.8216 | [ -1.680, -0.197] |
| CeA-DMS | Stress | RR2 | -0.5702 | [ -1.400, -0.170] |
| CeA-DMS | Stress | RR5 | -0.8809 | [ -1.423, -0.446] |
| CeA-DMS | Stress | RR10 | -0.7794 | [ -1.232, -0.428] |

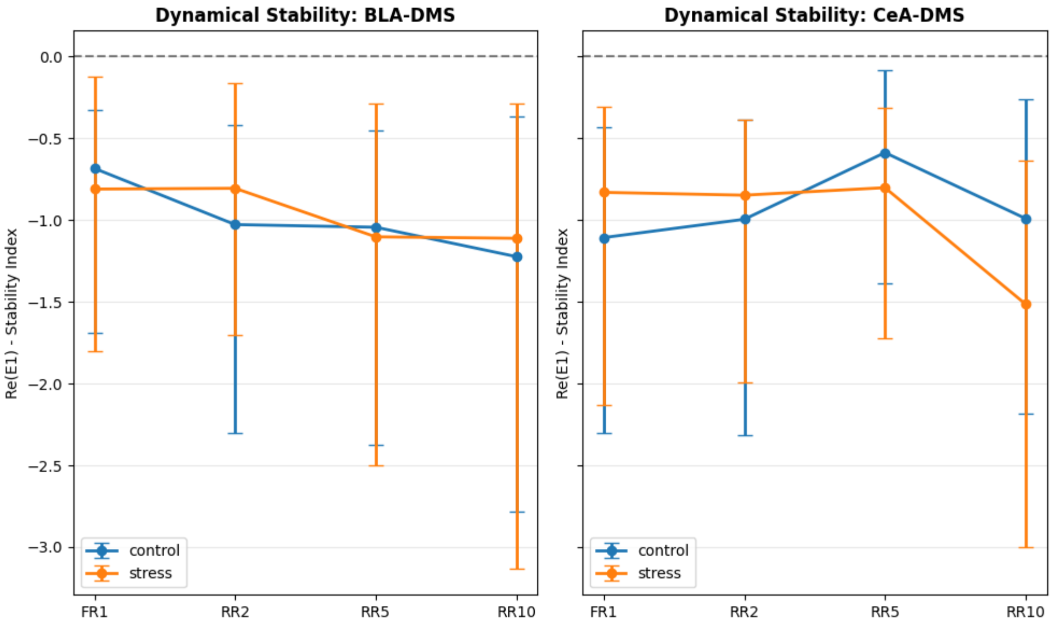


Figure D8: Statistical Distribution of Dominant Jacobian Eigenvalues After Reward. Forest plot displaying the mean and 95% confidence intervals of the real part of the dominant Jacobian eigenvalue ($Re(E_1)$) obtained from the reconstructed univariate dynamical model for the BLA-DMS and CeA-DMS pathways during reinforcement (reward-onset) phase. The distributions were estimated from $n = 1000$ bootstrap iterations. Across sessions (FR1-RR10), the estimated eigenvalue distributions remained predominantly negative, consistent with local stability of the reconstructed trajectories around the empirical equilibrium. The substantial overlap between control (blue) and stress (orange) groups indicates broadly similar stability estimates within the fitted phenomenological model.

Table D16: Statistical Robustness of Largest Real Part of Eigenvalue of Each Session

| Pathway | Group | Session | Mean $ReE_1$ | 95% CI |
|---|---|---|---|---|
| BLA-DMS | Control | FR1 | -0.6849 | [ -1.691, -0.329] |
| BLA-DMS | Control | RR2 | -1.0283 | [ -2.300, -0.418] |
| BLA-DMS | Control | RR5 | -1.0446 | [ -2.377, -0.454] |
| BLA-DMS | Control | RR10 | -1.2243 | [ -2.780, -0.367] |
| BLA-DMS | Stress | FR1 | -0.8113 | [ -1.799, -0.122] |
| BLA-DMS | Stress | RR2 | -0.8067 | [ -1.706, -0.163] |
| BLA-DMS | Stress | RR5 | -1.1033 | [ -2.501, -0.290] |
| BLA-DMS | Stress | RR10 | -1.1122 | [ -3.135, -0.288] |
| CeA-DMS | Control | FR1 | -1.1080 | [ -2.302, -0.435] |
| CeA-DMS | Control | RR2 | -0.9960 | [ -2.313, -0.385] |
| CeA-DMS | Control | RR5 | -0.5885 | [ -1.387, -0.085] |
| CeA-DMS | Control | RR10 | -0.9915 | [ -2.181, -0.259] |
| CeA-DMS | Stress | FR1 | -0.8315 | [ -2.134, -0.310] |
| CeA-DMS | Stress | RR2 | -0.8487 | [ -1.992, -0.385] |
| CeA-DMS | Stress | RR5 | -0.8029 | [ -1.725, -0.316] |
| CeA-DMS | Stress | RR10 | -1.5152 | [ -2.997, -0.639] |

Table D17: Mean and Standard Deviation Before and After Perturbation for Each Group, Pathway and Sessions

| | | | Bout-Initiating Presses | | | | Reward | | | |
|---|---|---|---|---|---|---|---|---|---|---|
| Group | Pathway | Session | Before Perturbation | | After Perturbation | | Before Perturbation | | After Perturbation | |
| | | | Mean | Standard Deviation | Mean | Standard Deviation | Mean | Standard Deviation | Mean | Standard Deviation |
| Control | BLA-DMS | FR-1 | -0.1646 | 0.273 | 0.0276 | 0.3599 | -0.1148 | 0.3286 | 0.0013 | 0.5163 |
| | | RR-2 | 0.0396 | 0.3091 | 0.1811 | 0.5335 | 0.0612 | 0.317 | 0.0177 | 0.5781 |
| | | RR-5 | -0.0949 | 0.2009 | 0.0239 | 0.3012 | -0.1299 | 0.2749 | -0.0006 | 0.4203 |
| | | RR-10 | -0.0327 | 0.2788 | -0.0599 | 0.3453 | -0.1798 | 0.259 | 0.0033 | 0.43 |
| | CeA-DMS | FR-1 | -0.1787 | 0.4134 | -0.2178 | 0.4639 | -0.1696 | 0.2589 | -0.1512 | 0.4079 |
| | | RR-2 | -0.0686 | 0.2367 | -0.2642 | 0.3378 | -0.1929 | 0.3713 | -0.0913 | 0.4067 |
| | | RR-5 | -0.0531 | 0.2112 | -0.1903 | 0.3328 | -0.0859 | 0.2628 | 0.0034 | 0.3829 |
| | | RR-10 | -0.0401 | 0.2659 | -0.1484 | 0.3567 | -0.174 | 0.2733 | -0.0149 | 0.4906 |
| Stress | BLA-DMS | FR-1 | -0.0477 | 0.2545 | -0.4853 | 0.2363 | -0.1056 | 0.2504 | -0.5228 | 0.39 |
| | | RR-2 | -0.0977 | 0.3392 | -0.5101 | 0.3448 | -0.1962 | 0.346 | -0.4532 | 0.3779 |
| | | RR-5 | -0.1946 | 0.2426 | -0.4142 | 0.4051 | -0.2327 | 0.2769 | -0.3854 | 0.4245 |
| | | RR-10 | -0.0601 | 0.2117 | -0.3598 | 0.4095 | -0.1126 | 0.2546 | -0.3761 | 0.4283 |
| | CeA-DMS | FR-1 | -0.079 | 0.2885 | -0.0084 | 0.2949 | -0.0623 | 0.2761 | 0.0746 | 0.2255 |
| | | RR-2 | -0.1014 | 0.2535 | 0.0244 | 0.3791 | -0.2146 | 0.2228 | 0.1005 | 0.3762 |
| | | RR-5 | -0.1004 | 0.2245 | 0.1262 | 0.2934 | -0.1458 | 0.2221 | 0.2809 | 0.3931 |
| | | RR-10 | -0.0586 | 0.2616 | 0.1567 | 0.3645 | -0.0959 | 0.2289 | 0.3647 | 0.2996 |

This consistent group-level difference across the tested range demonstrates that our findings are robust to the choice of smoothing parameters. This approach aligns with recent methodological advancements, such as the smoothed pseudo-population bootstrap, which advocates for smoothing to enhance the efficiency and reliability of uncertainty quantification in complex, non-smooth data distributions like neuronal calcium transients.

Note: To optimize computational efficiency during multi-parameter sensitivity testing, 200 bootstrap iterations were performed for each factor. Results confirm that the directional impact of stress on informational hysteresis is independent of bandwidth selection.

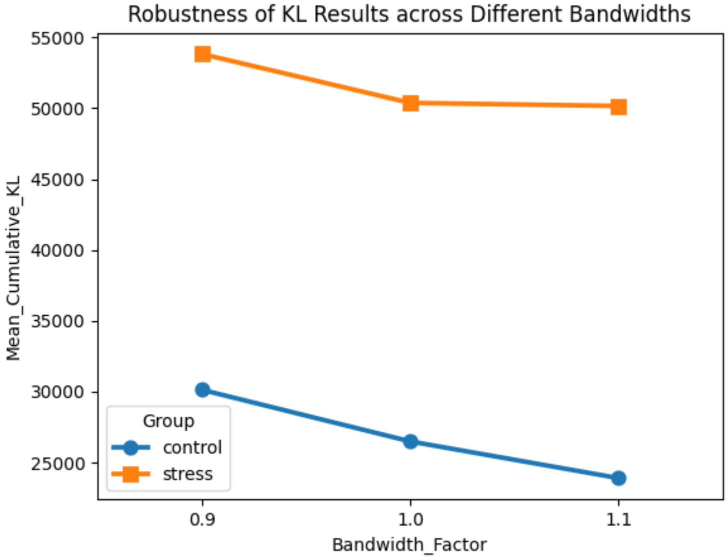


Figure D9: Sensitivity analysis of KDE bandwidth selection for KL divergence estimation. Mean cumulative KL divergence for the BLA-DMS pathway (footshock) under three different bandwidth scaling factors (0.9, 1.0, and 1.1 of the data-driven Scott's Rule). While the absolute magnitude of the KL metric decreases slightly as the smoothing bandwidth increases (from 0.9 to 1.1), the directional shift—where the Stress group exhibits significantly higher informational hysteresis than the Control group—remains invariant across all tested parameters.

### References

Table D18: Population-level Dynamical Model Coefficients of Bout-Initiating Pressing of Both Target Pathways

| | | BLA-DMS to CeA-DMS | | | CeA-DMS to BLA-DMS | | |
|---|---|---|---|---|---|---|---|
| Group | Session | Term | Mean | 95% CI | Term | Mean | 95% CI |
| Control | FR1 | $x_{CeA}$ | -54.3788 | [-106.3857,-12.9669]* | $x_{BLA}$ | -29.1248 | [-53.539,-9.1401]* |
| | | $\dot{x}_{CeA}$ | -25.3815 | [-32.2152,-16.3534]* | $\dot{x}_{BLA}$ | -24.5639 | [-32.7806,-13.1737]* |
| | | $x^2_{CeA}$ | 19.4912 | [-21.6885,66.2179] | $x^2_{BLA}$ | -41.574 | [-128.3198,12.9708] |
| | | $x_{CeA}\dot{x}_{CeA}$ | 15.2289 | [-11.9704,43.0468] | $x_{BLA}\dot{x}_{BLA}$ | -14.0613 | [-29.7352,8.6172] |
| | | $\dot{x}^2_{CeA}$ | -0.4411 | [-2.0755,3.0235] | $\dot{x}^2_{BLA}$ | -0.9757 | [-3.7375,1.5205] |
| | | $x_{BLA}$ | 3.8417 | [-38.1831,57.8168] | $x_{CeA}$ | -9.351 | [-53.9587,14.6855] |
| | | $\dot{x}_{BLA}$ | -0.0039 | [-11.8715,9.0577] | $\dot{x}_{CeA}$ | 0.0525 | [-1.933,2.9489] |
| | | $x_{CeA}x_{BLA}$ | 15.4236 | [-24.5184,78.5969] | $x_{CeA}x_{BLA}$ | 10.2463 | [-35.5353,58.673] |
| | | $x_{BLA}\dot{x}_{CeA}$ | -3.6904 | [-38.085,32.2286] | $x_{CeA}\dot{x}_{BLA}$ | -4.9837 | [-23.7554,9.7735] |
| | | $\dot{x}_{BLA}x_{CeA}$ | 1.5477 | [-16.5416,24.7061] | $\dot{x}_{CeA}x_{BLA}$ | 2.0336 | [-22.6211,42.2287] |
| | | $\dot{x}_{BLA}x_{BLA}$ | -0.6331 | [-4.1389,2.9228] | $\dot{x}_{CeA}x_{CeA}$ | -0.8334 | [-3.4726,1.8395] |
| Control | RR2 | $x_{CeA}$ | -53.2852 | [-133.265,-9.6493]* | $x_{BLA}$ | -36.2488 | [-102.1649,12.5141] |
| | | $\dot{x}_{CeA}$ | -29.8049 | [-41.0995,-19.708]* | $\dot{x}_{BLA}$ | -28.9246 | [-38.0586,-20.6601]* |
| | | $x^2_{CeA}$ | 31.5156 | [-8.6962,68.1336] | $x^2_{BLA}$ | -29.3516 | [-98.0962,25.9677] |
| | | $x_{CeA}\dot{x}_{CeA}$ | -0.7185 | [-35.6572,30.7766] | $x_{BLA}\dot{x}_{BLA}$ | 3.2338 | [-19.145,25.1164] |
| | | $\dot{x}^2_{CeA}$ | -1.3184 | [-3.1241,0.7439] | $\dot{x}^2_{BLA}$ | -0.4395 | [-3.5096,2.0831] |
| | | $x_{BLA}$ | -6.6579 | [-38.5672,20.4471] | $x_{CeA}$ | -22.4161 | [-65.903,10.2782] |
| | | $\dot{x}_{BLA}$ | 2.0072 | [-5.6841,12.4024] | $\dot{x}_{CeA}$ | -1.6942 | [-10.4005,6.2309] |
| | | $x_{CeA}x_{BLA}$ | -17.8755 | [-49.67,10.9387] | $x_{CeA}x_{BLA}$ | 22.9977 | [-8.944,65.9113] |
| | | $x_{BLA}\dot{x}_{CeA}$ | 1.8726 | [-28.63,36.7707] | $x_{CeA}\dot{x}_{BLA}$ | 5.614 | [-25.8833,39.4991] |
| | | $\dot{x}_{BLA}x_{CeA}$ | 1.1153 | [-21.0205,22.4045] | $\dot{x}_{CeA}x_{BLA}$ | -2.7249 | [-29.9563,21.5702] |
| | | $\dot{x}_{BLA}x_{BLA}$ | 0.4647 | [-2.1836,3.1435] | $\dot{x}_{CeA}x_{CeA}$ | 0.0905 | [-3.5099,3.911] |
| Control | RR5 | $x_{CeA}$ | -38.7982 | [-71.2968,-17.0835]* | $x_{BLA}$ | -31.7505 | [-68.7426,-8.3237]* |
| | | $\dot{x}_{CeA}$ | -29.6613 | [-34.7489,-24.1258]* | $\dot{x}_{BLA}$ | -27.753 | [-34.0114,-21.8248]* |
| | | $x^2_{CeA}$ | 11.7315 | [-9.786,36.7703] | $x^2_{BLA}$ | -25.0792 | [-85.1128,12.1288] |
| | | $x_{CeA}\dot{x}_{CeA}$ | 1.4064 | [-21.9198,29.644] | $x_{BLA}\dot{x}_{BLA}$ | -14.6038 | [-38.4871,8.592] |
| | | $\dot{x}^2_{CeA}$ | 0.7885 | [-2.1722,3.7472] | $\dot{x}^2_{BLA}$ | -1.3356 | [-3.9776,1.5136] |
| | | $x_{BLA}$ | -15.2149 | [-37.1866,5.9694] | $x_{CeA}$ | -1.4841 | [-40.5088,24.672] |
| | | $\dot{x}_{BLA}$ | 0.0947 | [-3.6129,3.9903] | $\dot{x}_{CeA}$ | 0.001 | [-4.5385,4.2258] |
| | | $x_{CeA}x_{BLA}$ | -5.9491 | [-30.4878,16.0943] | $x_{CeA}x_{BLA}$ | 24.4106 | [-13.1603,68.513] |
| | | $x_{BLA}\dot{x}_{CeA}$ | -2.7327 | [-20.0016,15.0187] | $x_{CeA}\dot{x}_{BLA}$ | -21.7178 | [-50.0637,8.2407] |
| | | $\dot{x}_{BLA}x_{CeA}$ | 1.877 | [-17.3515,20.5619] | $\dot{x}_{CeA}x_{BLA}$ | -3.6448 | [-25.272,22.9472] |
| | | $\dot{x}_{BLA}x_{BLA}$ | 0.3831 | [-2.6203,3.1663] | $\dot{x}_{CeA}x_{CeA}$ | 1.4549 | [-2.1212,5.1736] |
| Control | RR10 | $x_{CeA}$ | -48.038 | [-120.3989,-13.1295]* | $x_{BLA}$ | -40.6142 | [-87.0743,-6.6051]* |
| | | $\dot{x}_{CeA}$ | -28.9533 | [-33.9399,-23.3488]* | $\dot{x}_{BLA}$ | -28.6271 | [-34.6751,-22.4719]* |
| | | $x^2_{CeA}$ | 16.2441 | [-44.8327,99.8714] | $x^2_{BLA}$ | -26.2534 | [-109.9782,27.0757] |
| | | $x_{CeA}\dot{x}_{CeA}$ | 2.3718 | [-19.1513,25.0171] | $x_{BLA}\dot{x}_{BLA}$ | -6.2134 | [-25.8493,12.2774] |
| | | $\dot{x}^2_{CeA}$ | 0.1878 | [-2.943,2.3145] | $\dot{x}^2_{BLA}$ | -0.6939 | [-2.7079,1.6208] |
| | | $x_{BLA}$ | -11.6334 | [-50.57,16.6227] | $x_{CeA}$ | -12.1821 | [-66.3658,35.8943] |
| | | $\dot{x}_{BLA}$ | 0.5926 | [-3.6637,4.521] | $\dot{x}_{CeA}$ | -0.8044 | [-4.4138,2.7065] |
| | | $x_{CeA}x_{BLA}$ | 2.3742 | [-45.3075,52.3685] | $x_{CeA}x_{BLA}$ | 24.3214 | [-11.9912,78.993] |
| | | $x_{BLA}\dot{x}_{CeA}$ | 1.7543 | [-21.7826,26.0952] | $x_{CeA}\dot{x}_{BLA}$ | -8.1482 | [-31.215,16.0071] |
| | | $\dot{x}_{BLA}x_{CeA}$ | 7.298 | [-10.3076,24.6506] | $\dot{x}_{CeA}x_{BLA}$ | -8.2333 | [-34.1784,15.401] |
| | | $\dot{x}_{BLA}x_{BLA}$ | 0.9015 | [-1.2768,3.2041] | $\dot{x}_{CeA}x_{CeA}$ | 0.3097 | [-3.3213,3.9681] |
| Stress | FR1 | $x_{CeA}$ | -37.1336 | [-79.9109,-7.1022]* | $x_{BLA}$ | -1.2583 | [-19.8529,14.1595] |
| | | $\dot{x}_{CeA}$ | -30.8602 | [-41.7932,-20.2627]* | $\dot{x}_{BLA}$ | -28.2258 | [-34.7462,-21.7232]* |
| | | $x^2_{CeA}$ | 2.9784 | [-33.6353,39.4361] | $x^2_{BLA}$ | 23.0707 | [1.1936,52.4903]* |
| | | $x_{CeA}\dot{x}_{CeA}$ | 3.9179 | [-17.5091,28.5306] | $x_{BLA}\dot{x}_{BLA}$ | 1.2094 | [-11.6085,15.2722] |
| | | $\dot{x}^2_{CeA}$ | -0.0599 | [-2.4762,1.6447] | $\dot{x}^2_{BLA}$ | -1.8748 | [-5.5099,0.9211] |
| | | $x_{BLA}$ | -3.5931 | [-25.8749,33.3418] | $x_{CeA}$ | -1.206 | [-22.7559,20.9614] |
| | | $\dot{x}_{BLA}$ | -0.1721 | [-5.284,3.6893] | $\dot{x}_{CeA}$ | 2.3819 | [-3.8422,9.1137] |
| | | $x_{CeA}x_{BLA}$ | 19.681 | [-8.7993,68.2862] | $x_{CeA}x_{BLA}$ | 7.0132 | [-15.5718,28.5249] |
| | | $x_{BLA}\dot{x}_{CeA}$ | 3.5659 | [-32.7873,48.2716] | $x_{CeA}\dot{x}_{BLA}$ | 5.076 | [-8.0673,20.7058] |
| | | $\dot{x}_{BLA}x_{CeA}$ | -3.228 | [-22.0391,17.3801] | $\dot{x}_{CeA}x_{BLA}$ | 4.9383 | [-16.7006,29.8684] |
| | | $\dot{x}_{BLA}x_{BLA}$ | -0.1032 | [-3.8582,3.8374] | $\dot{x}_{CeA}x_{CeA}$ | 0.7971 | [-2.7171,4.063] |
| Stress | RR2 | $x_{CeA}$ | -26.5284 | [-79.1783,-4.6957]* | $x_{BLA}$ | -10.9022 | [-46.9009,10.3549] |
| | | $\dot{x}_{CeA}$ | -30.1741 | [-44.7242,-14.4774]* | $\dot{x}_{BLA}$ | -25.7279 | [-35.1094,-15.8241]* |
| | | $x^2_{CeA}$ | -25.0065 | [-87.1866,17.5633] | $x^2_{BLA}$ | 27.7792 | [9.446,54.424]* |
| | | $x_{CeA}\dot{x}_{CeA}$ | -2.5549 | [-26.0184,21.0414] | $x_{BLA}\dot{x}_{BLA}$ | 5.9384 | [-13.5194,24.8067] |
| | | $\dot{x}^2_{CeA}$ | -0.296 | [-3.1019,2.6101] | $\dot{x}^2_{BLA}$ | 0.8985 | [-1.8306,4.9814] |
| | | $x_{BLA}$ | -8.4024 | [-59.5639,28.1539] | $x_{CeA}$ | -18.0398 | [-43.6364,2.2489] |
| | | $\dot{x}_{BLA}$ | -0.2921 | [-5.0894,3.4705] | $\dot{x}_{CeA}$ | 0.692 | [-7.0322,9.2795] |
| | | $x_{CeA}x_{BLA}$ | 24.8449 | [-3.7387,72.8904] | $x_{CeA}x_{BLA}$ | -2.5064 | [-29.4409,21.8997] |
| | | $x_{BLA}\dot{x}_{CeA}$ | 0.85 | [-29.8768,32.4534] | $x_{CeA}\dot{x}_{BLA}$ | 3.6075 | [-14.0986,24.4418] |
| | | $\dot{x}_{BLA}x_{CeA}$ | -4.2753 | [-28.6923,22.2004] | $\dot{x}_{CeA}x_{BLA}$ | 1.6442 | [-19.4634,24.448] |
| | | $\dot{x}_{BLA}x_{BLA}$ | -0.1487 | [-4.8099,4.5271] | $\dot{x}_{CeA}x_{CeA}$ | 0.2129 | [-4.0232,4.2793] |
| Stress | RR5 | $x_{CeA}$ | -28.7232 | [-49.855,-10.6811]* | $x_{BLA}$ | -31.3253 | [-83.4871,10.4012] |
| | | $\dot{x}_{CeA}$ | -21.3577 | [-36.7323,-4.8758]* | $\dot{x}_{BLA}$ | -29.737 | [-39.0426,-18.5262]* |
| | | $x^2_{CeA}$ | -33.2245 | [-93.0387,10.5166] | $x^2_{BLA}$ | 30.3312 | [-1.9181,71.6459] |
| | | $x_{CeA}\dot{x}_{CeA}$ | -1.2989 | [-19.5916,19.1948] | $x_{BLA}\dot{x}_{BLA}$ | -0.1025 | [-23.9626,25.8226] |
| | | $\dot{x}^2_{CeA}$ | 0.8297 | [-1.4513,3.5435] | $\dot{x}^2_{BLA}$ | -0.6636 | [-3.3978,1.8092] |
| | | $x_{BLA}$ | -5.278 | [-47.776,30.6993] | $x_{CeA}$ | -0.1136 | [-27.1079,28.4284] |
| | | $\dot{x}_{BLA}$ | -0.1333 | [-4.0493,3.8989] | $\dot{x}_{CeA}$ | -1.2108 | [-15.2533,10.6527] |
| | | $x_{CeA}x_{BLA}$ | 12.6145 | [-12.1008,41.6946] | $x_{CeA}x_{BLA}$ | -5.1647 | [-34.4038,20.9884] |
| | | $x_{BLA}\dot{x}_{CeA}$ | 0.4611 | [-19.2789,17.7502] | $x_{CeA}\dot{x}_{BLA}$ | -4.3623 | [-38.8316,24.6348] |
| | | $\dot{x}_{BLA}x_{CeA}$ | 14.3963 | [-27.2196,52.4758] | $\dot{x}_{CeA}x_{BLA}$ | 1.4853 | [-17.8161,17.5684] |
| | | $\dot{x}_{BLA}x_{BLA}$ | 0.7657 | [-2.3299,4.4171] | $\dot{x}_{CeA}x_{CeA}$ | -0.212 | [-3.0066,3.4282] |
| Stress | RR10 | $x_{CeA}$ | -18.5431 | [-43.317,3.8353] | $x_{BLA}$ | -33.526 | [-82.5731,-1.7561]* |
| | | $\dot{x}_{CeA}$ | -25.5133 | [-35.6774,-14.8346]* | $\dot{x}_{BLA}$ | -27.5773 | [-35.3713,-19.6408]* |
| | | $x^2_{CeA}$ | -35.4398 | [-97.7617,16.7951] | $x^2_{BLA}$ | 22.6745 | [-12.3246,52.295] |
| | | $x_{CeA}\dot{x}_{CeA}$ | 0.7827 | [-16.3502,17.111] | $x_{BLA}\dot{x}_{BLA}$ | 3.4295 | [-25.315,27.5332] |
| | | $\dot{x}^2_{CeA}$ | 0.2008 | [-2.009,2.9737] | $\dot{x}^2_{BLA}$ | 0.2537 | [-2.777,2.9065] |
| | | $x_{BLA}$ | -0.4239 | [-50.4516,43.1699] | $x_{CeA}$ | -8.8894 | [-27.9183,10.891] |
| | | $\dot{x}_{BLA}$ | -0.3512 | [-5.1922,3.9351] | $\dot{x}_{CeA}$ | 0.0543 | [-7.9794,7.6854] |
| | | $x_{CeA}x_{BLA}$ | 18.9929 | [-16.5425,58.2538] | $x_{CeA}x_{BLA}$ | -11.1212 | [-46.9859,26.8651] |
| | | $x_{BLA}\dot{x}_{CeA}$ | 2.9045 | [-22.8722,25.7578] | $x_{CeA}\dot{x}_{BLA}$ | 0.3338 | [-25.4013,26.5216] |
| | | $\dot{x}_{BLA}x_{CeA}$ | 8.861 | [-24.2198,40.0348] | $\dot{x}_{CeA}x_{BLA}$ | -2.0835 | [-18.0292,15.0899] |
| | | $\dot{x}_{BLA}x_{BLA}$ | -1.2846 | [-5.4848,3.0091] | $\dot{x}_{CeA}x_{CeA}$ | 0.2588 | [-2.7118,4.1397] |

* means of statistical significance

Table D19: Population-level Dynamical Model Coefficients of Reward of Both Target Pathways

| | | BLA-DMS to CeA-DMS | | | CeA-DMS to BLA-DMS | | |
|---|---|---|---|---|---|---|---|
| Group | Session | Term | Mean | 95% CI | Term | Mean | 95% CI |
| Control | FR1 | $x_{CeA}$ | -63.7191 | [-132.5,-15.6442]* | $x_{BLA}$ | -31.2635 | [-79.8491,-2.2074]* |
| | | $\dot{x}_{CeA}$ | -29.2891 | [-35.7959,-19.4803]* | $\dot{x}_{BLA}$ | -25.571 | [-33.769,-14.9142]* |
| | | $x_{CeA}^2$ | 17.4216 | [-23.0267,53.5251] | $x_{BLA}^2$ | 37.5757 | [-17.0984,88.5443] |
| | | $x_{CeA}\dot{x}_{CeA}$ | 0.811 | [-28.2784,45.7123] | $x_{BLA}\dot{x}_{BLA}$ | 4.3809 | [-24.5301,24.2201] |
| | | $\dot{x}_{CeA}^2$ | -0.2234 | [-2.6302,2.6496] | $\dot{x}_{BLA}^2$ | 0.5576 | [-1.9472,2.9577] |
| | | $x_{BLA}$ | 0.3075 | [-28.9662,21.0143] | $x_{CeA}$ | -2.8931 | [-43.8313,35.0481] |
| | | $\dot{x}_{BLA}$ | 1.6408 | [-5.9636,8.317] | $\dot{x}_{CeA}$ | -0.6723 | [-4.7634,5.3278] |
| | | $x_{CeA}x_{BLA}$ | -1.0822 | [-39.0953,33.6319] | $x_{CeA}x_{BLA}$ | 10.455 | [-39.4599,62.1427] |
| | | $x_{BLA}\dot{x}_{CeA}$ | 3.5582 | [-32.3532,39.315] | $x_{CeA}\dot{x}_{BLA}$ | -0.9153 | [-25.992,20.7509] |
| | | $\dot{x}_{BLA}x_{CeA}$ | -7.0002 | [-23.2163,11.0721] | $\dot{x}_{CeA}x_{BLA}$ | 13.9446 | [-26.0481,55.0133] |
| | | $\dot{x}_{BLA}x_{BLA}$ | -0.2141 | [-3.8768,3.0199] | $\dot{x}_{CeA}x_{CeA}$ | 0.5191 | [-2.6141,4.0564] |
| Control | RR2 | $x_{CeA}$ | -74.8599 | [-170.67,-13.5072]* | $x_{BLA}$ | -58.8953 | [-146.6517,-17.889]* |
| | | $\dot{x}_{CeA}$ | -28.3124 | [-34.0463,-21.0298]* | $\dot{x}_{BLA}$ | -27.6345 | [-34.8747,-19.8039]* |
| | | $x_{CeA}^2$ | -19.9512 | [-118.975,38.152] | $x_{BLA}^2$ | 32.6839 | [-40.3802,109.3426] |
| | | $x_{CeA}\dot{x}_{CeA}$ | -0.9172 | [-29.8932,24.9326] | $x_{BLA}\dot{x}_{BLA}$ | 1.9411 | [-24.7584,25.0802] |
| | | $\dot{x}_{CeA}^2$ | -0.7027 | [-2.4226,1.5484] | $\dot{x}_{BLA}^2$ | -0.6947 | [-2.35,1.627] |
| | | $x_{BLA}$ | -16.4234 | [-78.2539,14.298] | $x_{CeA}$ | -5.143 | [-45.297,31.9394] |
| | | $\dot{x}_{BLA}$ | -1.5035 | [-7.7868,3.8177] | $\dot{x}_{CeA}$ | 1.0432 | [-4.3133,8.5542] |
| | | $x_{CeA}x_{BLA}$ | 41.7172 | [-16.3359,106.5755] | $x_{CeA}x_{BLA}$ | -37.1241 | [-98.338,12.5337] |
| | | $x_{BLA}\dot{x}_{CeA}$ | -3.0585 | [-36.1238,23.945] | $x_{CeA}\dot{x}_{BLA}$ | -3.3795 | [-24.6581,13.9261] |
| | | $\dot{x}_{BLA}x_{CeA}$ | 1.5288 | [-20.3167,18.5849] | $\dot{x}_{CeA}x_{BLA}$ | -4.9131 | [-31.7474,28.9534] |
| | | $\dot{x}_{BLA}x_{BLA}$ | 0.3716 | [-2.2099,3.4117] | $\dot{x}_{CeA}x_{CeA}$ | 0.541 | [-1.8937,3.2678] |
| Control | RR5 | $x_{CeA}$ | -35.1379 | [-97.1459,-0.3558]* | $x_{BLA}$ | -56.7473 | [-127.5276,-13.2471]* |
| | | $\dot{x}_{CeA}$ | -29.8535 | [-34.6174,-24.9233]* | $\dot{x}_{BLA}$ | -29.0495 | [-34.5744,-24.0736]* |
| | | $x_{CeA}^2$ | -10.1089 | [-90.9002,43.0576] | $x_{BLA}^2$ | 29.0594 | [-32.9625,103.6676] |
| | | $x_{CeA}\dot{x}_{CeA}$ | -6.5444 | [-26.2318,17.6448] | $x_{BLA}\dot{x}_{BLA}$ | 3.1445 | [-25.3218,32.8103] |
| | | $\dot{x}_{CeA}^2$ | 0.1101 | [-2.1137,2.0976] | $\dot{x}_{BLA}^2$ | -0.0923 | [-2.3872,2.563] |
| | | $x_{BLA}$ | -23.732 | [-68.2616,10.2806] | $x_{CeA}$ | -3.8426 | [-45.3105,36.6873] |
| | | $\dot{x}_{BLA}$ | -0.9811 | [-4.2596,2.6283] | $\dot{x}_{CeA}$ | 0.9511 | [-3.8911,7.4287] |
| | | $x_{CeA}x_{BLA}$ | 12.0859 | [-34.516,71.1657] | $x_{CeA}x_{BLA}$ | -37.9096 | [-103.7271,8.1616] |
| | | $x_{BLA}\dot{x}_{CeA}$ | 11.2882 | [-5.2026,34.1308] | $x_{CeA}\dot{x}_{BLA}$ | -14.2738 | [-46.028,14.5672] |
| | | $\dot{x}_{BLA}x_{CeA}$ | -1.1402 | [-25.6915,23.4554] | $\dot{x}_{CeA}x_{BLA}$ | -1.4312 | [-23.2558,20.8058] |
| | | $\dot{x}_{BLA}x_{BLA}$ | 0.4355 | [-2.4738,3.6345] | $\dot{x}_{CeA}x_{CeA}$ | -0.3211 | [-3.0567,2.6251] |
| Control | RR10 | $x_{CeA}$ | -53.9003 | [-152.989,0.9554] | $x_{BLA}$ | -66.1952 | [-168.9844,-12.4533]* |
| | | $\dot{x}_{CeA}$ | -30.3354 | [-35.9831,-23.4916]* | $\dot{x}_{BLA}$ | -28.32 | [-33.5599,-22.5294]* |
| | | $x_{CeA}^2$ | -16.9346 | [-129.611,66.8032] | $x_{BLA}^2$ | 37.2214 | [-41.8594,128.6759] |
| | | $x_{CeA}\dot{x}_{CeA}$ | 4.0001 | [-18.5473,28.5141] | $x_{BLA}\dot{x}_{BLA}$ | 6.5591 | [-13.334,20.5396] |
| | | $\dot{x}_{CeA}^2$ | -0.0506 | [-1.7297,1.44] | $\dot{x}_{BLA}^2$ | -0.2649 | [-2.497,1.4589] |
| | | $x_{BLA}$ | -21.5252 | [-77.5126,13.9993] | $x_{CeA}$ | -7.3366 | [-54.7169,40.9289] |
| | | $\dot{x}_{BLA}$ | -0.7598 | [-6.1106,4.0354] | $\dot{x}_{CeA}$ | 0.0543 | [-4.7121,4.9974] |
| | | $x_{CeA}x_{BLA}$ | 28.8916 | [-49.7226,152.4046] | $x_{CeA}x_{BLA}$ | -32.7652 | [-112.6417,30.0089] |
| | | $x_{BLA}\dot{x}_{CeA}$ | -6.6688 | [-32.6051,17.1618] | $x_{CeA}\dot{x}_{BLA}$ | 1.1047 | [-24.1307,20.462] |
| | | $\dot{x}_{BLA}x_{CeA}$ | 7.243 | [-11.255,37.4371] | $\dot{x}_{CeA}x_{BLA}$ | 1.9824 | [-19.0682,25.2618] |
| | | $\dot{x}_{BLA}x_{BLA}$ | -0.429 | [-3.0902,1.6389] | $\dot{x}_{CeA}x_{CeA}$ | 0.7995 | [-1.7455,3.5885] |
| Stress | FR1 | $x_{CeA}$ | -36.4426 | [-117.97,0.5687] | $x_{BLA}$ | -13.4218 | [-53.2329,17.7905] |
| | | $\dot{x}_{CeA}$ | -24.9794 | [-37.5587,-12.5023]* | $\dot{x}_{BLA}$ | -25.7574 | [-39.0702,-12.8613]* |
| | | $x_{CeA}^2$ | -36.8376 | [-112.971,14.1816] | $x_{BLA}^2$ | 22.8229 | [-2.2999,55.7604] |
| | | $x_{CeA}\dot{x}_{CeA}$ | -6.8896 | [-28.3923,14.4882] | $x_{BLA}\dot{x}_{BLA}$ | 4.4581 | [-17.3801,25.823] |
| | | $\dot{x}_{CeA}^2$ | 0.4684 | [-1.3553,2.6723] | $\dot{x}_{BLA}^2$ | 0.2841 | [-2.8146,3.8476] |
| | | $x_{BLA}$ | 5.3561 | [-21.2826,46.4225] | $x_{CeA}$ | 23.8481 | [-8.739,53.8513] |
| | | $\dot{x}_{BLA}$ | -0.7386 | [-4.9119,3.7283] | $\dot{x}_{CeA}$ | -3.508 | [-15.8219,9.418] |
| | | $x_{CeA}x_{BLA}$ | 17.7392 | [-28.676,71.6887] | $x_{CeA}x_{BLA}$ | 0.8356 | [-31.6723,33.7384] |
| | | $x_{BLA}\dot{x}_{CeA}$ | -11.2347 | [-53.0909,15.9854] | $x_{CeA}\dot{x}_{BLA}$ | -8.5715 | [-31.6585,12.4547] |
| | | $\dot{x}_{BLA}x_{CeA}$ | 6.9279 | [-13.4039,27.2444] | $\dot{x}_{CeA}x_{BLA}$ | -6.8175 | [-28.0207,15.2696] |
| | | $\dot{x}_{BLA}x_{BLA}$ | -2.2747 | [-5.9343,1.4817] | $\dot{x}_{CeA}x_{CeA}$ | -0.9981 | [-4.53,1.6066] |
| Stress | RR2 | $x_{CeA}$ | -33.9737 | [-81.4017,-6.1657]* | $x_{BLA}$ | -20.8664 | [-57.2732,6.8924] |
| | | $\dot{x}_{CeA}$ | -26.9646 | [-39.0794,-12.8915]* | $\dot{x}_{BLA}$ | -29.0254 | [-44.1673,-13.0756]* |
| | | $x_{CeA}^2$ | -18.7255 | [-78.77,22.3293] | $x_{BLA}^2$ | 21.2358 | [-11.0838,52.9956] |
| | | $x_{CeA}\dot{x}_{CeA}$ | -3.5388 | [-23.9721,20.4668] | $x_{BLA}\dot{x}_{BLA}$ | 0.3114 | [-25.0889,29.1353] |
| | | $\dot{x}_{CeA}^2$ | 0.2659 | [-1.8775,2.9907] | $\dot{x}_{BLA}^2$ | 0.5782 | [-2.1561,3.4222] |
| | | $x_{BLA}$ | 9.0799 | [-22.2961,43.8437] | $x_{CeA}$ | 11.3185 | [-22.2613,42.0485] |
| | | $\dot{x}_{BLA}$ | -1.5429 | [-9.2769,1.9996] | $\dot{x}_{CeA}$ | 5.0521 | [-7.2591,20.1773] |
| | | $x_{CeA}x_{BLA}$ | 19.2759 | [-9.279,58.8671] | $x_{CeA}x_{BLA}$ | -11.5931 | [-53.1596,17.3722] |
| | | $x_{BLA}\dot{x}_{CeA}$ | 10.725 | [-18.527,51.8099] | $x_{CeA}\dot{x}_{BLA}$ | 9.0752 | [-14.0992,36.57] |
| | | $\dot{x}_{BLA}x_{CeA}$ | 2.7608 | [-18.3218,26.5381] | $\dot{x}_{CeA}x_{BLA}$ | -0.8977 | [-25.8329,24.7231] |
| | | $\dot{x}_{BLA}x_{BLA}$ | -0.0139 | [-3.2493,3.5607] | $\dot{x}_{CeA}x_{CeA}$ | 0.2955 | [-3.3654,4.0565] |
| Stress | RR5 | $x_{CeA}$ | -24.1733 | [-72.5317,6.7114] | $x_{BLA}$ | -25.7686 | [-83.3483,12.8304] |
| | | $\dot{x}_{CeA}$ | -30.2505 | [-43.7619,-16.1942]* | $\dot{x}_{BLA}$ | -25.8617 | [-38.6042,-13.7752]* |
| | | $x_{CeA}^2$ | -21.4512 | [-66.5301,27.774] | $x_{BLA}^2$ | 26.8718 | [-33.0602,78.2952] |
| | | $x_{CeA}\dot{x}_{CeA}$ | 1.7399 | [-16.1616,30.1938] | $x_{BLA}\dot{x}_{BLA}$ | 0.3074 | [-27.7318,25.5828] |
| | | $\dot{x}_{CeA}^2$ | 0.5146 | [-1.4876,2.4932] | $\dot{x}_{BLA}^2$ | 0.5848 | [-2.152,4.4049] |
| | | $x_{BLA}$ | -9.856 | [-42.9484,18.3157] | $x_{CeA}$ | -1.2114 | [-38.8271,38.3427] |
| | | $\dot{x}_{BLA}$ | 0.5822 | [-6.3364,9.1395] | $\dot{x}_{CeA}$ | 2.0352 | [-8.4784,15.4467] |
| | | $x_{CeA}x_{BLA}$ | 6.8592 | [-23.6643,41.3629] | $x_{CeA}x_{BLA}$ | -18.0266 | [-72.0765,24.0544] |
| | | $x_{BLA}\dot{x}_{CeA}$ | 0.393 | [-28.867,31.5416] | $x_{CeA}\dot{x}_{BLA}$ | 3.9668 | [-21.7191,34.195] |
| | | $\dot{x}_{BLA}x_{CeA}$ | -0.6885 | [-26.4349,26.4129] | $\dot{x}_{CeA}x_{BLA}$ | -8.6574 | [-31.469,10.7028] |
| | | $\dot{x}_{BLA}x_{BLA}$ | -1.1131 | [-4.5578,1.9544] | $\dot{x}_{CeA}x_{CeA}$ | 0.0917 | [-3.3289,3.4834] |
| Stress | RR10 | $x_{CeA}$ | -41.0836 | [-108.046,14.6808] | $x_{BLA}$ | -28.7362 | [-88.2665,3.4026] |
| | | $\dot{x}_{CeA}$ | -28.0479 | [-43.3901,-15.4437]* | $\dot{x}_{BLA}$ | -24.6554 | [-37.9969,-12.2315]* |
| | | $x_{CeA}^2$ | -42.0699 | [-109.615,50.1911] | $x_{BLA}^2$ | 22.4898 | [-33.3971,75.2735] |
| | | $x_{CeA}\dot{x}_{CeA}$ | 0.1707 | [-25.2704,22.4619] | $x_{BLA}\dot{x}_{BLA}$ | 4.325 | [-21.9373,28.1337] |
| | | $\dot{x}_{CeA}^2$ | -0.3584 | [-2.0014,1.7979] | $\dot{x}_{BLA}^2$ | -0.1345 | [-2.5135,2.4612] |
| | | $x_{BLA}$ | -26.5499 | [-92.5534,31.2197] | $x_{CeA}$ | -7.6708 | [-46.0614,24.7455] |
| | | $\dot{x}_{BLA}$ | 2.1102 | [-17.5767,20.993] | $\dot{x}_{CeA}$ | 2.1248 | [-11.4362,13.3194] |
| | | $x_{CeA}x_{BLA}$ | -0.9876 | [-54.7945,54.4127] | $x_{CeA}x_{BLA}$ | -14.2875 | [-71.0832,15.243] |
| | | $x_{BLA}\dot{x}_{CeA}$ | -13.431 | [-62.4875,34.2638] | $x_{CeA}\dot{x}_{BLA}$ | 2.3578 | [-29.8339,30.2362] |
| | | $\dot{x}_{BLA}x_{CeA}$ | 3.2964 | [-27.5415,34.3215] | $\dot{x}_{CeA}x_{BLA}$ | -4.4279 | [-29.9438,18.7915] |
| | | $\dot{x}_{BLA}x_{BLA}$ | -0.6505 | [-4.7874,2.9784] | $\dot{x}_{CeA}x_{CeA}$ | -1.4075 | [-4.501,1.5081] |

* means of statistical significance

Table D20: Robustness of Mean Cumulative KL Divergence across Bandwidth Perturbations

| Bandwidth Factor | Group | Mean Cumulative $D_{KL}$ | 95% CI |
|---|---|---|---|
| 0.9 (Narrow) | Control | 30133.0249 | (7094.8575, 86741.2723) |
| 0.9 (Narrow) | Stress | 53832.4717 | (25992.5547, 84713.6858) |
| 1.0 (Optimal) | Control | 26487.6454 | (5842.6635, 74563.2858) |
| 1.0 (Optimal) | Stress | 50378.8123 | (23324.2554, 78289.6453) |
| 1.1 (Wide) | Control | 23895.2978 | (7945.3240, 63032.1457) |
| 1.1 (Wide) | Stress | 50159.8250 | (22653.1008, 81956.6318) |
